# High retention of genomic variation and fitness-related traits in the effective population of reintroduced wolves in Yellowstone National Park

**DOI:** 10.1101/2022.02.18.481090

**Authors:** Bridgett M. vonHoldt, Alexandra L. DeCandia, Kira A. Cassidy, Erin E. Stahler, Janet S. Sinsheimer, Douglas W. Smith, Daniel R. Stahler

## Abstract

Cooperatively breeding species exhibit numerous strategies to avoid mating with close relatives, inherently reducing effective population size. For species of management concern, accurate estimates of inbreeding and trait depression are crucial for the species’ future. We utilized genomic and pedigree data for Yellowstone National Park gray wolves to investigate the contributions of foundation stock lineages, genetic architecture of the effective population, and putative fitness consequences of inbreeding. Our dataset spans 25 years and seven generations since reintroduction, encompassing 152 nuclear families and 329 litters. We found over 87% of the pedigree foundation genomes persisted and report influxes of allelic diversity from two translocated wolves from a divergent source in Montana. As expected for group-living species, mean kinship significantly increased over time, although we found high retention of genetic variation. Strikingly, the effective population carried a significantly lower level of genome-wide inbreeding coefficients and autozygosity with shorter decays for linkage disequilibrium relative to the non-breeding population. Lifespan and heterozygosity were higher in the effective population, although individuals who had their first litter at an older age also had higher inbreeding coefficients. Our findings highlight genetic contributions to fitness, and the importance of effective population size and gene flow to counteract loss of genetic variation in a wild, free-ranging social carnivore. It is crucial for managers to mitigate factors that significantly reduce effective population size and genetic connectivity, which supports the dispersion of genetic variation that aids in rapid evolutionary responses to environmental challenges.

## Introduction

Territorial, cooperatively-breeding species can exhibit a diversity of breeding strategies (e.g. monogamy, polygamy, polyandry) and reproductive skew (Keller and Reeve 1994; Clutton-Brock 2016). Yet regardless of these patterns, access to reproduction is restricted and effective population size (N_e_) is reduced (Jennions and Macdonald 1994; Frankham 1995; Komdeur and Deerenberg 1997). Estimates of N_e_ provide critical information during the application and interpretation of evolutionary and population genetic theory (Wright 1931, 1969; Crow and Kimura 1970; Lanfear et al. 2014), as well as for wildlife management programs (Rowe and Beebee 2004). Species that have experienced either a natural population decline or bottleneck are expected to have a dramatically reduced census and effective size, and an increased probability of inbreeding and genetic identity by descent or autozygosity (Hedrick and Kalinowski 2000; Charlesworth and Willis 2009). Consequently, inbreeding can dramatically impact N_e_ estimates through increased genetic correlations and decreased frequency of heterozygotes (Ellegren 1999; Wang et al. 2016). Viability is then threatened by inbreeding or trait depression, the negative fitness consequence expected in inbred offspring (Lynch and Walsh 1998; Charlesworth and Willis 2009; Hedrick and Garcia-Dorado 2016).

Inbreeding depression is often measured in fitness-related traits of life history (e.g. fecundity, survival, morphological measurements) and often only in captive populations (Hedrick and Kalinowski 2000). Measuring autozygosity further provides an assessment of the full effect of deleterious alleles, as individuals that carry greater levels of autozygosity are expected to exhibit decreased fitness (Charlesworth and Willis 2009).

Although there has been much attention paid to inbreeding avoidance and trait depression in captive breeding programs from pedigree estimates (Ralls et al. 1988; Laikre and Ryman 1991; Laikre et al. 1993; Kalinowski et al. 2000; Kalinowski and Hedrick 2001; Jiménez-Mena et al. 2016), much less is known about the genetic basis of depression in wild populations, although it is expected to exist (Lacy 1997; Keller and Waller 2002; Curik et al. 2017).

Further, pedigrees that are unbalanced, incorrect, or incomplete can lead to errors in estimating inbreeding coefficients and trait depression (Curik et al. 2017). One critical aspect of pedigree-based errors is the assumption that pedigree founders are unrelated. Genome data now provide an opportunity to directly and accurately measure autozygosity, which quantifies the fraction of the genome that is contained within long stretches of homozygosity delineated along chromosomes (*F_ROH_*; McQuillan et al. 2008). Keller et al. (2011) also showed that *F_ROH_* outperformed inbreeding estimates derived from pedigree data.

We studied the dynamic landscape of inbreeding and life-history traits in a pedigreed population of gray wolves (*Canis lupus*). As a cooperatively- breeding species, wolf populations are subdivided with limited access to reproduction that is reinforced by a social hierarchy and social rank-related stress reflected in glucocorticoid levels (Sands and Creel 2004; vonHoldt et al. 2008). Several studies of small, isolated, bottlenecked or captive gray wolf populations across their Holarctic range have found genetic evidence of high inbreeding levels with a corresponding reduction in fitness and related population genetic health metrics. Two primary examples of such isolated island populations of wolves are those inhabiting Isle Royale in Lake Superior and Scandinavia (Laikre and Ryman 1991, 1993; Ellegren 1999; Flagstad et al. 2003; Liberg et al. 2005; Räikkönen et al. 2009, 2013; Adams et al. 2011; Hedrick et al. 2014; Åkesson et al. 2016, 2021; Robinson et al. 2019). Specific to the wolves inhabiting the contiguous United States, effective government wildlife control programs eradicated nearly all gray wolves by the early 1930s (except in Minnesota; Fritts et al. 1997). Over five decades later, substantial support developed to establish a reintroduction plan for gray wolf restoration to the Rocky Mountains (Fritts et al. 1997). During the winters of 1995 and 1996, the US Fish and Wildlife Service released 35 wolves in central Idaho, and in a joint effort with the National Park Service, released 31 wolves in Yellowstone National Park (YNP). Additionally, there was the translocation of 10 wolf pups from the Sawtooth pack in northwestern Montana to YNP where they were released in 1997. Foundation stock (a.k.a. YNP founders) were captured from two source locations in western Canada, and vonHoldt et al. (2008) utilized 26 microsatellites to provide valuable genetic insights with respect to their relatedness and population genetic health during the first decade of recovery. It was reported that YNP founder breeding pairs were unrelated with near absolute inbreeding avoidance despite living in groups of related individuals. Yet, given the resolution limits of microsatellite data for investigating genomic architecture, little is known about the nature of genetic variation or chromosomal regions that are identical by descent (IBD).

Building upon the recent release of their updated pedigree constructed using genome-wide SNP data (vonHoldt et al. 2020), we explore here the genetics of YNP founders’ lineages, demography, genomic contributions, inbreeding, and consequences thereof in YNP gray wolves during their 25 years post-reintroduction. We assessed pedigree- and marker-based aspects of the wolf population with the unique opportunity to integrate life-history fitness correlates with respect to reproduction. With the pedigree reconstructed utilizing genomic data, these two perspectives have provided new details regarding the genetic viability, reproduction, and recovery status of this social carnivore.

## Materials and Methods

### Yellowstone National Park’s reintroduced wolf population

We analyzed 474 YNP gray wolves with pedigree and 391 with genome-wide SNP data derived from blood and tissue samples collected between 1995 and 2020 (see below). After reintroduction, the YNP population rapidly expanded, reaching a high of 174 wolves in 2003. However, the population stabilized over the last decade (Smith et al. 2020), averaging 97.9 (±3.9 SE) at the year-end official census count. For this study, the original 31 wolves translocated to YNP and the parents of the 10 translocated orphaned pups were designated as *YNP founders*. This is not to be confused with the animals at the foundation of the pedigree, known as the *pedigree founders*, which included two wolves sampled in Canada who were not translocated but known to be related to YNP wolves. The original and updated genealogy for this population confirmed close kinships (e.g. parent/offspring of full-siblings) and genetic similarities among a subset of the YNP founders (vonHoldt et al. 2008, 2020). These kinship ties were expected given the nature of the wild capture methods and release. Several individuals were captured together, likely representing a pack or a subset thereof. These relationships are reflected by increased allele sharing in the genetic-based analysis below, while the pedigree-based analysis incorporated these known relationships. These relationships are critical to account for as most treatments of founders assume they are unrelated (i.e. their kinship *f_xy_* and inbreeding coefficients *F* are set to 0; Jiménez-Mena et al. 2016).

### Pedigree-based analysis

We updated the full YNP wolf pedigree by adding three new individuals indicated in Supplemental Table S1 and used the same genomic preparation and analytical methods described by vonHoldt et al. (2020).

Briefly, we constructed a parentage-informative dataset of 736 SNPs through strict filtering in *PLINK v1.9* (Chang et al. 2015) with parameter settings recommended for pedigree reconstruction (Huisman 2017): Hardy-Weinberg equilibrium (HWE) (--hwe 0.001), minor allele frequency (MAF) (--maf 0.45), and statistical linkage disequilibrium (LD) (--indep-pairwise 50 5 0.2). We calculated pairwise relatedness coefficients (*r*) using the *coancestry* function within the R v3.6.0 (2019) package *related* and implemented the dyadic likelihood estimator (dyadml=1; Milligan 2003) with allowance for inbreeding (allow.inbreeding=TRUE) (Pew et al. 2015). We then used these estimates alongside observation data and likely parent-offspring (P-O) pairs assigned by the R v3.6.0 package *sequoia* (Huisman 2017) to update the full YNP wolf pedigree to include 871 P-O pairs among 474 wolves (n, females=226, males=239, unknown=9; Table S1) (vonHoldt et al. 2020). Wolves that lacked genetically-confirmed parents or offspring were excluded from the pedigree analysis. For individuals where the likely birthday was a range of years, we assigned it the earliest birth year. We completed several of the described methods for the full pedigreed population, as well as annually between 1995 and 2020.

We used *PedScope©* v.2.4.01, a proprietary program used for animal population management and breeding recommendations, to perform pedigree- based analyses. The pedigree contained 26 pedigree founders, defined as individuals where no parents are included in the pedigree (Table S1).

Correlations were assessed using the product moment correlation coefficient (r). We estimated several measures of gene diversity in the pedigreed population. We estimated the number of founder equivalents (*fe*) and founder genome equivalents (*fg*). The former measured gene diversity in a specified set of the pedigree (*e.g.* pedigree founders, annual, or entire) and represents the number of equally contributing founders expected to produce the observed level of genetic diversity in the specified population. The latter is a related metric that accounts for drift, where *fg* represents an estimate that incorporates random loss of alleles. Further, we estimated the number of effective ancestors (*fa*) and identified influential ancestors as those with the greatest contributions towards *fa* using an algorithm developed by Boichard et al. (1997). Estimates of *fa* are similar to that of *fe* but account for bottlenecks in the pedigree.

We surveyed genome uniqueness (*GU*) defined as the animal’s likelihood that it carries pedigree founder alleles not present in another individual in the pedigree. We also estimated mean kinship (*MK*), Wright’s pedigree-based inbreeding coefficient (*F*), and Ballou’s direct method for estimating ancestral inbreeding coefficient. In a pedigree-based approach, pairwise kinship values describe the probability that two alleles shared are IBD, a measure also estimated by the pedigreed population’s inbreeding coefficient F. We obtained the pairwise additive genetic relationship matrix (“A” matrix) to also assess pairwise relatedness. Differences in pedigree metrics were assessed using a 1-way ANOVA in R v3.6.0.

We counted the number of litters per individual as the total number of years a specific mating pair reproduced and included only offspring where birth year was known. We also included litter counts if only one parent was known. We used the log_10_ of the number of genetic litters as a proxy for fitness, surveyed as a function of inbreeding coefficients, where the slope of the fitted line is inferred as the inbreeding load (*-B*) (Keller and Waller 2002). We further used the pedigree to estimate the total number of lethal equivalents (LE) per diploid genome (*2B*). The inbreeding load is often defined as the reduction in survival expected in a completely homozygous individual. We then estimated the expected depression in fitness (δ) following δ=1-e^-*B*\**F*^, where *B* is the estimated number of lethal equivalents per gamete and *F* is the inbreeding coefficient.

### Genome-wide SNP genotype data

Although the pedigree relationships are valuable, we utilized a marker- based genomics approach to explore the impact of pedigree structure on genomic variation and life-history related traits (Kardos et al. 2015). A subset of wolves in the pedigree were excluded from the genetic analyses due to low sequence coverage or lack of a genetic sample (*i.e.* only observation data supported their pedigree information). As such, the pedigree contains more individuals than analyzed with RADseq data. Further, we included observational data regarding parentage and reproduction. Such an exception was made when reproduction was observed but a genetic sample was unavailable. We analyzed 56 576 SNP loci (referred to as 56K) genotyped across 391 gray wolves (n, females=193, males=195, unknown=3), which were previously collected using a modified restriction-site associated DNA sequencing (RADseq) protocol by Ali et al. (2016) and mapped to the reference dog genome Canfam3.1 (vonHoldt et al. 2020). These public data were previously filtered to exclude individuals with >20% missing data and sites with >10% per-site missing data, MAF<1%, or significant deviation from HWE (*p*<0.0001), leading to a final genotyping rate of 96.4%. We estimated LD decay using the --r2 flag across 391 YNP wolves genotyped for 56K SNPs. We further removed SNPs with a genotype correlation of r^2^>0.5 using the -- indep-pairwise 50 5 0.5 flag to construct a pruned set of 24,235 statistically unlinked SNPs (referred to as 24K). Individual wolves with confirmed offspring via genetics or observation were categorized as “breeding” individuals (n=152), while wolves lacking offspring were considered “non- breeding” individuals (n=235), with four wolves of unknown breeding status (Table S2). We also completed several of the described methods for the full pedigreed population overall as well as annually and censored, between 1995 and 2018, with the pruned SNP set. Differences in annual estimates were assessed using a 1-way ANOVA in R. We estimated standard observed and expected heterozygosity (H_O_ and H_E_, respectively) in *PLINK* with the --hardy and --het flags.

We also obtained the predicted impact and consequence of each allele from *Variant Effect Predictor* (VEP) (McLaren et al. 2016). Based on *VEP* impact annotations, we grouped the 17 loci with “*high*” predicted impact and 381 loci with “*moderate*” impact into a category (n=398 loci) inferred to negatively impact the individual fitness. This category was further detailed as the “*high*” impact loci were annotated as the gain/loss of a stop, loss of a start, or a splice acceptor/donor, while loci with “*moderate*” impact were missense variants and splice regions. We also grouped 844 loci annotated as “*low*” impact and 29 213 loci with a “*modifier*” impact in a category inferred to unlikely impact fitness (n=30 057 loci).

### Estimating the inbreeding coefficient from genotype data

We estimated individual inbreeding coefficients (F) from the pruned SNP set and runs of homozygosity (ROH) with the R v3.6.0 function *detectRUNS* (Biscarini et al. 2018) as:

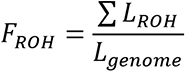

where *L_ROH_* is the summed length of all ROHs detected in an individual and *L_genome_* is the length of the genome that is used. As RADseq data are inherently clustered as sequence stacks across the genome at restriction enzyme cut sites, we used the consecutive window-free option for detecting homozygous and heterozygous tracks (Marras et al. 2015). Tracks were detected by a minimum of 10 SNPs in a track of at least 10 000 nucleotides with a maximum gap of 10^6^ nucleotides between SNPs, and allowed for a maximum of a single opposite or missing genotype in the track. We further annotated each ROH with respect to their composition of alleles with either a functional impact or consequence (see VEP methods above) using the *intersectBed* function of *Bedtools* v2.28.0 (Quinlan and Hall 2010).

### Modeling reproduction, life history, and genetic parameters

We examined the age at first litter (*Age_first_litter*) using Cox proportional hazards regression for survival (*survival* version 3.2-11) with time (in years) to the earliest of three events first litter, death or last documented observation. Animals who died or were lost to observation before breeding were considered right censored data in the analysis. We used the exact method to deal with ties when two or more animals had the same survival times. We standardized the potential predictors (observed heterozygosity, *F_ROH_* and VEP consequence categories), and included sex and the first five principle components (PCs) as covariates in all analyses. We fit the following model structure:

### Fit=coxph(Surv(Age_litter,Censoring)∼Predictor,ties=“exact”,data)

For the Cox proportional hazards regression analysis, although *p*-values are reported, we elected the best fit models through assessment of the Akaike information criterion (AIC).

## Results

### Pedigree structure

Using *Pedscope©*, the pedigree composed of 26 founders of the pedigree (individuals where no parents are found within the pedigree), 152 nuclear families (individuals that share the same parents), and 319 litters. Compared to the census size of 41 YNP founder individuals, we inferred 19.92 pedigree founder equivalents (*fe*), 18.53 founder genome equivalents (*fg*), and 18.79 effective ancestors (*fa*) in the pedigree analysis. There was a maximum ancestry depth of seven generations (Average±s.d.=3.8±1.6), with an average kinship (MK) of 3.2%. We restricted the founder analysis to the wolves translocated in years 1995 and 1996, which included the individuals translocated from the Sawtooth pack in northwestern Montana. We found that wolf 009F had the highest contribution as measured by the percent contribution in the current pedigree due to that individual (*pC*=12%; average=3.8%), followed by 005F with 9% to the full pedigree of 474 wolves, and then by their respective mates 010M and 004M both with 8% contributions each (Table 1).

**Table 1.** Estimates of the pedigree founder contributions and genomic uniqueness in Yellowstone gray wolves. Genome uniqueness (*GU*) is the probability that a gene from the pedigree foundation was inherited ‘uniquely’ with respect to other pedigree founders. (Abbreviations: *pAR*, proportion of alleles retained; *pC*, proportion of genes in the pedigreed population due to that founder; YNP, Yellowstone National Park)

| Founder ID | Source Pack (age class and sex)* | YNP Pack | *pC* | *pAR* | *GU* |
| --- | --- | --- | --- | --- | --- |
| 004M | Petite Lake (adult male) | Mollies | 0.08 | 0.94 | 0.00 |
| 005F | Petite Lake (adult female) | Mollies | 0.09 | 1.00 | 0.00 |
| 009F | McLeod (adult female) | Rose Creek | 0.12 | 1.00 | 0.00 |
| 010M | Rick’s (adult male) | Rose Creek | 0.08 | 0.99 | 0.00 |
| 013M | Berland (adult male) | Yellowstone Delta | 0.02 | 0.98 | 0.01 |
| 014F | Berland (adult female) | Yellowstone Delta | 0.02 | 0.99 | 0.00 |
| 015M | Berland (male pup) | Yellowstone Delta | 0.00 | 0.50 | 0.33 |
| 027F | Halfway (adult female) | Nez Perce | 0.04 | 0.99 | 0.00 |
| 028M | Halfway (adult male) | Nez Perce | 0.04 | 0.99 | 0.00 |
| 035M | Chief (adult male) | Lone Star | 0.00 | 0.88 | 0.25 |
| 036F | Besa (adult female) | Lone Star | 0.00 | 0.50 | 1.00 |
| 038M | Prophet (adult male) | Druid Peak | 0.02 | 0.99 | 0.00 |
| 039F | Besa (adult female) | Druid Peak | 0.03 | 0.88 | 0.00 |
| Y38 | Unknown (adult male) | *** | 0.02 | 0.50 | 0.00 |
| Y53 | Kravec (adult female) | *** | 0.01 | 0.87 | 0.06 |
| Sawtooth F** | Sawtooth (adult female) | Nez Perce | 0.04 | 1.00 | 0.00 |
| Sawtooth M** | Sawtooth (adult male) | Nez Perce | 0.04 | 1.00 | 0.00 |

\* The name of the pack in Canada from which the founder was captured.
\*\* These two entries represent the known (and unsampled) parents of 10 full-sibling pups translocated (and genetically sampled) from the Sawtooth pack in northwestern Montana and fostered by the Nez Perce pack. Two of the translocated Sawtooth individuals assumed a breeding status in Yellowstone and represent gene flow (outbreeding) events. \*\*\* Offspring of these individuals were translocated to the Chief Joseph pack in YNP.

Nearly all pedigree founders had ≥87% of their alleles retained in the total pedigree (Average±s.d.=88%±20%), with only three individuals at 50% retention (015M, 036F, and a Canada source wolf Y38 whose offspring were translocated to the Chief Joseph pack in YNP) (Table 1).

Average levels of founder genome uniqueness were low (*GU*=0.10±0.3), with only a single pedigree founder (036F) displaying absolute uniqueness (*GU*=1), and four additional individuals with non-zero values (013M, 015M, 035M, Y53) indicating the likelihoods that alleles are not present in another individual in the pedigree (Table 1). A survey of the total pedigree also revealed 20 influential pedigree founders, similarly highlighting the largest marginal contributions by mated pair 009F (12%) and 010M (8%), and mated pair 004M and 005F (8% and 9%, respectively) (Table 2).

**Table 2.** Influential ancestors identified by their marginal contribution (MC) to the total pedigree of YNP wolves. (Abbreviations: F, female; M, male)

| Field ID | Source Pack (age class) | YNP pack | MC |
| --- | --- | --- | --- |
| 004M | Petite Lake (dominant) | Mollies | 0.08 |
| 005F | Petite Lake (dominant) | Mollies | 0.09 |
| 006M | Petite Lake (pup) | Mollies | 0.02 |
| 007F | Mcleod (pup) | Rose Creek | 0.03 |
| 009F | Mcleod (dominant) | Rose Creek | 0.12 |
| 010M | Rick's (dominant) | Rose Creek | 0.08 |
| 013M | Berland (dominant) | Yellowstone Delta | 0.01 |
| 014F | Berland (dominant) | Yellowstone Delta | 0.01 |
| 027F | Halfway (dominant) | Nez Perce | 0.01 |
| 028M | Halfway (dominant) | Nez Perce | 0.01 |
| 034M | Unknown (dominant) | Chief Joseph | 0.04 |
| 038M | Prophet (dominant) | Druid Peak | 0.02 |
| 039F | Besa (dominant) | Druid Peak | 0.03 |
| 040F | Besa pack (pup) | Druid Peak | 0.01 |
| 042F | Besa pack (pup) | Druid Peak | 0.01 |
| 048F* | - | Nez Perce | 0.07 |
| 072M | Sawtooth pack (pup) | Nez Perce | 0.07 |
| Y53 | Kravic (female) | *** | 0.01 |
| Sawtooth F** | Sawtooth (female) | Nez Perce | 0.01 |
| Sawtooth M** | Sawtooth (male) | Nez Perce | 0.01 |

\* The first generation of offspring born in YNP's Nez Perce pack by founders 027F and 028M.
\*\* These two entries represent the known (and unsampled) parents of 10 full-sibling pups translocated (and genetically sampled) from the Sawtooth pack in northwestern Montana and fostered by the Nez Perce pack. Two of the translocated Sawtooth individuals assumed a breeding status in Yellowstone and represent gene flow (outbreeding) events.
\*\*\* Offspring of these individuals were translocated to the Chief Joseph pack in YNP.

We found a significant increase in annual MK with decreasing size of the genotyped population analyzed (F_(1,22)_=14.6, *p*=0.0009), an increase of genomic uniqueness over time (F=9.6, *p*=0.0053), an increase in founder equivalents (F=38.4, *p*=3.063×10^-6^), and a decrease in the number of founder genome equivalents over time (F=15.8, *p*=0.0006) (Table 3). The number of effective founders did not significantly change over time (F=0.7, *p*=0.395) (Table 3).

**Table 3.**
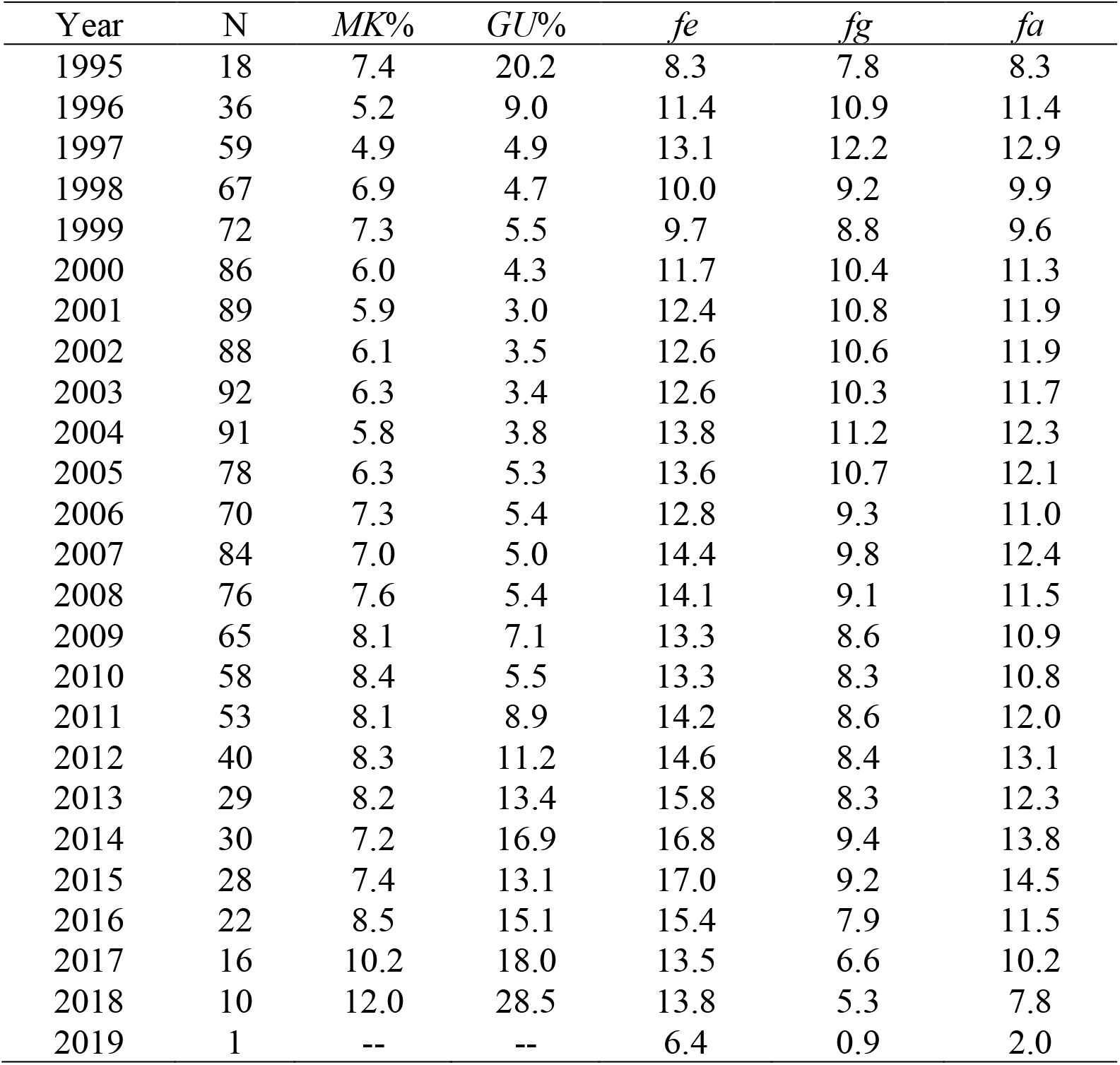
Annual metrics of the pedigreed gray wolf population in Yellowstone National Park with known birth and death dates. Genomic uniqueness (*GU*) is an intra-annual population metric. (Abbreviations: *fa*, number of effective founders; *fe*, number of found equivalents; fg, number of founder genome equivalents; *MK*, mean kinship; N, annual sample size analyzed)

When we excluded observations of 0 litters, 153 observations remained with low levels of inbreeding coefficients (average *F*=0.009±0.03; range=0.0- 0.25) and was negatively correlated to the number of litters (Pearson r=-0.076, *p*=0.35) (Fig. 1A). From this, we estimated the inbreeding load as the slope of the trendline between log_10_ litter number and inbreeding coefficient (*y*=- 0.595*x*+0.292, *-B*=0.595), and thus *2B*=0.595 lethal equivalents (LE) in a diploid genome. Following, we estimated fitness (δ) is expected to be depressed in 14% of progeny from a mating of first-degree relatives (*F*=0.25).

**Figure 1.**
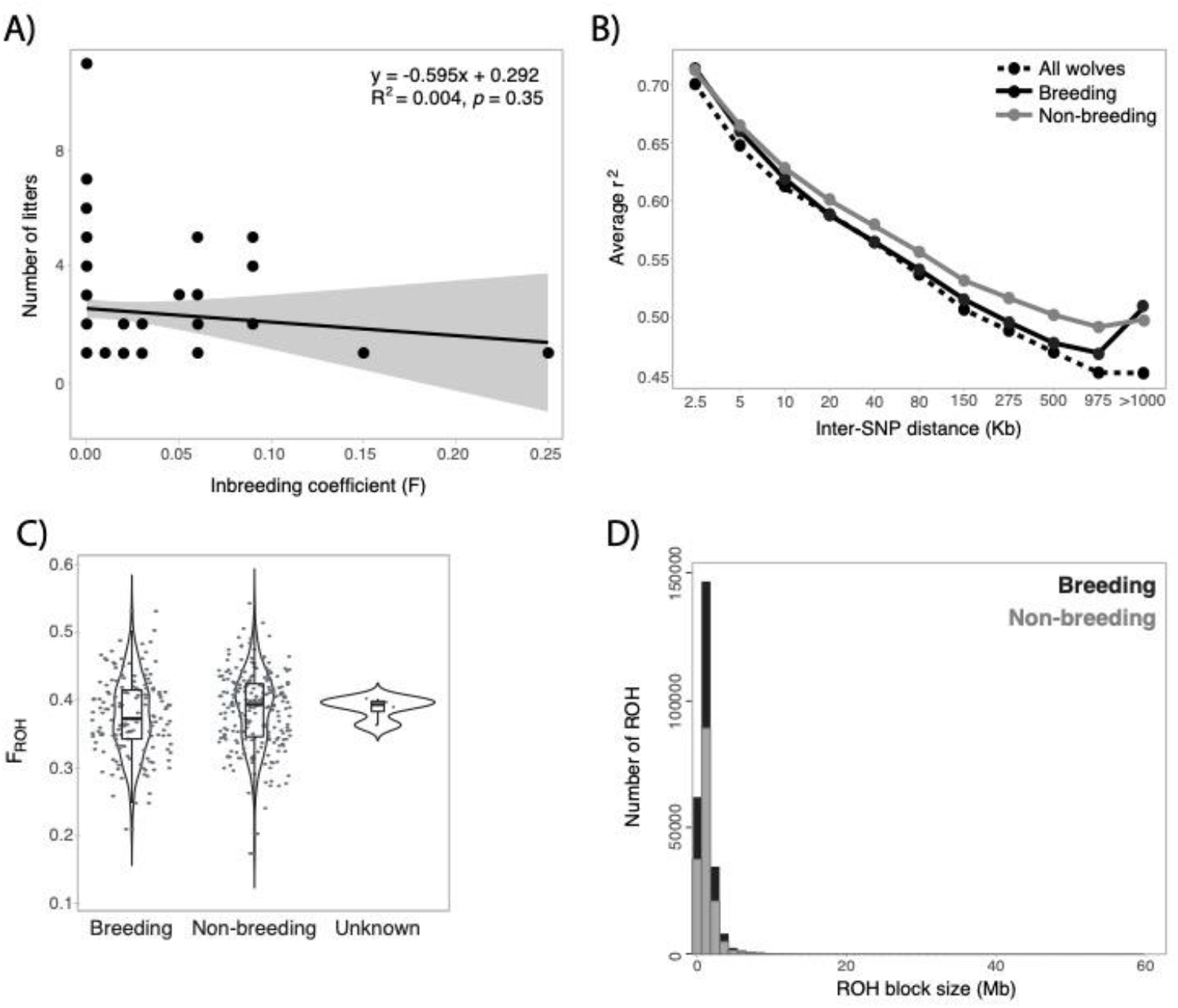
A) Number of litters (log10) as a function of pedigree-based inbreeding coefficient (*F*) estimates with data from 158 Yellowstone gray wolves (product moment correlation coefficient, r=-0.11; Spearman rank correlation coefficient, rs=0.27). **B)** Linkage disequilibrium decay for Yellowstone gray wolves (n, all=391, breeding=122, non-breeding=235, unknown=4) genotyped at 56K SNPs. **C)** Number of runs of homozygosity (ROH; minimum of 10 SNPs per 10Kb track) detected in 6Mb tracks and the related inbreeding coefficients (*F_ROH_*) estimated for 391 gray wolves. The length of the genome used for ROH estimates is 2 323 956 222bp for the pruned 24K SNP set. (Abbreviations: n, number of individuals). **D)** Violin box-and-whisker jitter plots of genome-wide inbreeding coefficients estimated from runs of homozygosity (*F_ROH_*) detected in 6Mb tracks for 24K pruned SNPs (minimum of 10 SNPs per 10Kb track). Welch two sample t-test statistics are provided.

### Lower inbreeding coefficients in the effective population

We obtained genome-wide SNP-based estimations for linkage, heterozygosity, and inbreeding for 391 Yellowstone gray wolves, 387 of which have known breeding status within the study period of 1995-2018 (Table S2). Linkage disequilibrium in the total YNP wolf population extended for an average of 150Kb before decaying below *r^2^*=0.5, with a similar length of LD decay found in the breeding population (230Kb) and substantially longer LD decay in non-breeding wolves (750Kb) (Fig 1B). We found no significant differences in heterozygosity between wolves with a confirmed breeding status and non-breeding wolves across the 56K SNP genotypes (Average±s.d. H_O_: breeding=0.792±0.02, non-breeding=0.794±0.02, 1-tailed *t*-test of equal variance *p*=0.0917). However, the breeding population had significantly lower levels of inbreeding estimates than non-breeding individuals (F_ROH_: breeding=0.376±0.06, non-breeding=0.387±0.06, *p*=0.0462) (Fig. 1C). We found statistical evidence for enrichment of longer ROH blocks in non- breeding individuals relative to the effective population (mean=1,311,428bp and 1,347,336bp, respectively; 1-tailed Mann-Whitney *U*=7868×10^9^, *p*=0.8.854×10^-5^) (Fig. 1D).

We observed stable annual heterozygosity estimates that were not significantly different between breeding and non-breeding wolves for 21 of the 24 years surveyed, likely due to small intra-annual genetic sample size relative to the census size (Table 4). During three years (2011-2013), non-breeding wolves carried significantly higher observed heterozygosity than the effective population (*p∼*0.05), although this trend can be noted in nearly every year surveyed and should be interpreted with caution given the small annual sample sizes. A similar but opposite trend was noted for inbreeding coefficients estimated from *F_ROH_*, where non-breeding wolves had a significantly higher inbreeding coefficient than breeding wolves for a single year (2012) (*F*=5.64, *p*=0.0210) (Table 4).

**Table 4.** Annual population averages of observed heterozygosity (H_O_) estimated from 56K SNPs and inbreeding coefficients (*F_ROH_*) from 24K pruned SNPs genotyped in breeding and non-breeding gray wolves of Yellowstone National Park (B and NB, respectively) with known years of birth and death. When census size also includes individuals of unknown breeding status. Significance was assessed by a 1-way ANOVA. (Abbreviations: N_G_, number of individuals genotyped; NS, not significant)

| Year | $N_B, N_{NB} (N_G)$ | $H_O$ | $F_{ROH}$ | ANOVA $F, p$ |
| --- | --- | --- | --- | --- |
| 1995 | 23, 6 (29) | 0.78, 0.79 | 0.34, 0.31 | NS |
| 1996 | 27, 16 (43) | 0.78, 0.79 | 0.33, 0.34 | NS |
| 1997 | 38, 24 (62) | 0.78, 0.79 | 0.35, 0.36 | NS |
| 1998 | 37, 28 (65) | 0.79, 0.80 | 0.36, 0.39 | NS |
| 1999 | 36, 35 (71) | 0.79, 0.80 | 0.37, 0.39 | NS |
| 2000 | 49, 37 (86) | 0.79, 0.79 | 0.36, 0.37 | NS |
| 2001 | 50, 37 (88) | 0.79, 0.79 | 0.37, 0.38 | NS |
| 2002 | 52, 39 (92) | 0.79, 0.79 | 0.37, 0.38 | NS |
| 2003 | 58, 44 (103) | 0.79, 0.80 | 0.38, 0.36 | NS |
| 2004 | 58, 52 (111) | 0.79, 0.80 | 0.38, 0.38 | NS |
| 2005 | 52, 49 (101) | 0.80, 0.80 | 0.38, 0.39 | NS |
| 2006 | 57, 36 (93) | 0.80, 0.80 | 0.39, 0.39 | NS |
| 2007 | 62, 46 (108) | 0.79, 0.79 | 0.38, 0.39 | NS |
| 2008 | 57, 42 (99) | 0.79, 0.79 | 0.39, 0.38 | NS |
| 2009 | 56, 27 (83) | 0.80, 0.79 | 0.39, 0.40 | NS |
| 2010 | 48, 26 (74) | 0.80, 0.80 | 0.39, 0.40 | NS |
| 2011 | 46, 29 (75) | 0.80, 0.81 | 0.39, 0.41 | $H_O$ 4.19, $p=0.0444$ |
| 2012 | 34, 24 (58) | 0.79, 0.81 | 0.38, 0.42 | $H_O$ 9.58, $p=0.0031$;<br>$F_{ROH}$ 5.64, $p=0.0210$ |
| 2013 | 33, 13 (46) | 0.79, 0.80 | 0.38, 0.40 | $H_O$ 3.80, $p=0.0576$ |
| 2014 | 32, 18 (50) | 0.79, 0.79 | 0.39, 0.40 | NS |
| 2015 | 28, 18 (46) | 0.79, 0.79 | 0.39, 0.39 | NS |
| 2016 | 23, 19 (42) | 0.79, 0.79 | 0.39, 0.40 | NS |
| 2017 | 20, 20 (40) | 0.79, 0.80 | 0.39, 0.40 | NS |
| 2018 | 12, 12 (24) | 0.79, 0.79 | 0.40, 0.40 | NS |

### Individuals with higher autozygosity estimates are older at first litter

We restricted the dataset to explore fitness and genetic data for 387 pedigreed wolves with sufficient life history information (known years of birth and known years of reproduction, death or last observation). We found that lifespan (average breeding=6.1 and non-breeding=2.1 years, 1-tailed t-test *p*=9.31×10^-43^) was on average higher in the effective population concomitant with lower inbreeding coefficients (average breeding=0.37 and non- breeding=0.38 years, 1-tailed t-test *p*=0.0466), and that breeding wolves were on average 2.9±1.2 years old (males=3.2±1.3, females=2.7±1.1; 1-tailed t-test *p*=0.0086) at the time of their first litter (Figs. 2A,B). However, several life- history and genetic traits show strong correlations, with positive correlations noted between lifespan, age at first litter, and number of litters while inbreeding coefficients were negatively correlated (Fig. 2C).

**Figure 2.**
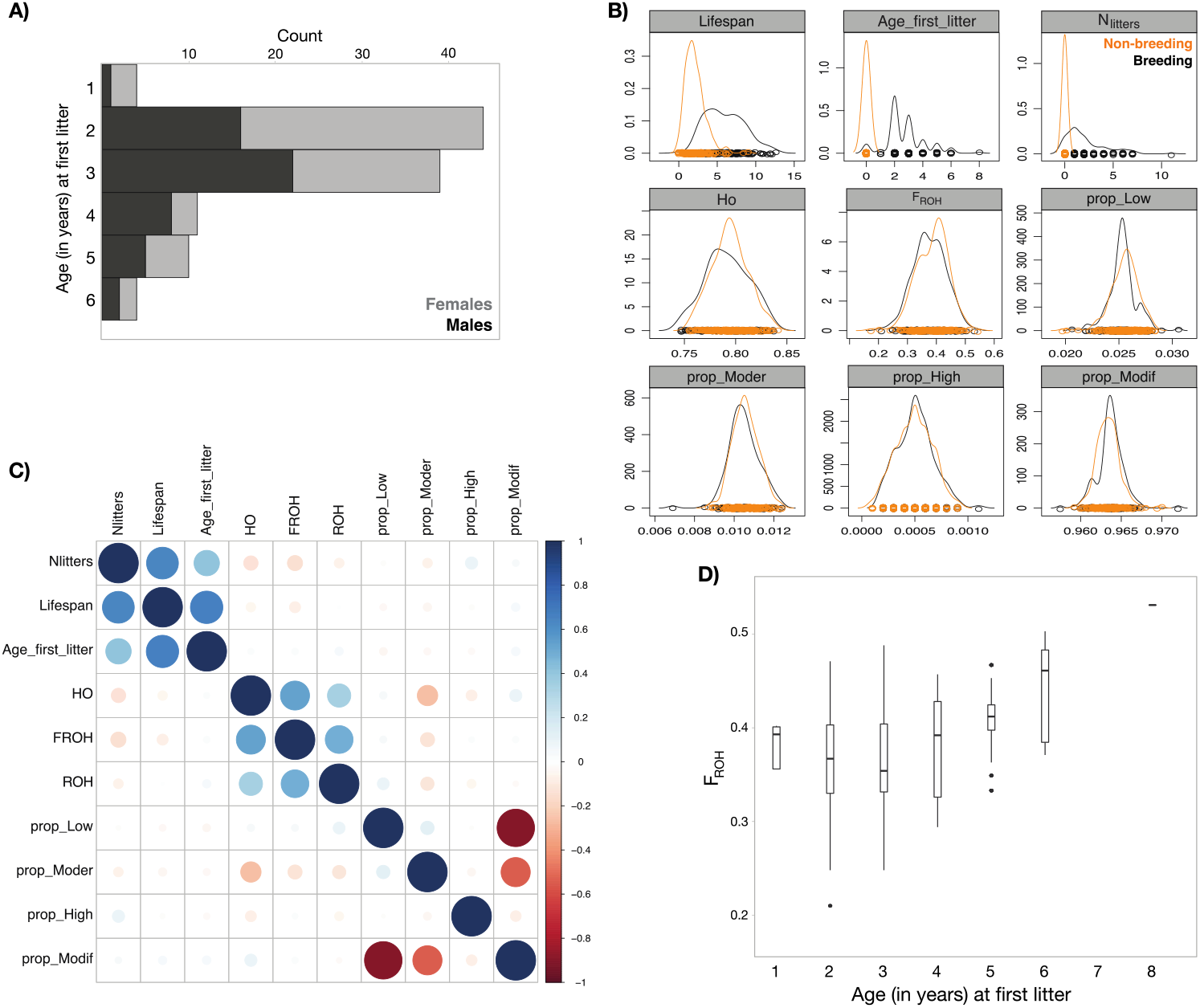
A) Distribution of the number of individuals per age at event of first litter (age_first_litter) when born after 1995, stratified by sex. The **B)** density plot of traits with respect to reproductive status (Abbreviations: N_litters_, number of litters; Ho, observed heterozygosity, prop_Low, proportion of SNPs with low predicted impact; prop_Moder, proportion of SNPs with moderate predicted impact; prop_High, proportion of SNPs with high predicted impact; prop_Modif, proportion of SNPs predicted to be modifiers), and **C)** pairwise correlation plot (scale bar indicates direction and magnitude of correlation) of life-history traits for 276 wolves with known year of birth and death. **D)** Box- and-whisker plot of genome-wide inbreeding coefficients estimated from runs of homozygosity (*F_ROH_*) as a function of age at first litter.

We further explored the relationship of similar genetic parameters with respect to a wolf’s age at first litter using Cox proportional hazards survival analysis. In all analyses we included sex as a covariate along with the first five PCs from SNPs covering the genome to account for the relatedness; similar results were obtained without these covariates (results not shown). There were 139 animals with known ages at their first litter and 235 non-breeding animals with known age at death or last observation for a total of 374 animals with PCs and phenotype data used in this analysis. Individuals who had their first litter at an older age had higher inbreeding coefficients (Fig. 2D). We found that a higher inbreeding coefficient significantly reduced the Cox proportional hazard and was associated with later reproduction. Thus, *F_ROH_* was a significant predictor of age at first litter (Log hazard coefficient=-0.2406, *p=*0.0203 Table 5). Sex was also significant, with males having a later age to their mate’s first litter than females (Log hazard coefficient=-0.5674, *p=*0.0012 Table 5).

**Table 5.**
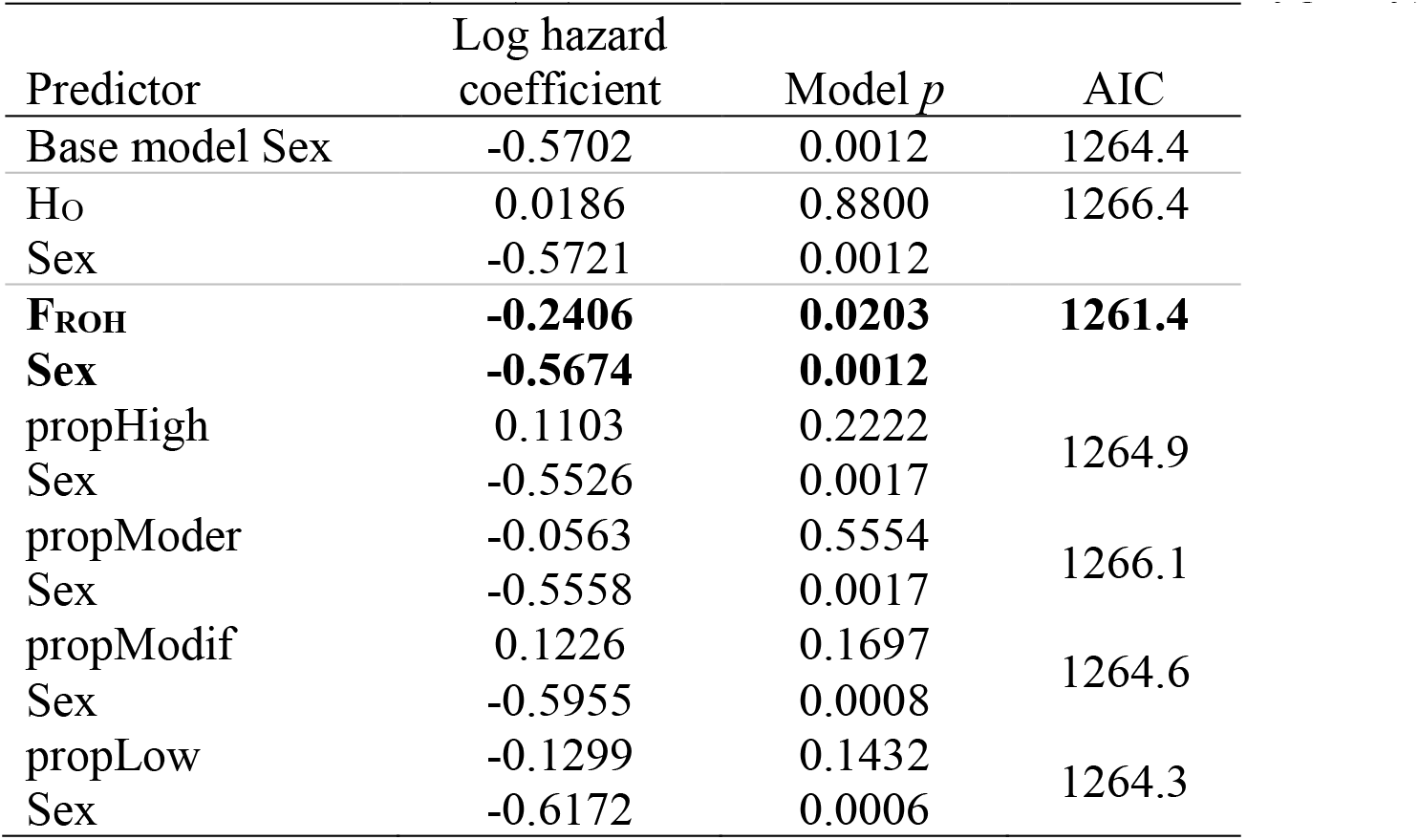
Cox proportional hazards regression with age at first litter as the outcome response. Time is recorded in years and represents time to first litter (n=139) or time to last observation or death (n=238) whichever is sooner. Data from animals who die or are no longer observed before reproducing are considered censored. The predictors are time invariant and have been standardized. The best fitting model (bolded) has the lowest Akaike information criterion (AIC). (Abbreviations: H_O_, observed heterozygosity)

## Discussion

The reintroduced gray wolves of Yellowstone National Park are unique in that they are observed every single year, with life events documented and supported by a wealth of molecular data. Despite the rising accessibility of genome sequencing, Yellowstone wolves stand among only a few other systems with similar multidimensional datasets that merge static, dynamic, and molecular perspectives (Stahler et al. 2020). We explored this interface by conducting a pedigree- and marker-based assessment of genetic variation over time within the effective population and modeled fitness-related traits. The pedigree is large and complex, capturing seven generations since the reintroduction of gray wolves to the Rocky Mountains of the United States in 1995 and 1996. Although Canadian founders were carefully selected and indeed have contributed to the genetic success of the population, equally important was the translocation of gray wolf pups from northwest Montana in 1997 that carried divergent genetics (vonHoldt et al. 2008, 2010). As two of these pups matured and assumed a social rank with reproductive access, this lineage continues to provide an influx of genetic variation distinct from the Canadian founders into YNP wolves through their descendants. This finding illustrates the critically positive impact that a few successful breeders have on gene flow into a population. Further, this is exemplary as a design for reintroduction programs to establish a genetically diverse founding population.

Given the known genealogy of this population, we identified a handful of founders that contributed a significant amount of kinship and genome uniqueness, with over 87% of the founders’ genomes persisting in the pedigree. The remaining fraction was lost through random drift or emigration. As expected for group-living species, mean kinship significantly increased over time; however, this appears to be mitigated by the continued reproductive success of the translocated individuals from northwest Montana (now the Nez Perce lineage). Further, concordant with vonHoldt et al. (2008), mate choice and inbreeding avoidance also mitigates the degree to which mean kinship increased over time in the pedigree, which is expected to be significantly higher under random breeding alone. We found that the number of effective founders and genomic uniqueness increased over time; this positive consequence is expected in a social species with structured mating, which can alleviate the potentially strong impacts of the reintroduction event which inevitably forces a population genetic bottleneck.

We utilized genome-wide SNP data to explore the distribution of genetic variation across the pedigree relative to time, reproduction, and fitness- related traits (e.g. litters, inbreeding). Overall, we document high retention of genetic variation over the 25 years since reintroduction. We found that the effective population of breeding wolves carried a significantly shorter distances at which LD decays. This is consistent with past findings that the YNP effective gray wolf population carried a higher mean allelic richness, which is expected to show more immediate changes than other allele diversity metrics like heterozygosity, than the census in nearly all years surveyed (1995- 2015) (Cornuet and Luikart 1996; DeCandia et al. 2021).

We hypothesized that reproductive status reflected overall fitness and predicted that the effective population showed increased genetic variation, reduced autozygosity levels, and that fitness-related traits reduced in value with increasing autozygosity levels. Elevations of autozygosity levels indicate both a lack of genetic diversity and inflation of alleles that are identical by descent via inbreeding and are often associated with increased expression of recessive deleterious traits (Keller 2002; Charlesworth and Willis 2009).

Indeed, we found remarkably lower levels of autozygosity and inbreeding coefficients in the effective population relative to the non-breeding population. Such a trend is far more significant on overall inference of genetic health than levels of homozygosity. Our finding is in direct contrast to the gray wolf population in Isle Royale National Park (IRNP; Michigan, USA), which had persisted at exceedingly low numbers for over a decade with negative fitness and genome-wide consequences recently documented (Robinson et al. 2019). We do not suggest the genetic health of YNP wolves is similar to IRNP; rather, they represent differences along the complex gradient of interactions between effective population size, genetic isolation, and fitness.

The suggested traits impacted by autozygosity in YNP gray wolves were reflected in reproduction, a fitness consequence concordant with studies of IBD and complex diseases in humans (Ceballos et al. 2018; Pemberton et al. 2018; Szpiech et al. 2013). After controlling for sex-specific differences in primiparity, individuals that are older at the time of their first litter carried higher levels of autozygosity, which was also positively associated with a significant reduction in the total number of litters for that individual. Proximate mechanisms for why these older individuals were unable to breed during previous years may be due to intra-pack composition (e.g. lower social status or no access to unrelated mates) or challenges with finding potential mates through dispersal. Ultimately, individuals with higher levels of autozygosity that breed later in life may be losing out to higher quality competitors before eventually breeding, indicating a fitness cost. Sex-specific differences in age at first litter likely reflect differential breeding opportunities through male-biased dispersal and female natal philopatry documented in YNP wolves (Stahler et al. 2020; Smith et al. 2020).

Albeit such trait depression is difficult to acquire a large enough sample size for a well-powered study in natural populations, especially those with longer generation times, similar trends have been reported in other systems (Keller and Waller 2002). Nielsen et al. (2012) utilized a long-term pedigree- based design for meerkats (*Suricata suricatta*), a cooperative and group-living species, and found that increased inbreeding coefficients were associated with negative morphologic consequences for pups with higher inbreeding coefficients. Increased pathogen load was found in Soay sheep (*Ovis aries*) in sheep with reduced genetic diversity (Coltman et al. 1999) and depressed fitness across developmental stages for the endangered red-cockaded woodpecker (*Picoides borealis*) (Daniels and Walters 2000). We also recently found similar trends for the YNP wolves whereas reductions in genome-wide diversity estimates are associated with increased disease severity with respect to sarcoptic mange infections (DeCandia et al. 2021). While YNP wolves overall are genetically diverse, we detected reproduction consequences associated with moderate increases in autozygosity as a result of reduced genetic variation. Similar patterns of reproductive consequences were found with maternal inbreeding levels in red deer (*Cervus elaphus*) being correlated with reduced offspring survival, further suggesting that depression of fitness traits in adult individuals is likely to be more accurately estimated from marker-based data (Huisman et al. 2016). Chu and colleagues (2019) reported a negative correlation between fecundity and autozygosity in a longitudinal study of domestic dogs, which also supported pedigree-based assessments of such (LeRoy et al. 2015).

Surveying lethal equivalents (LE) assumes that trait depression is caused by deleterious recessive variation found in the homozygous state (or the equivalent of alleles across loci with a similar contribution towards trait depression). Pedigree-based estimates of inbreeding revealed a negative correlation to the number of litters, with an estimation that each wolf likely carries nearly two LEs in their genome, similar to previous estimates made for humans, *Drosophila*, and great tits (Szulkin et al. 2007; Gao et al. 2015). This directly translated to an expected decrease in offspring survival to maturity by 22% due to a full-sibling mating. Nietlisbach and colleagues (2018) conducted a literature survey and reported that wild vertebrate populations carried on average 3.5 LEs. However, 13 of the 18 studies analyzed represented bird species, likely across a diversity of mating systems relative to the highly structured one of gray wolves.

### Long-term implications

These results provide a valuable baseline through which continued monitoring can evaluate long-term genetic health of wolves in YNP. Similar approaches could be applied across a larger geographic scale to monitor genetic health and connectivity of wolf populations throughout the contiguous U.S., a task that is identified in the final ESA delisting requirements (USFWS 2020). In light of our findings, the future genetic health of wolves in YNP depends upon the critical role of gene flow and preserving landscape corridors to support effective dispersal. The YNP wolf population, which represents the core of a larger population throughout the Greater Yellowstone Ecosystem, serves as an important source for wolves that disperse beyond the protective boundaries (vonHoldt et al. 2010). As the YNP wolf population has stabilized at lower densities over time, the pedigreed population exhibited an increase in genome uniqueness (e.g. Table 3), suggestive of successful effective dispersal that has been documented through field observations of radio-collared dispersers (Stahler et al. 2020). Although this study does not evaluate gene flow into YNP, our findings suggest that immigration of effective dispersers over time will be essential for safeguarding their future through the incorporation of new and adaptive genetic variation. Such variation provides an important mechanism for rapid evolutionary response to changing environmental challenges caused by disease, climate change and human alteration of habitats (Kardos et al. 2021).

To maintain larger-scale wolf population connectivity and counteract loss of genetic variation, natural dispersal dynamics should be promoted and anthropogenic factors that significantly reduce genetic connectivity and effective population size should be mitigated (vonHoldt et al. 2010). These goals face increased challenges under recent wolf management directives in some northern Rocky Mountain states that aim to significantly reduce the number of wolves on the landscape. For example, recent legislative actions in Montana (2021 Montana Code Annotated 87-1-901) and Idaho (2021 Idaho Senate Bill No.1211) include aggressive policies to reduce wolf population sizes to levels close to minimum threshold requirements to prevent ESA relisting (e.g. Idaho SB 1211). These actions include regulations that allow for killing wolves using methods beyond shooting, such as baiting, snaring, trapping, night-time hunting on private land, aerial gunning, unlimited quotas, large numbers of wolf tags per hunter (20 wolves in Montana) or unlimited (Idaho), and extensive hunting seasons (e.g. year-round in Idaho and 6 months in Montana). If continued, these specific regulations have the potential to significantly reduce regional wolf population sizes, not only limiting gene flow into YNP, but disrupting the genetic connectivity that was demonstrated to have occurred across a larger regional scale following the first decade of wolf recovery in the western U.S. (vonHoldt et al. 2010). Such management policies that fail to incorporate larger meta-population dynamics of dispersal and inter- regional connectivity with adequate effective population sizes could jeopardize the tremendous success of wolf recovery efforts and genetic health in the Western United States over the last 25 years.

## Acknowledgements

We are grateful for funding support from Yellowstone Forever and their many donors, especially Valerie Gates and Annie and Bob Graham. We also are appreciative to the many field technicians who continue to document in amazing details the lives of wolves in Yellowstone National Park. JSS is partially funded by the NIH (R35GM141798, R01AI153044, R01HG009120) and the National Science Foundation (DMS-1264153, DEB-0613730, and DEB-1245373).

## Data Archiving

The previously published data analyzed in this study is publicly available on NCBI’s Sequence Read Archive (PRJNA577957). The mapped bam files for three newly collected individuals can be found at PRJNA743299 with additional metadata found in Supplemental Table S1.

---

**Supplemental Table S1.**
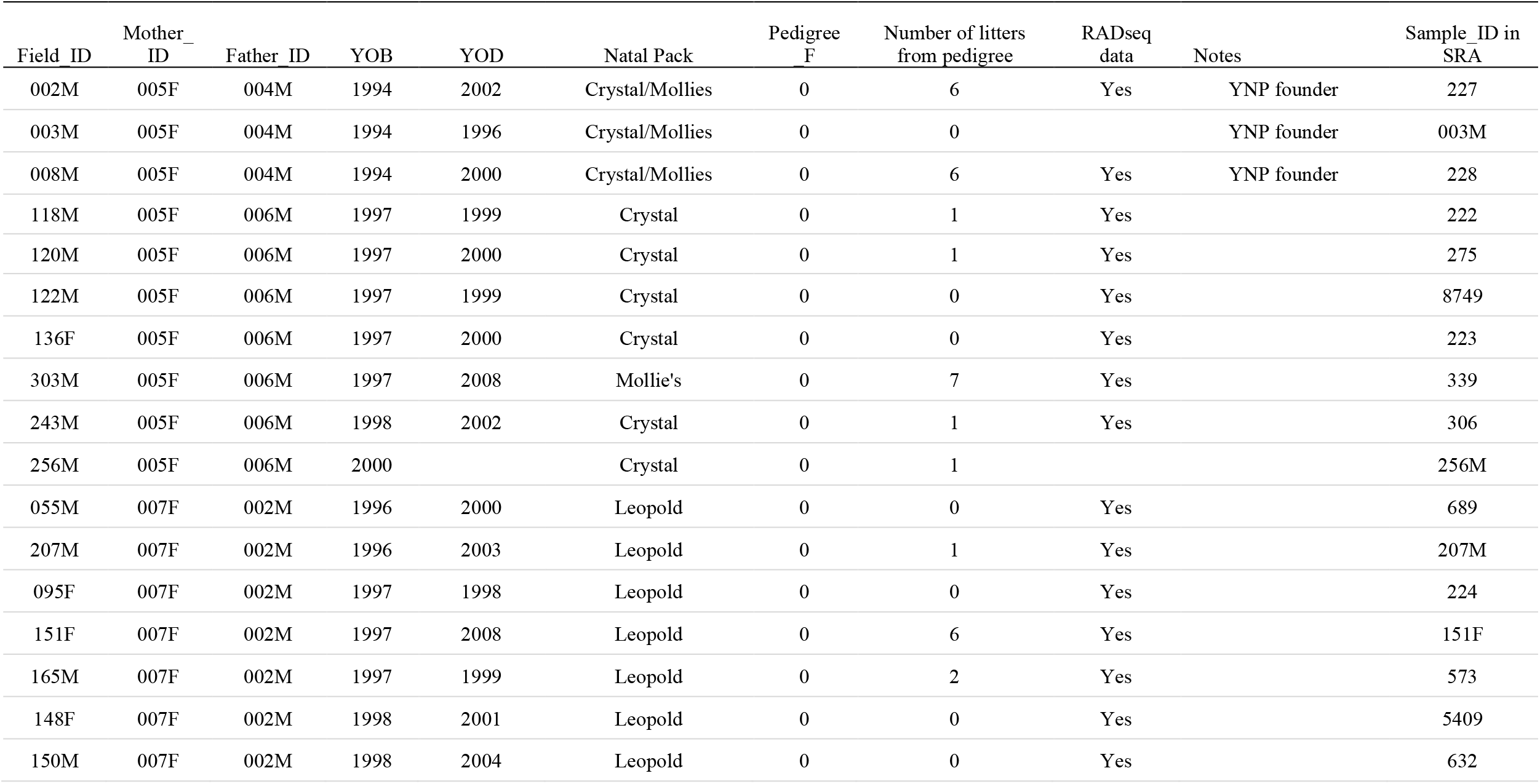
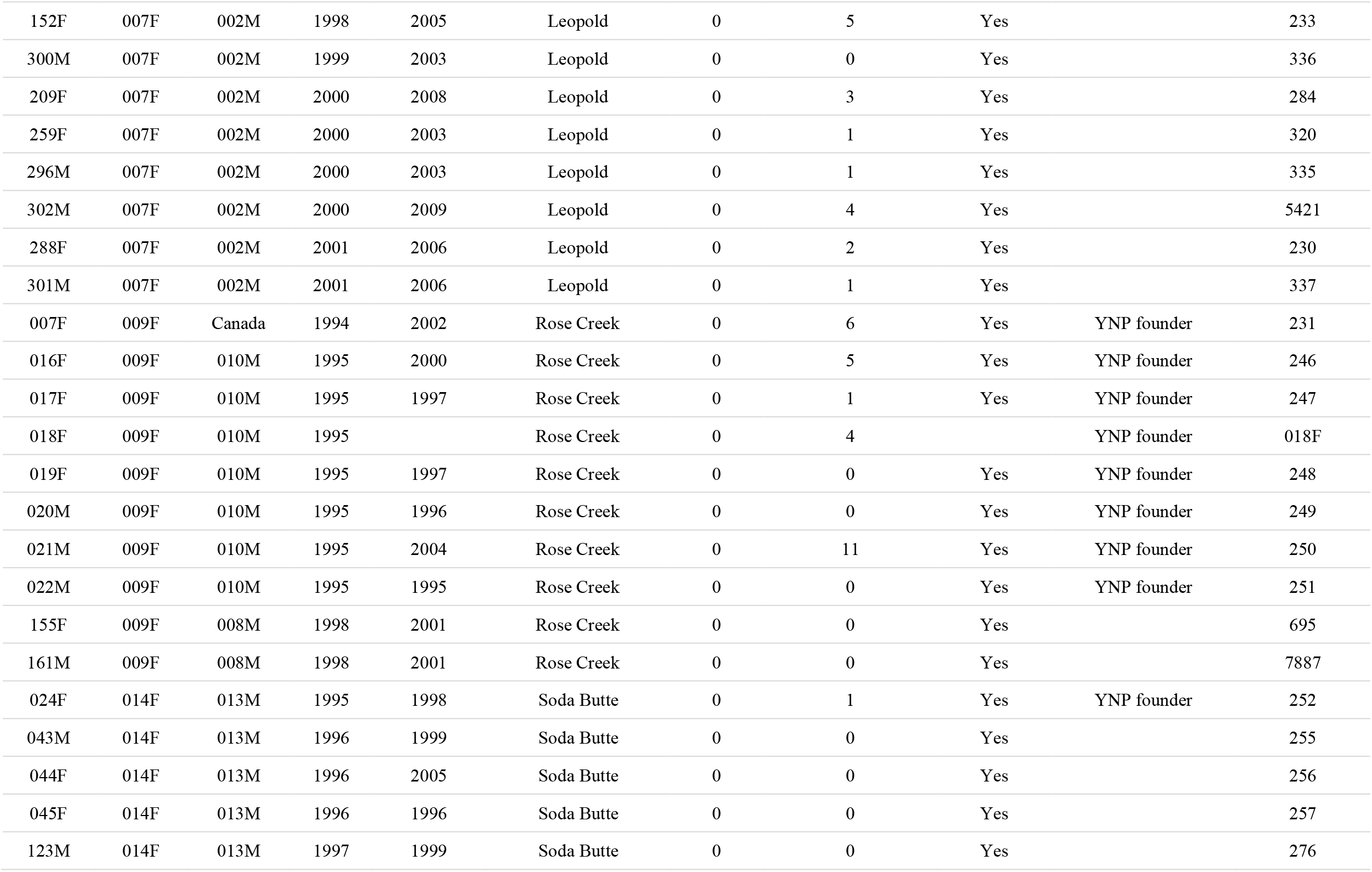
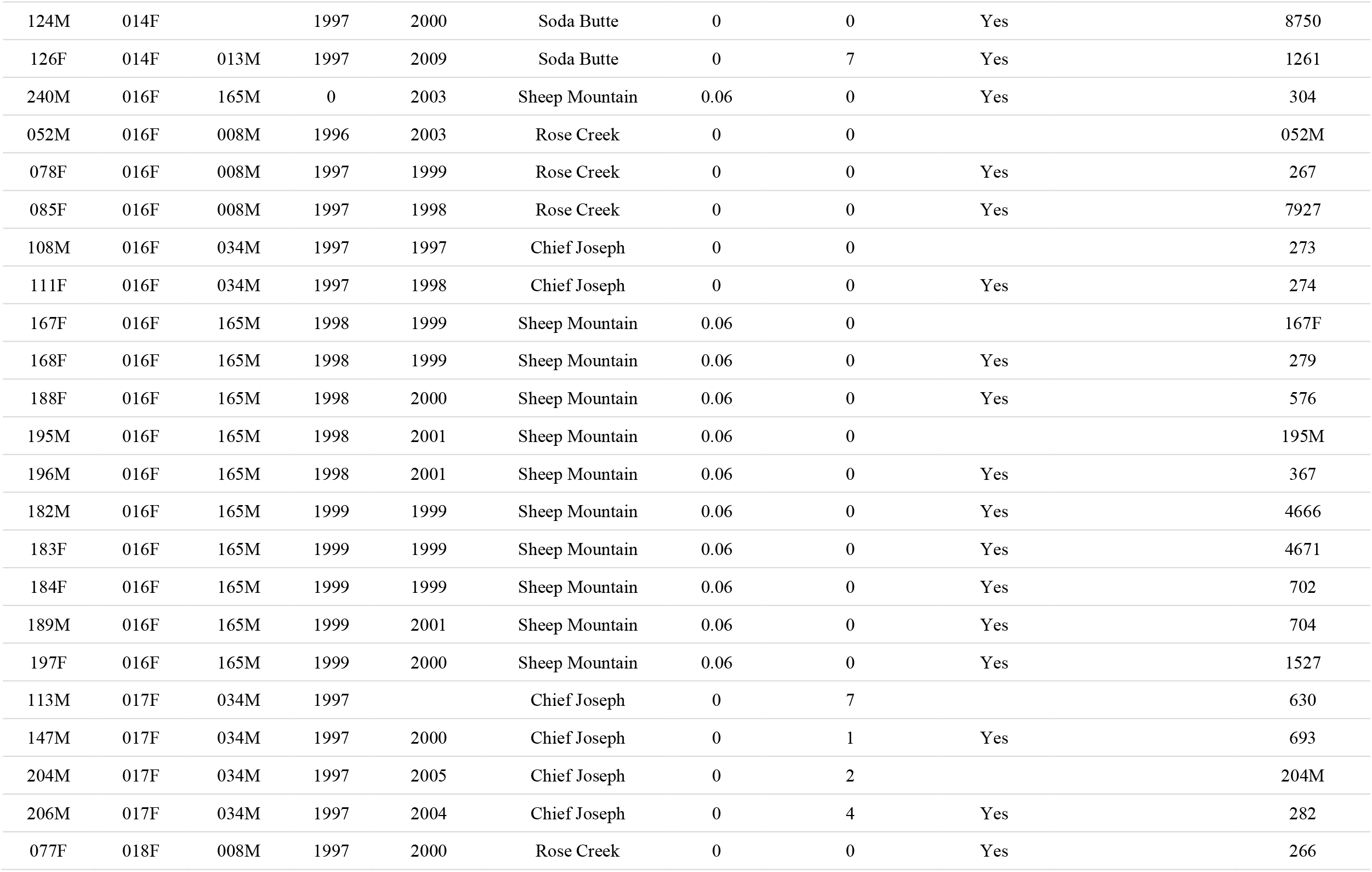
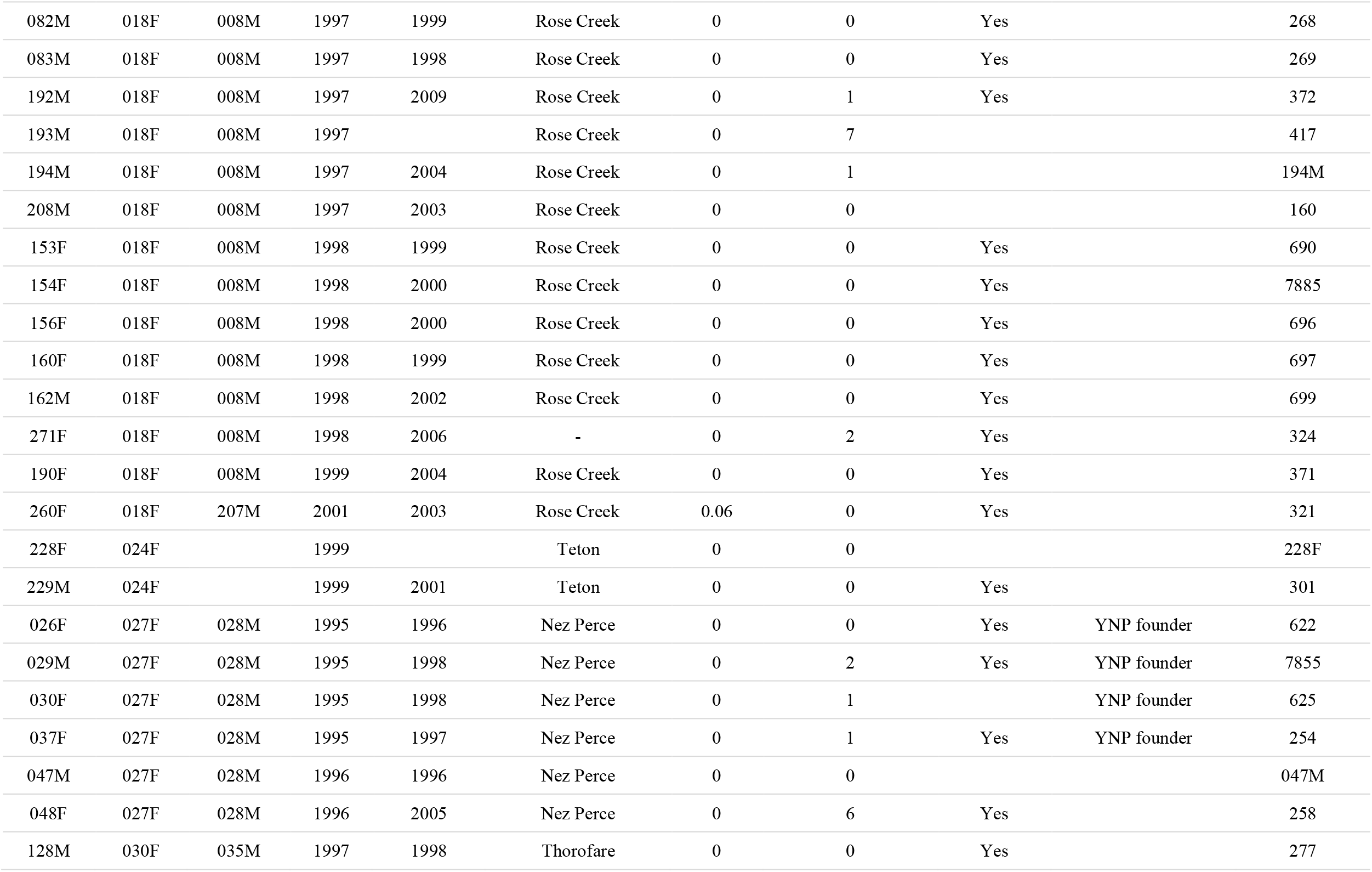
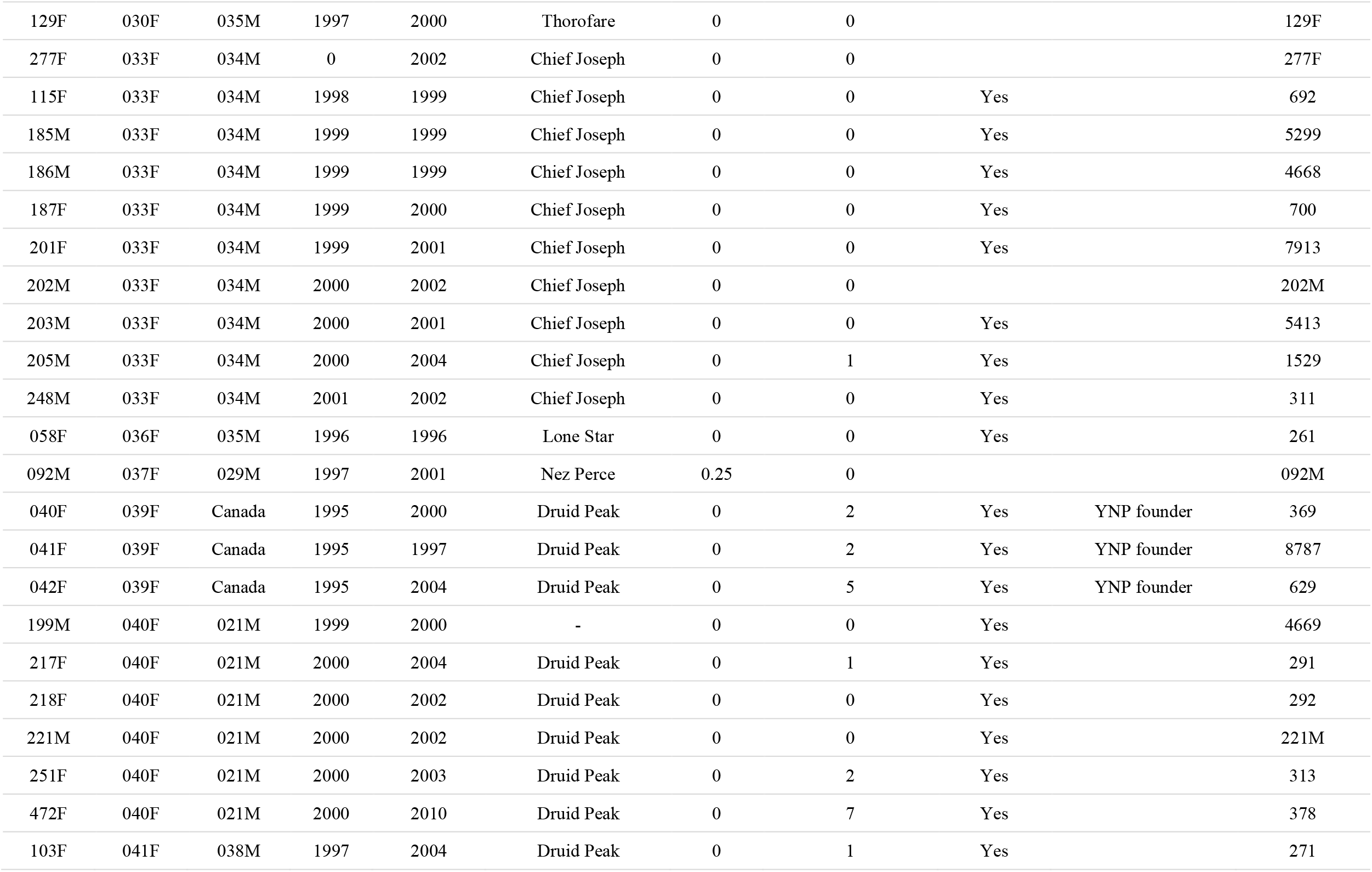
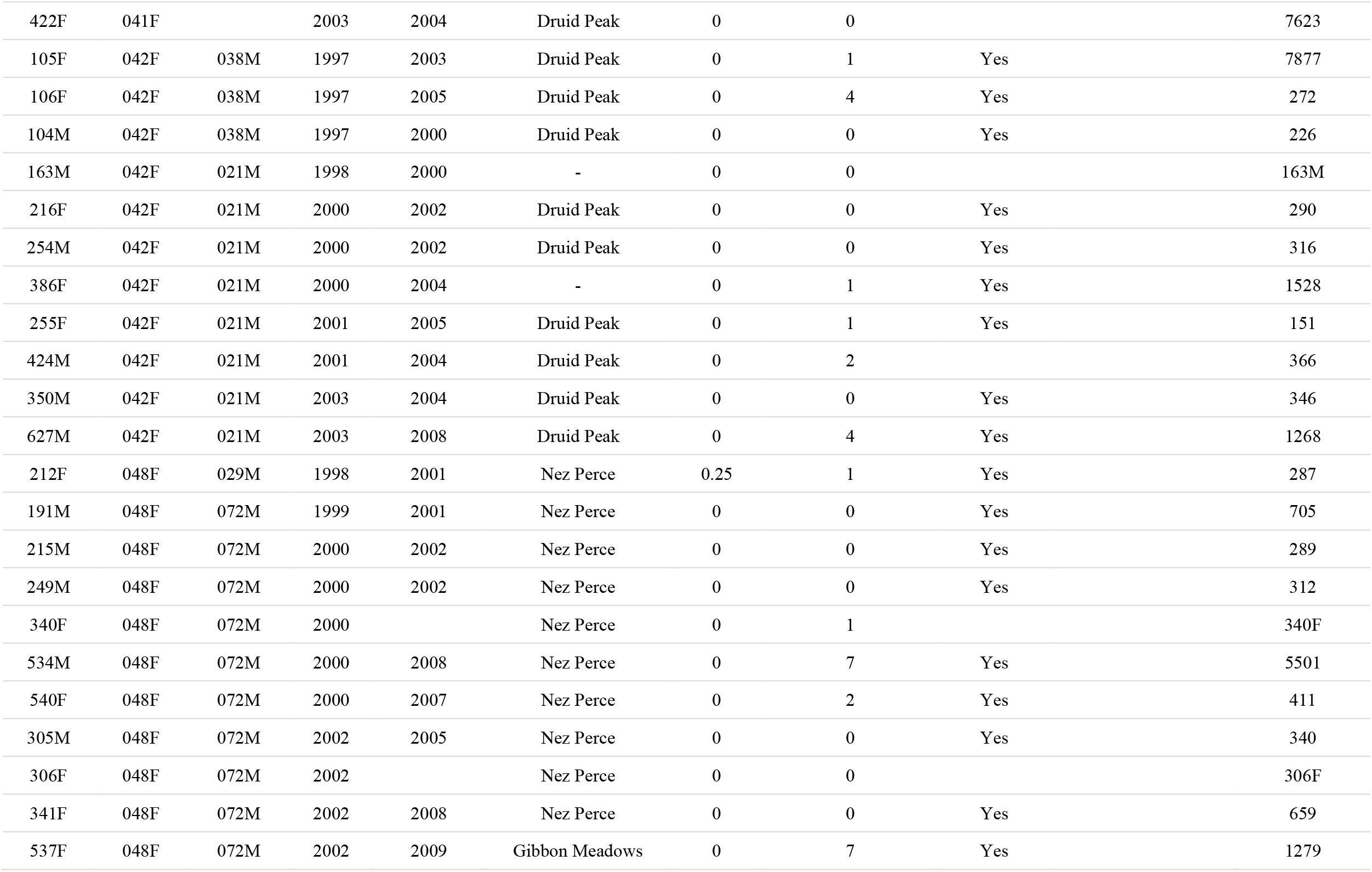
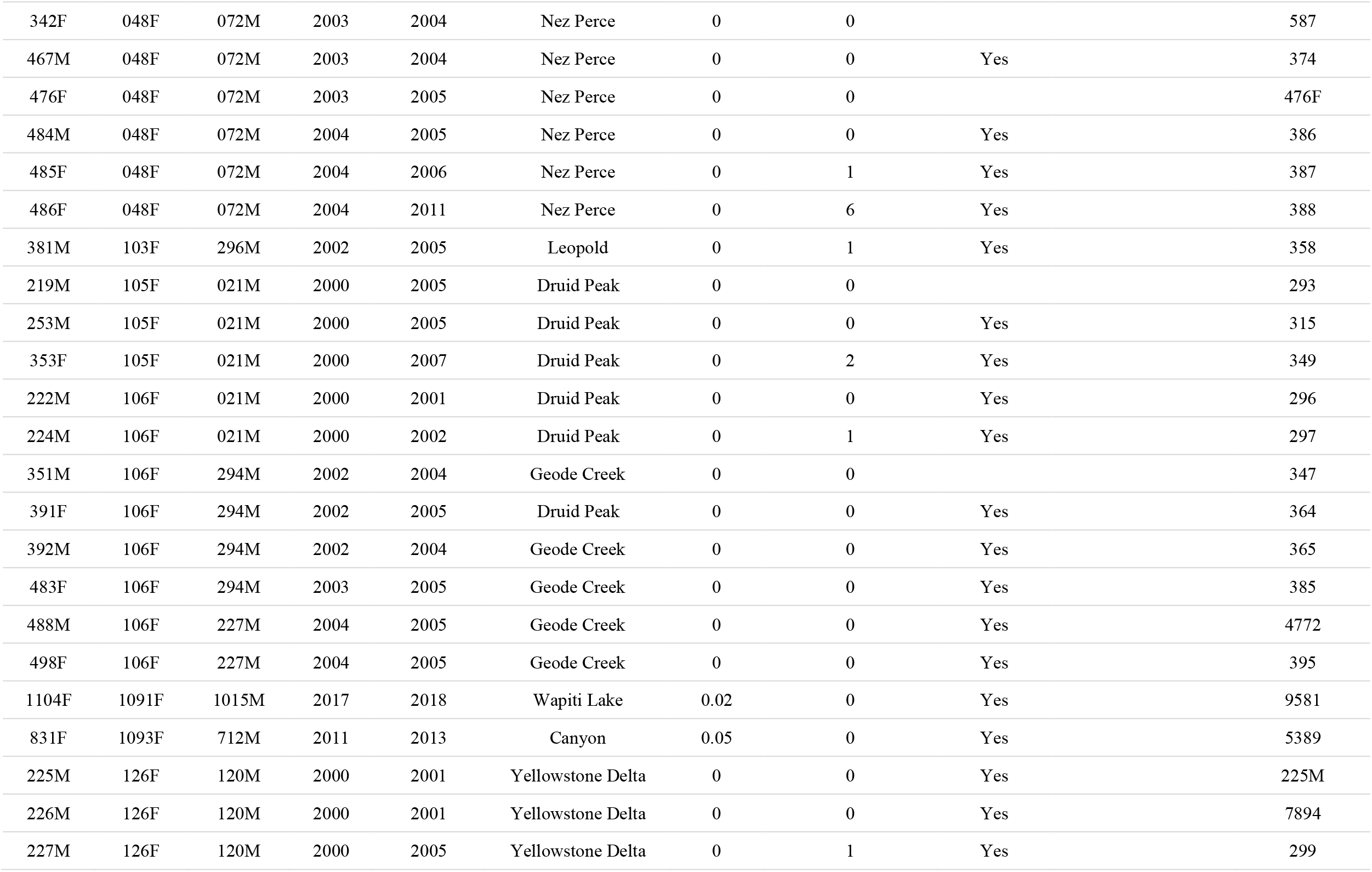
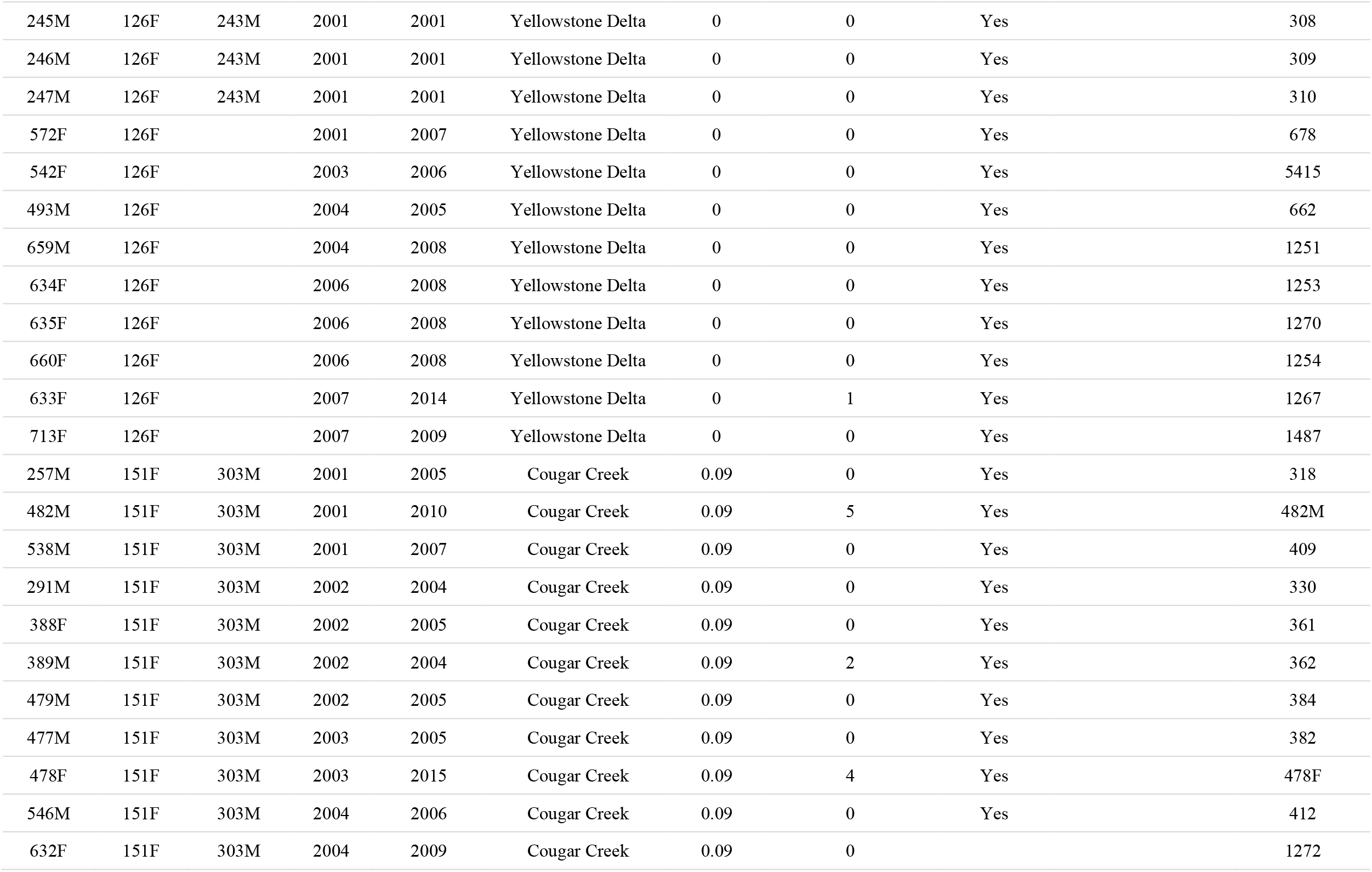
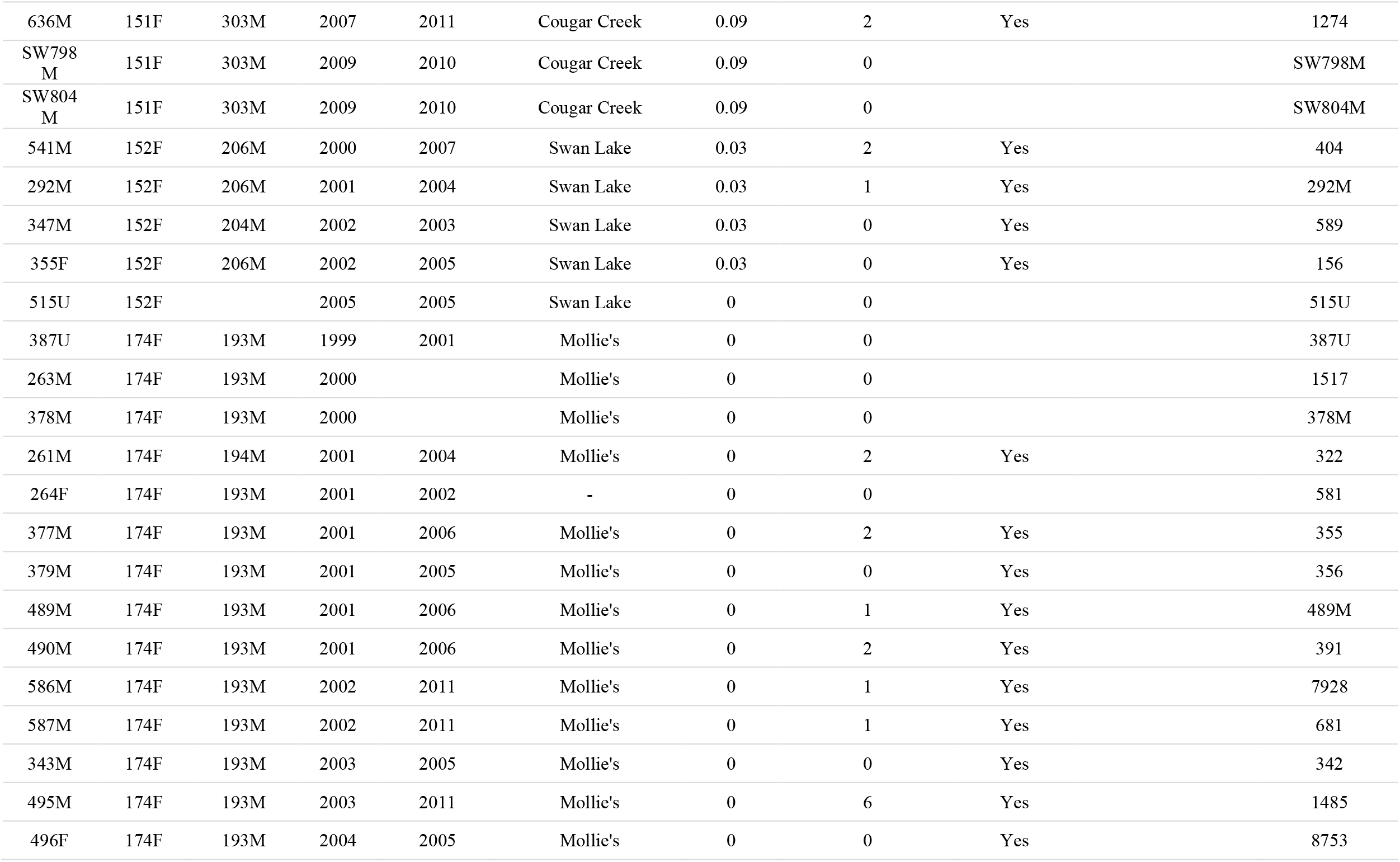
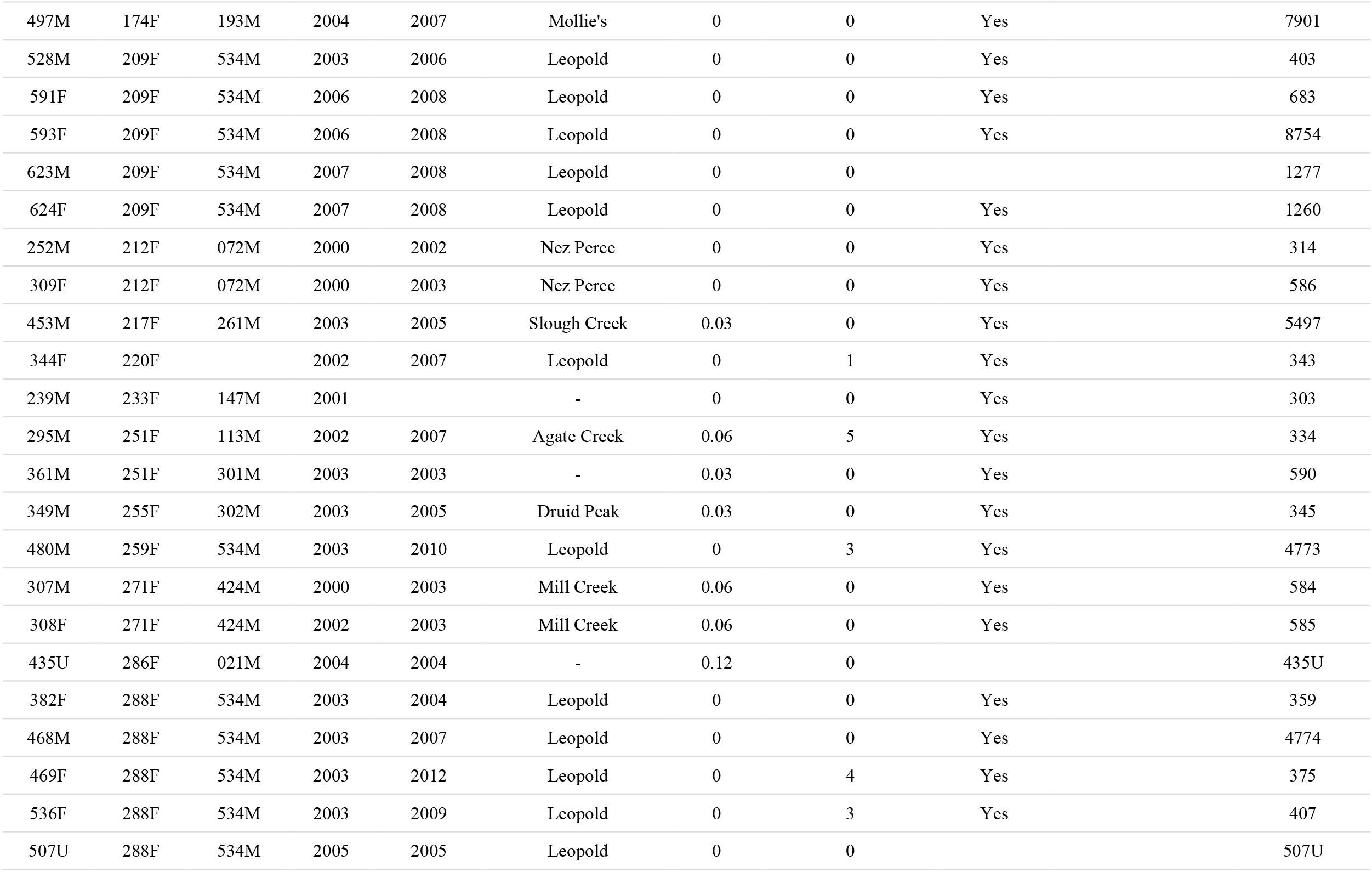
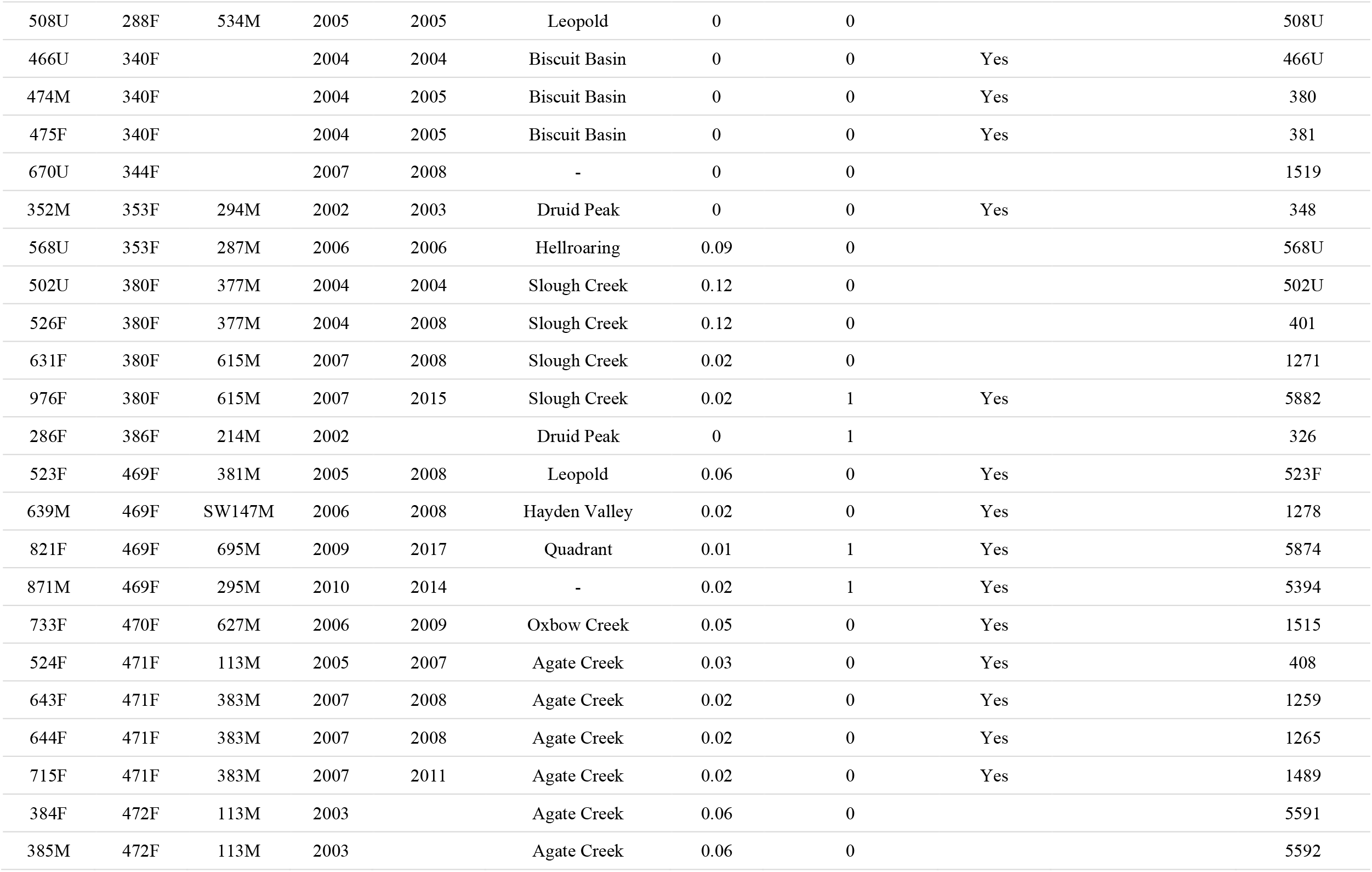
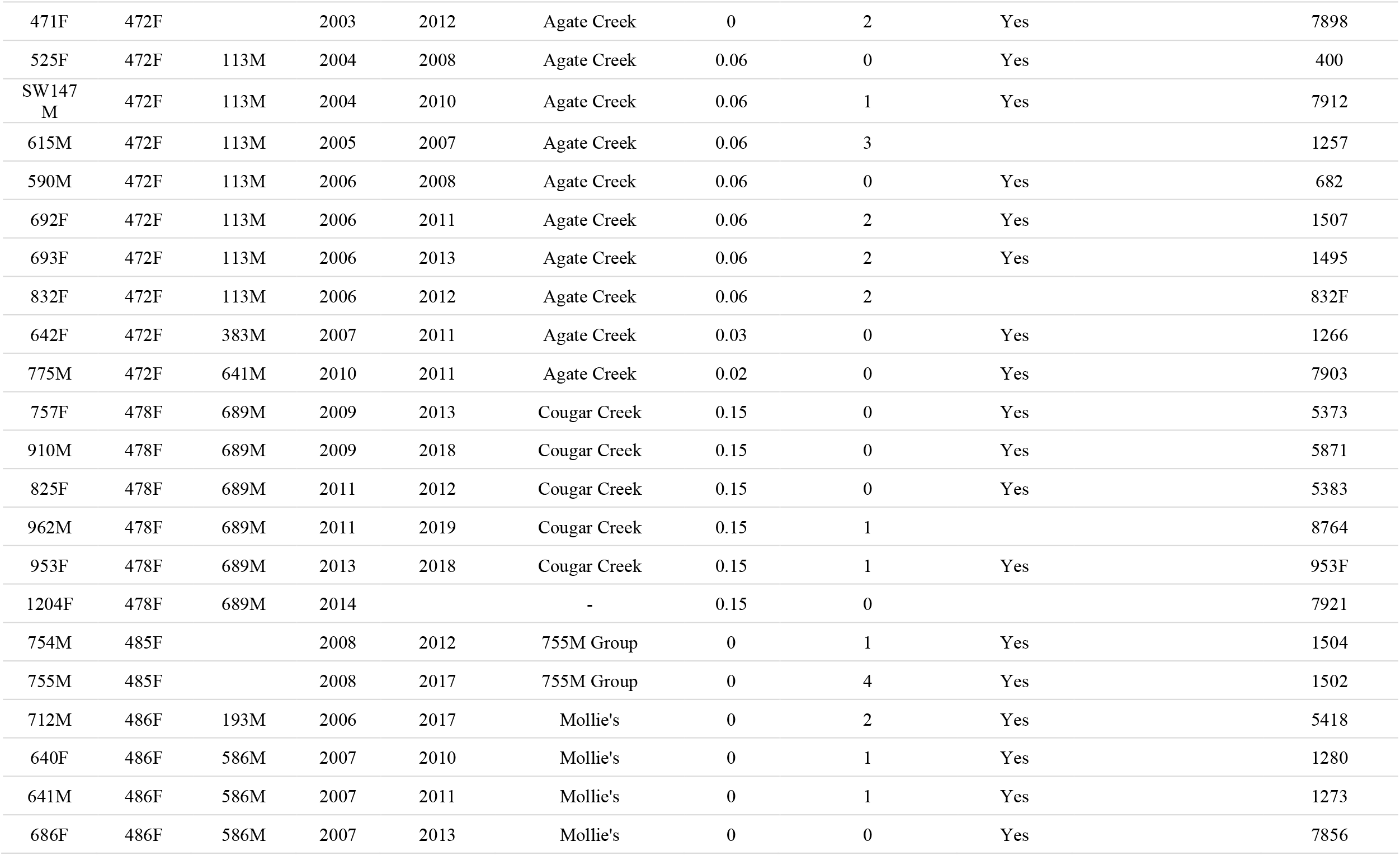
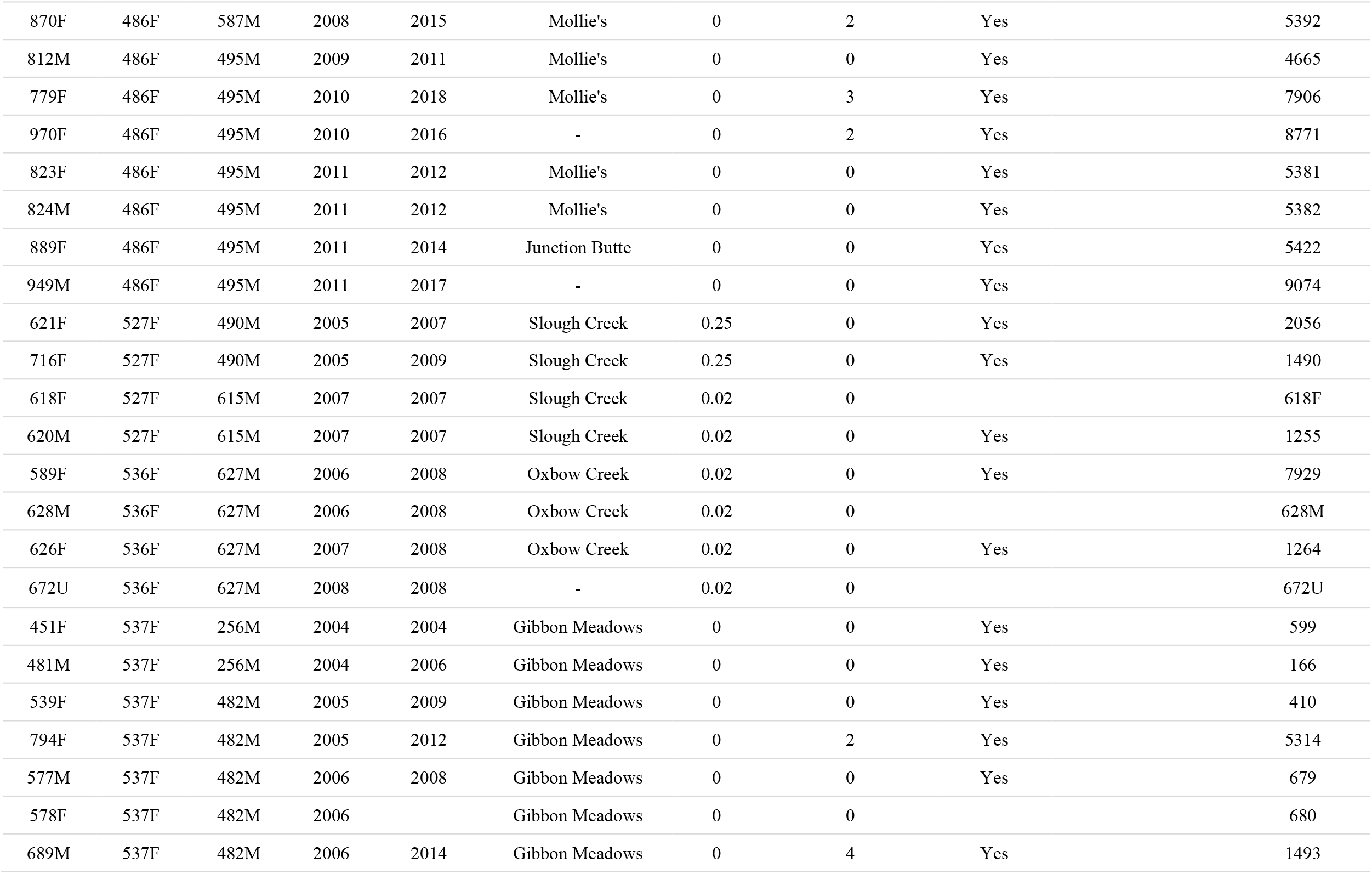
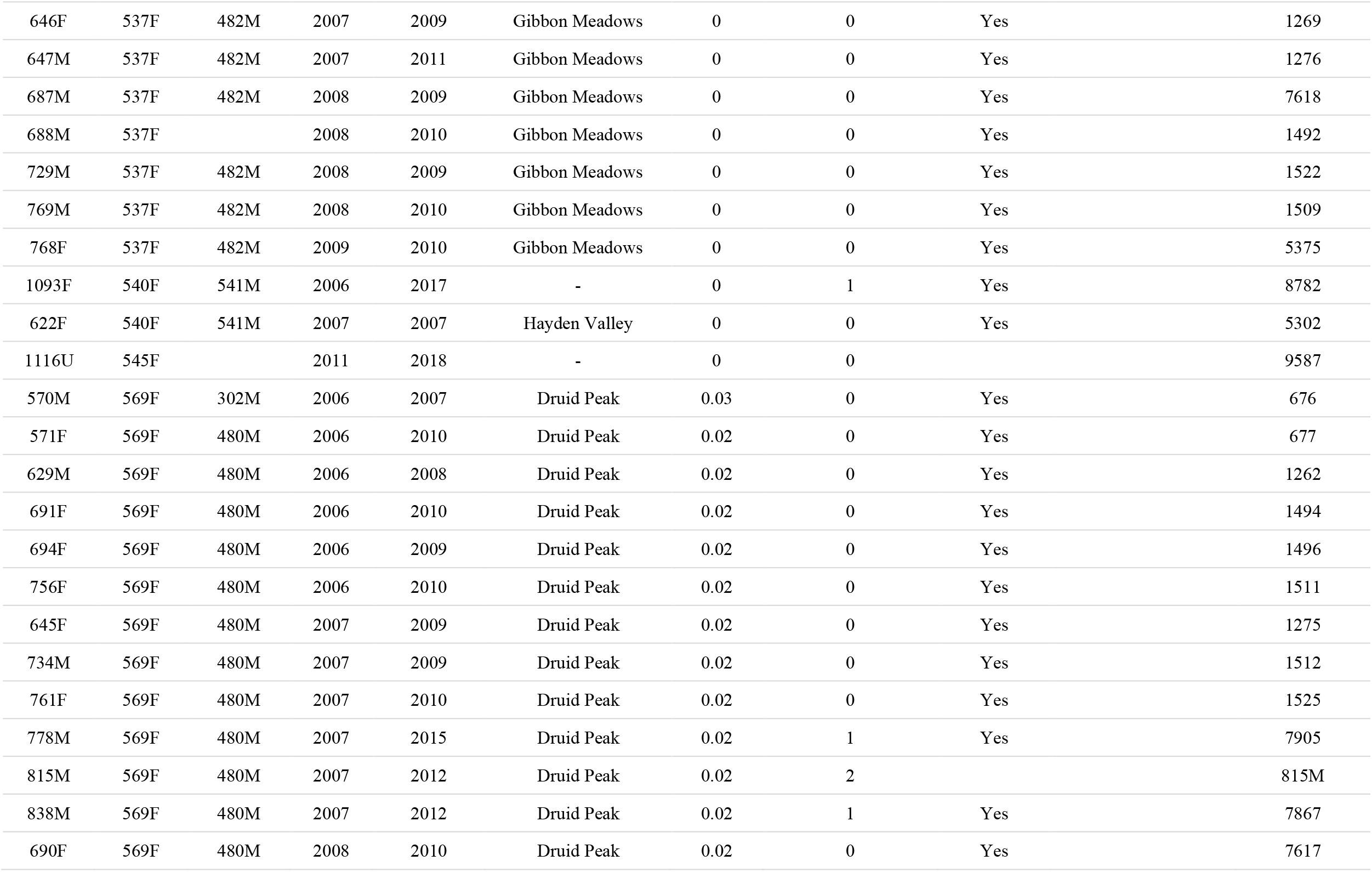
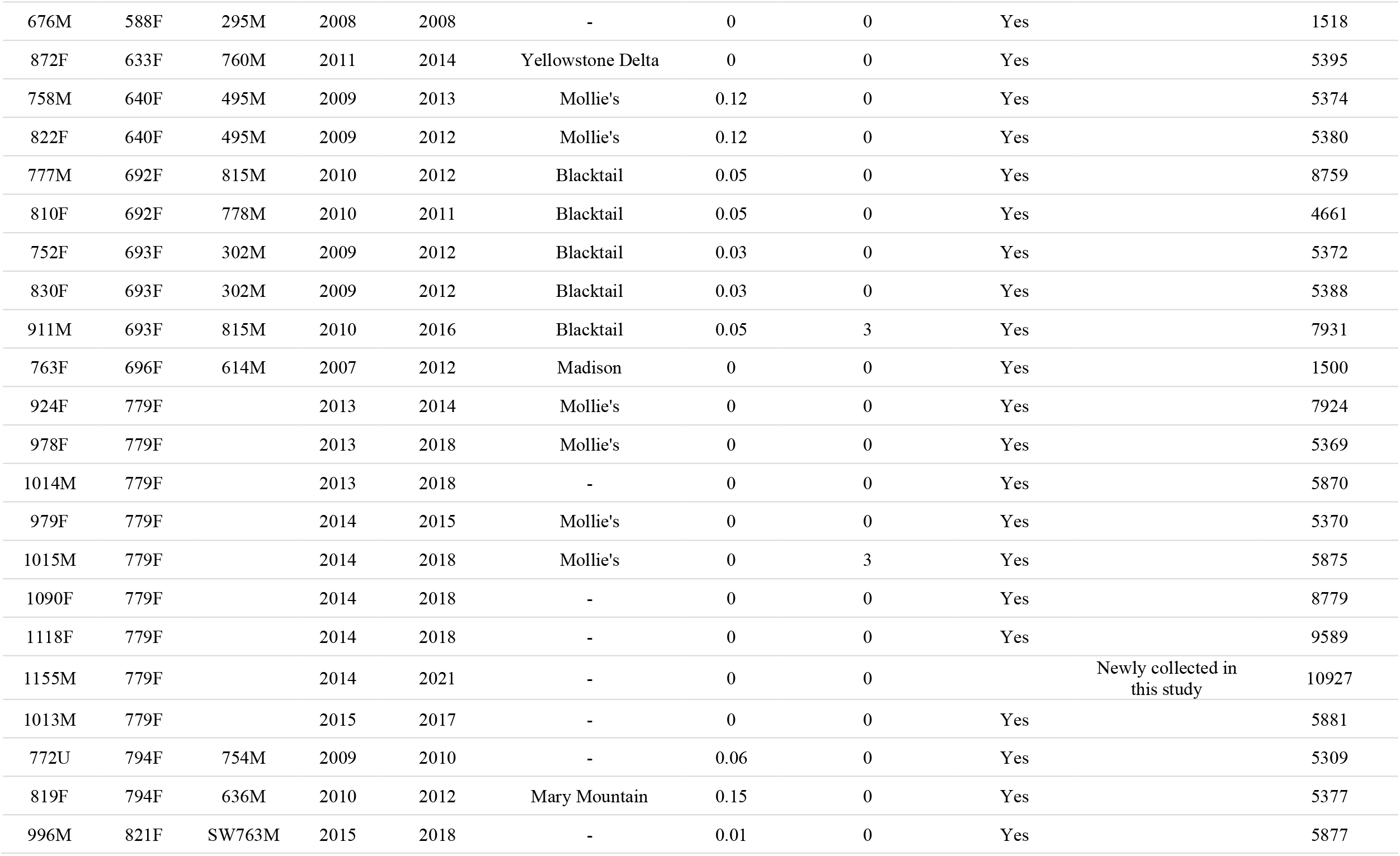
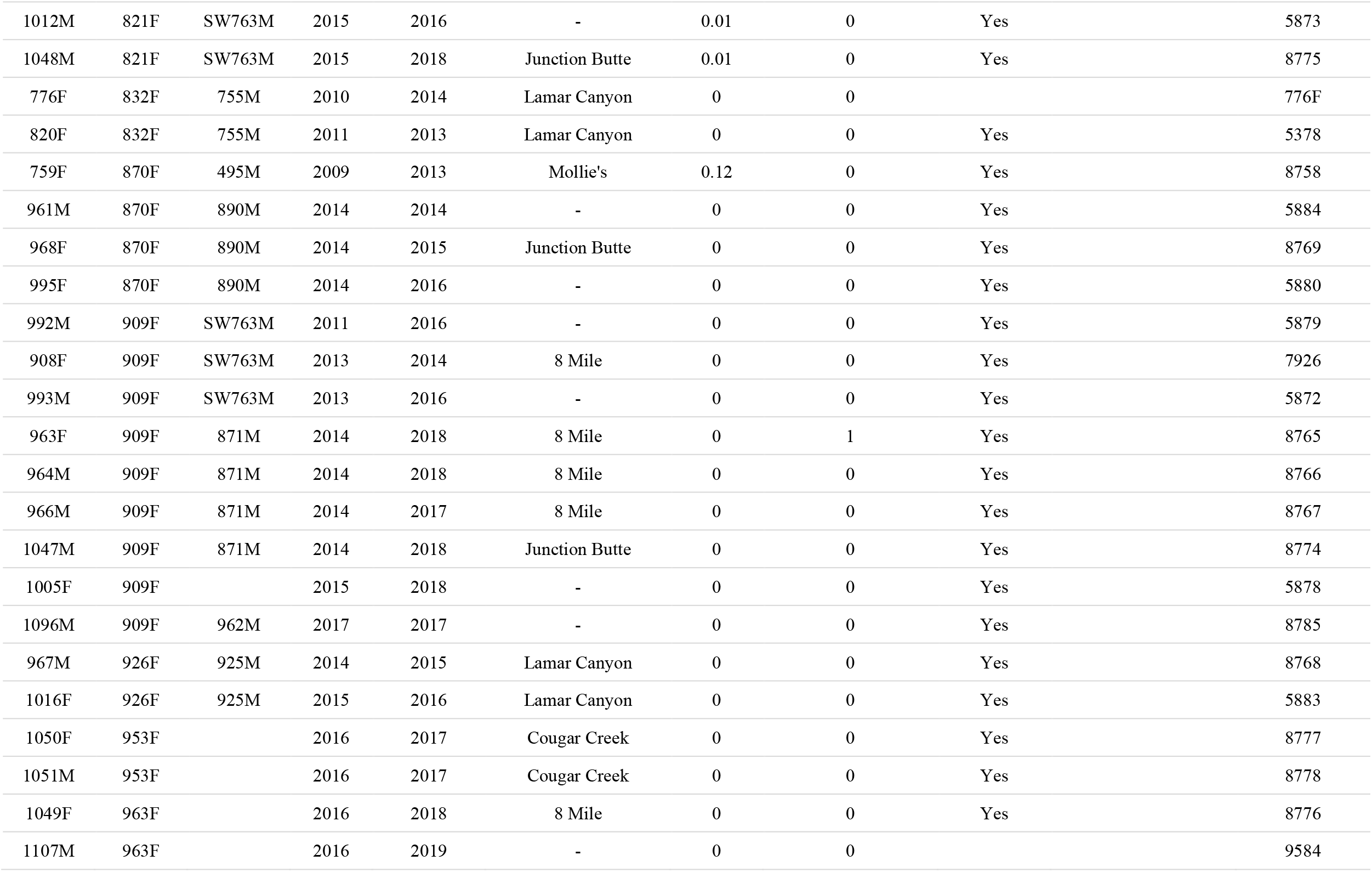
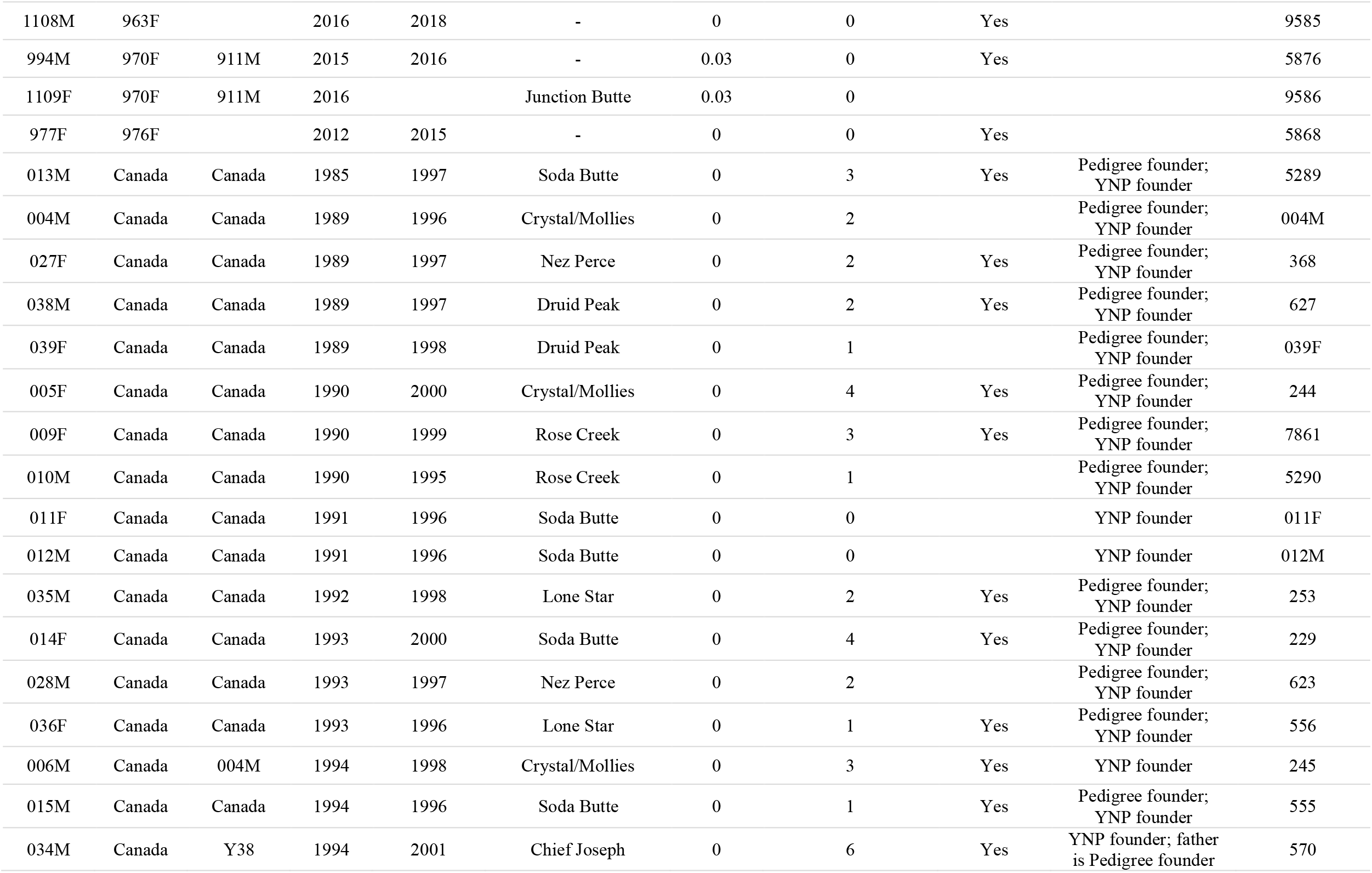
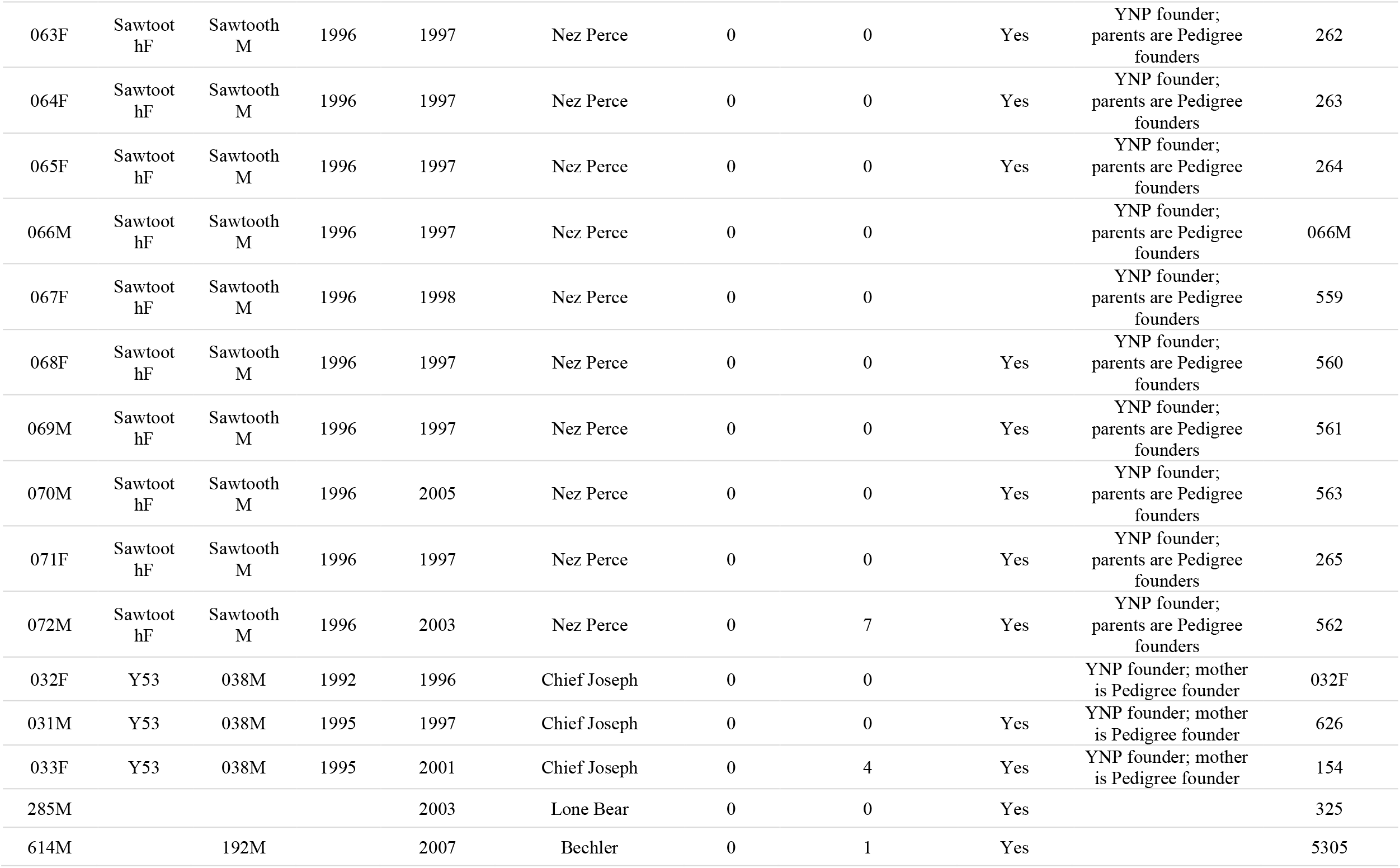
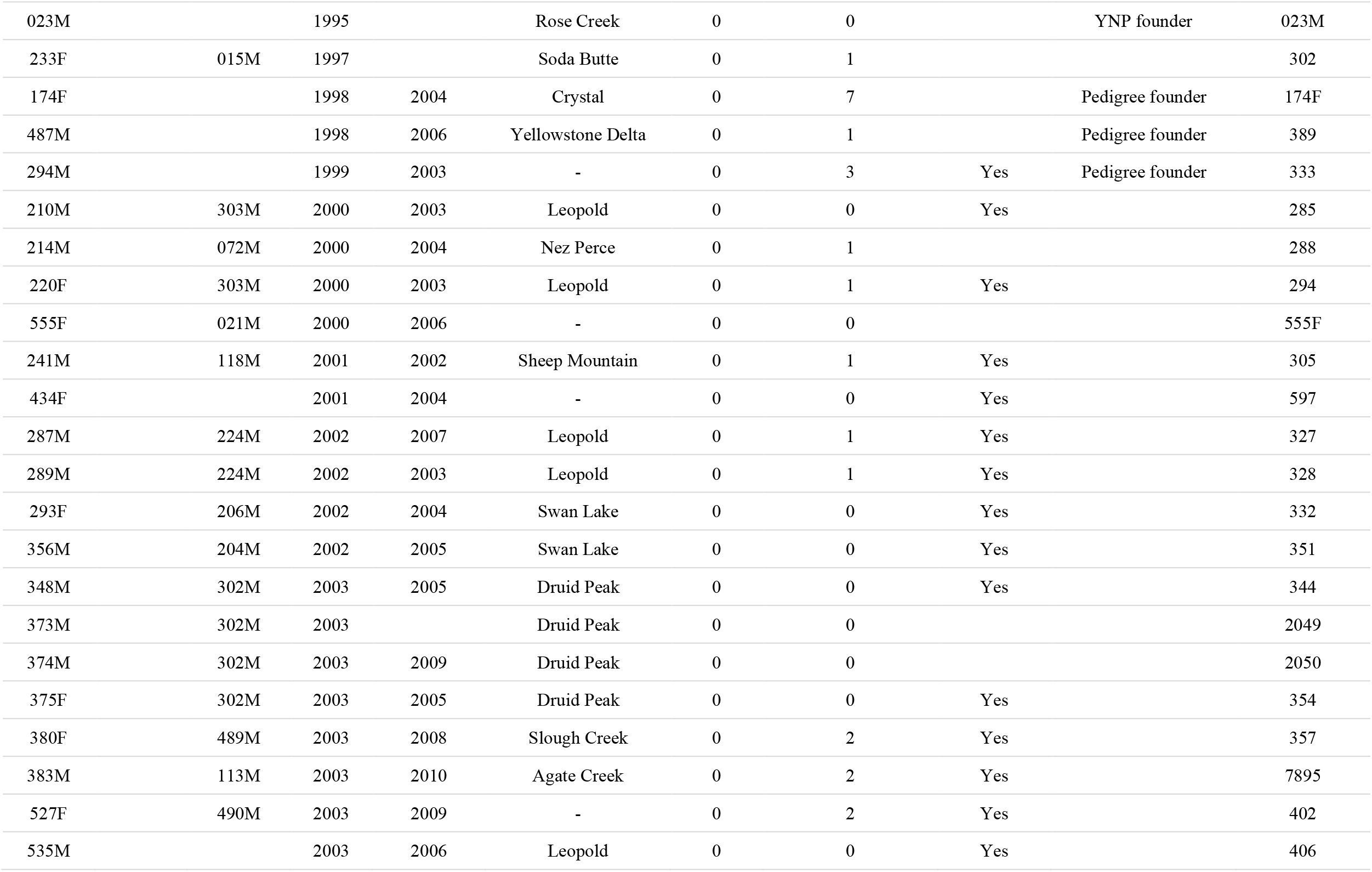
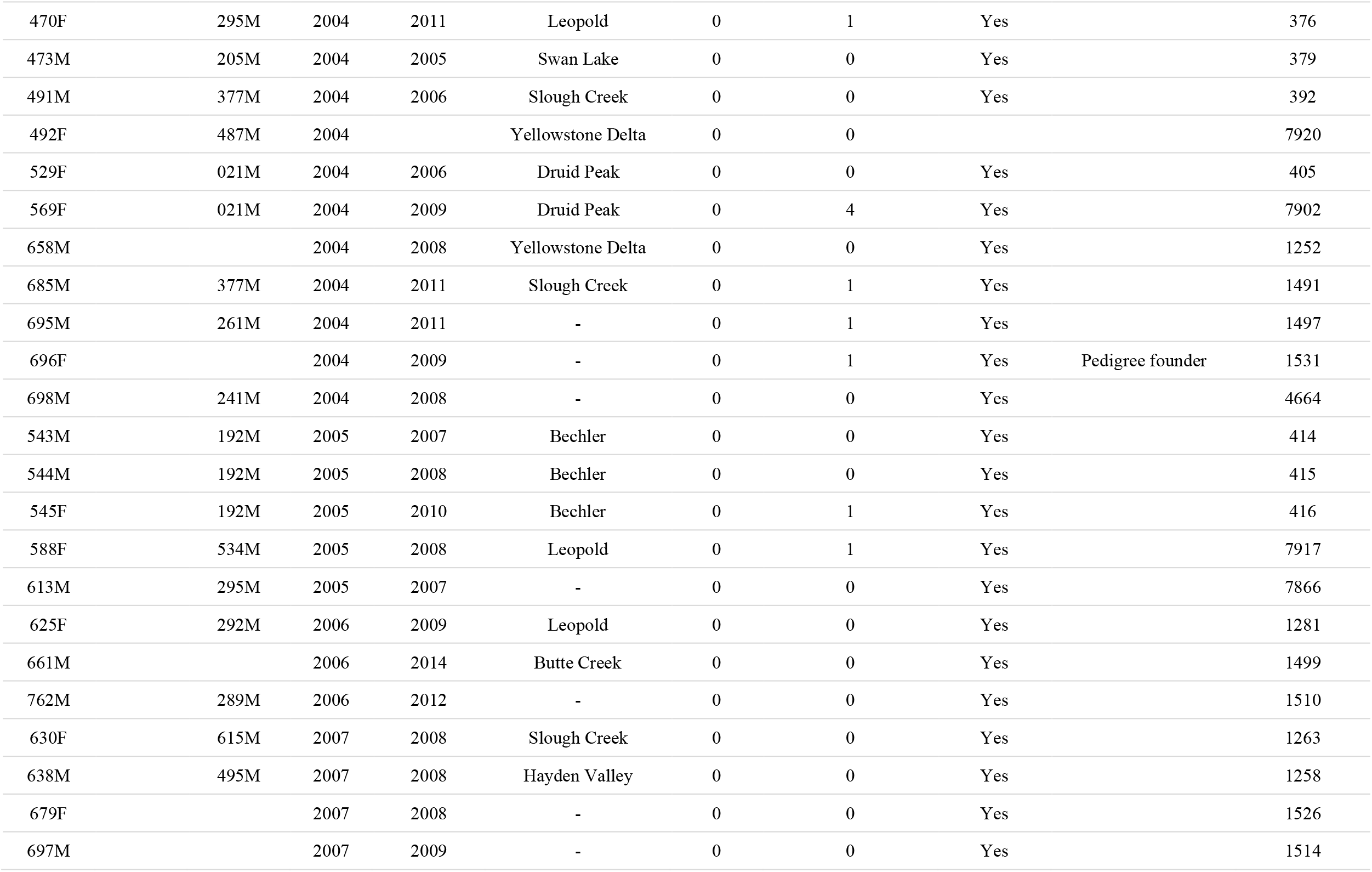
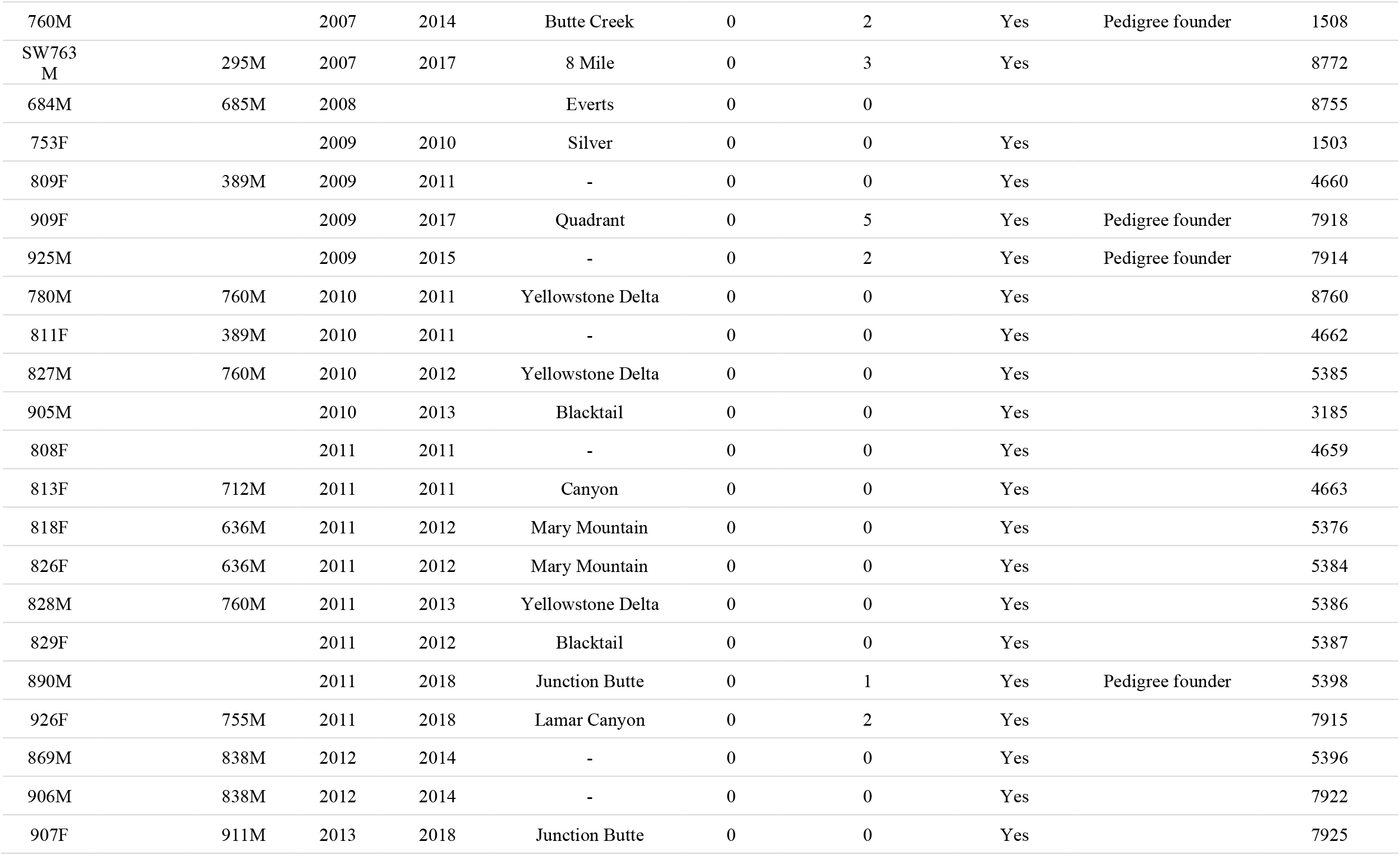
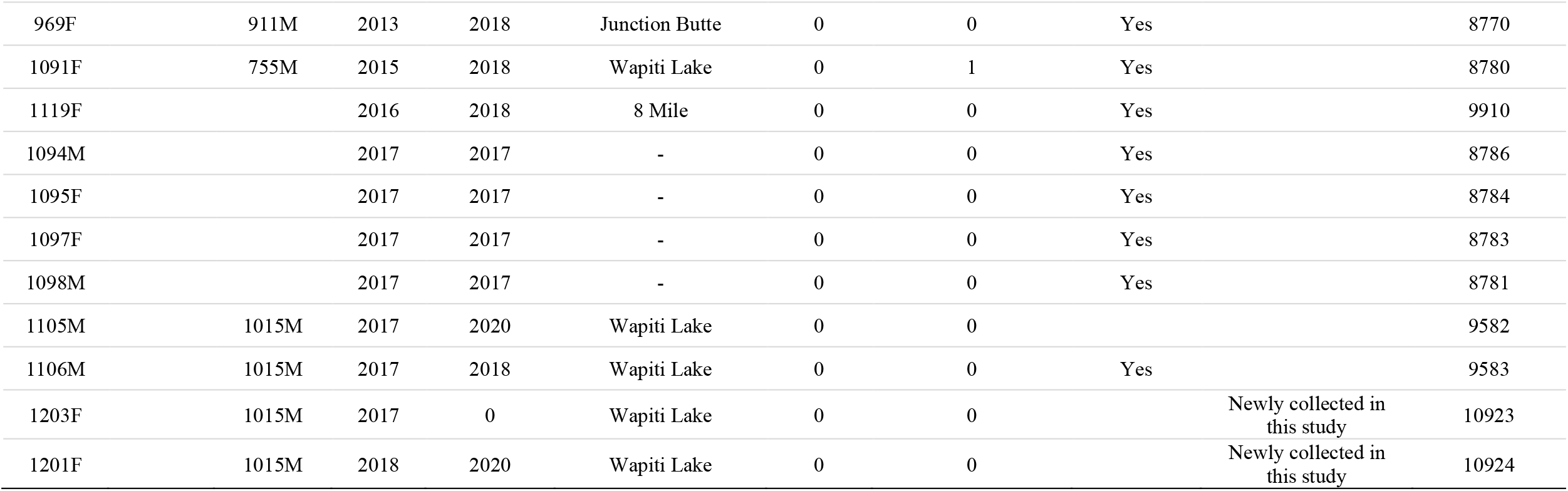
Sample information and meta-data for 474 Yellowstone gray wolves in pedigree analysis. Genetic or field-confirmed parentage as well as their natal pack name is provided. Individuals with genetic data included in this study are indicated in the “RADseq data” column. Individuals that founded the population are noted with one or both parents listed as “Canada” as per unknown pedigree information. (Abbreviations: F, female; M, male; YOB, year of birth; YOD, year of death)

**Supplemental Table S2.**
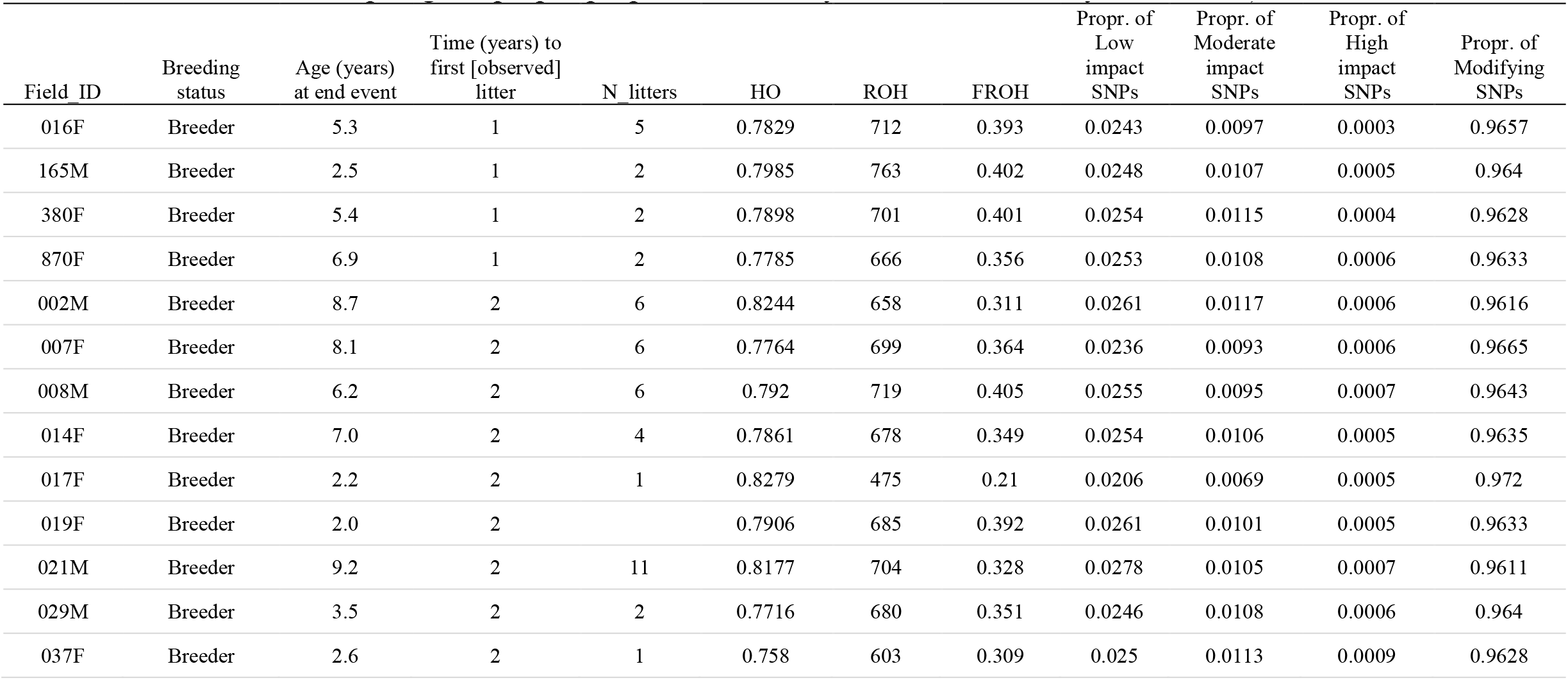
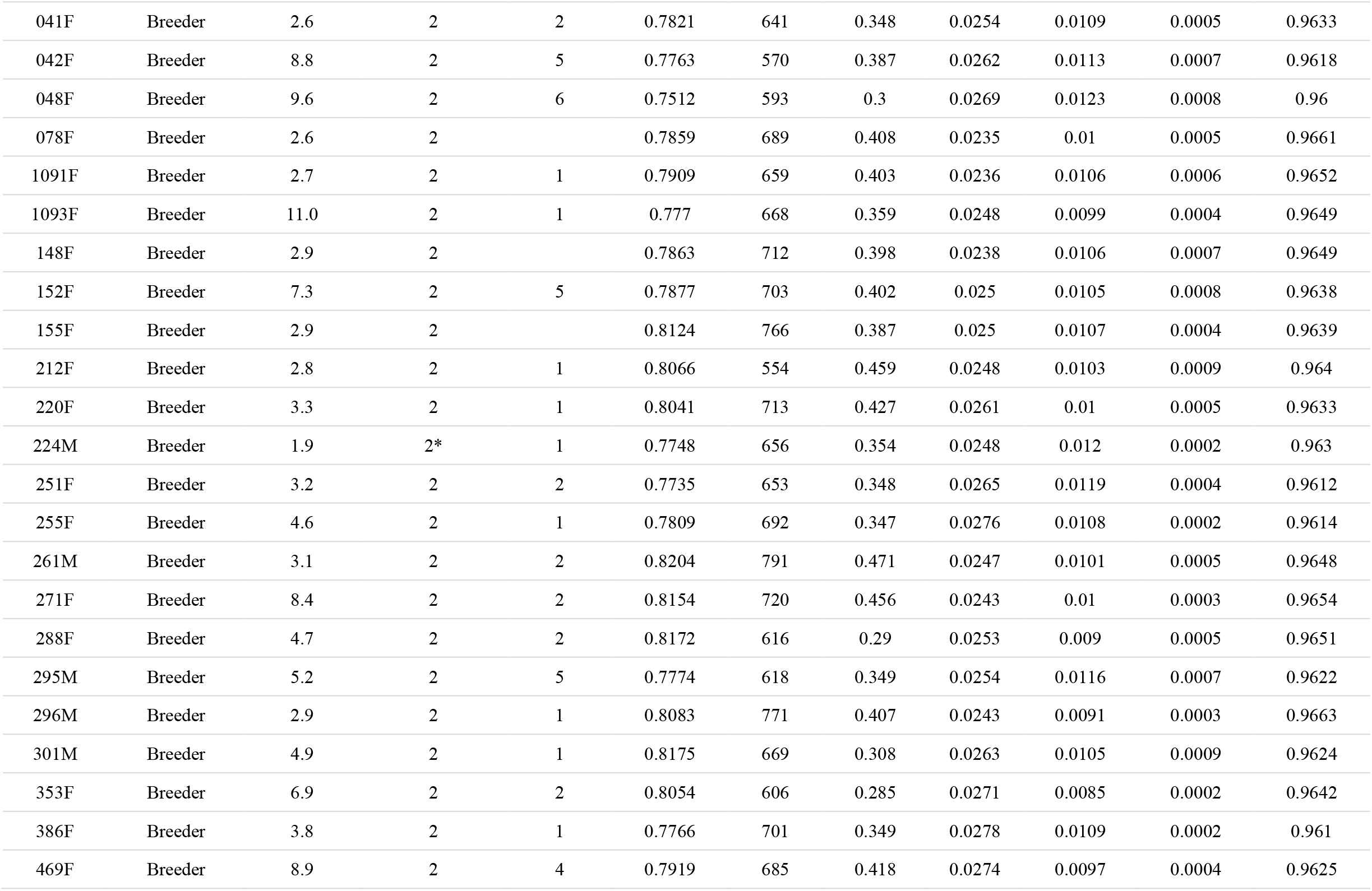
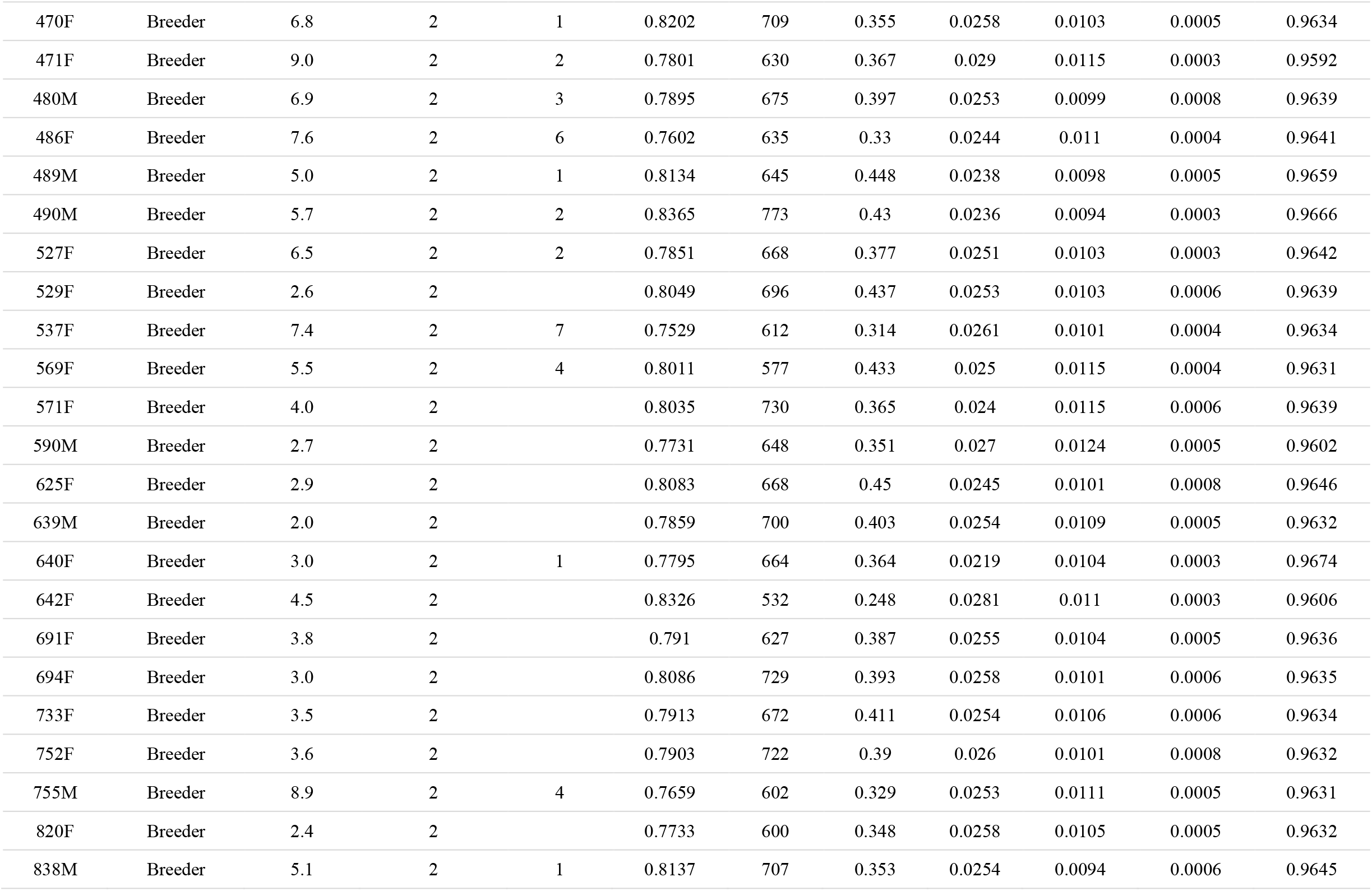
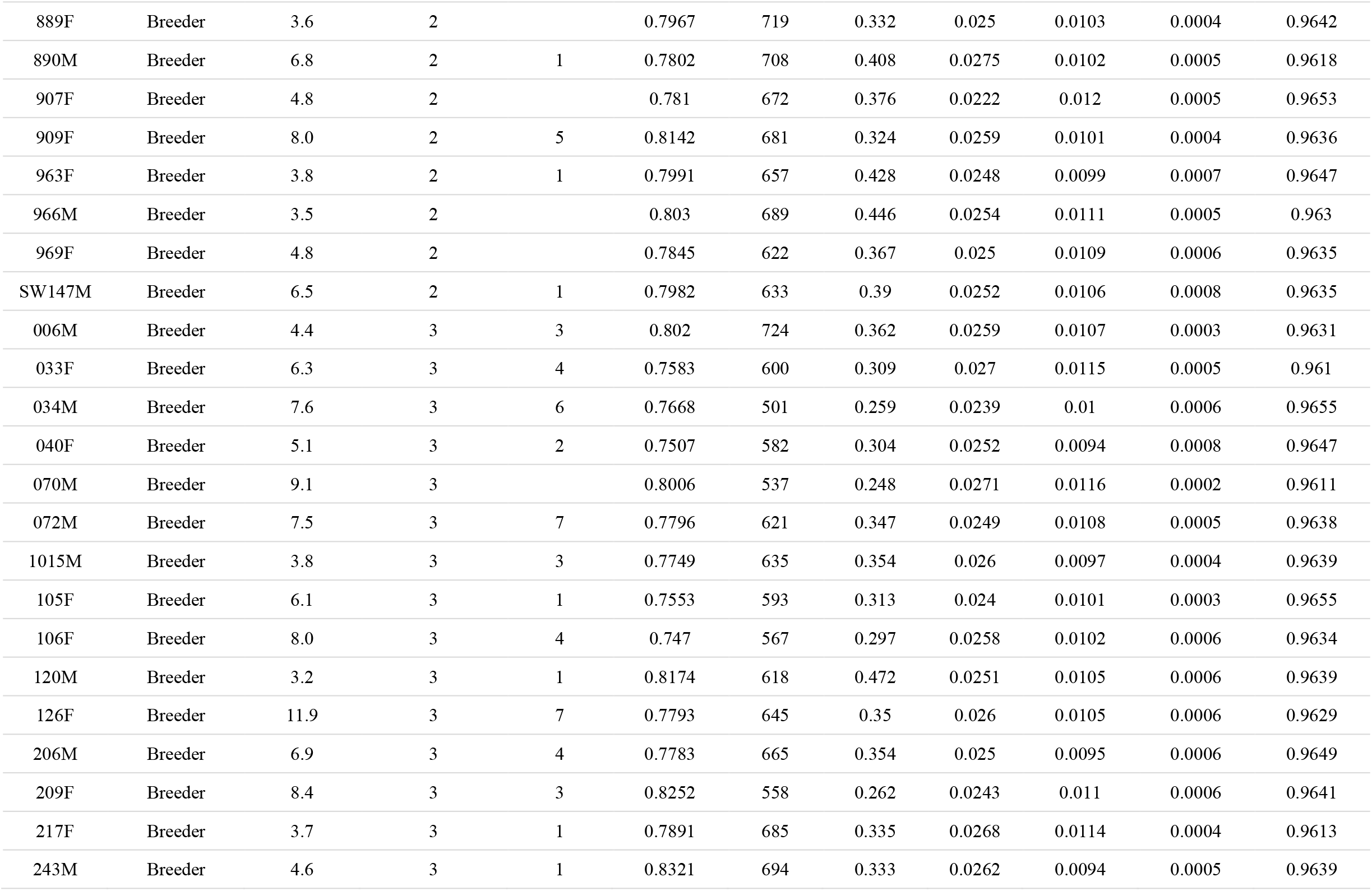
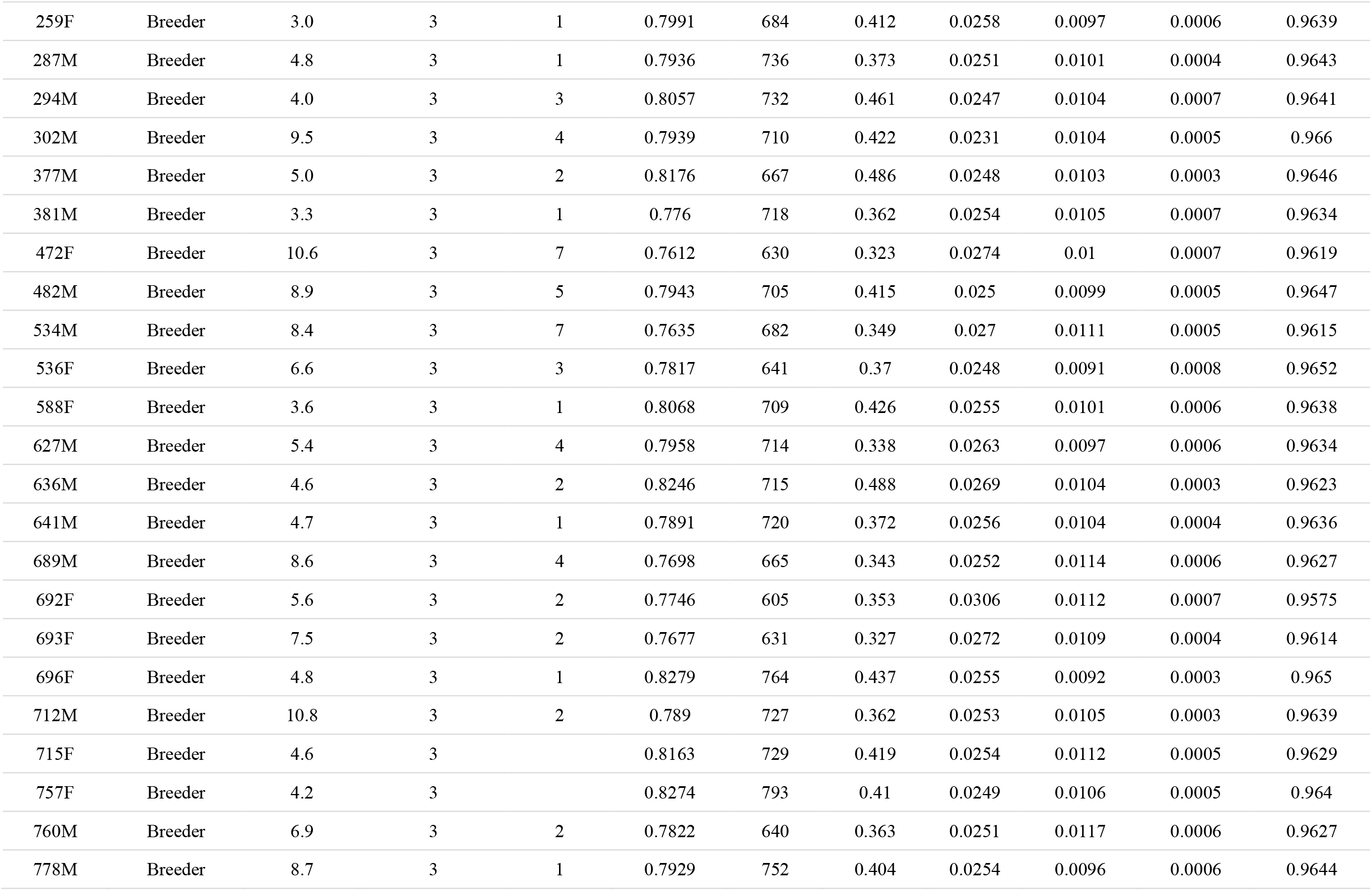
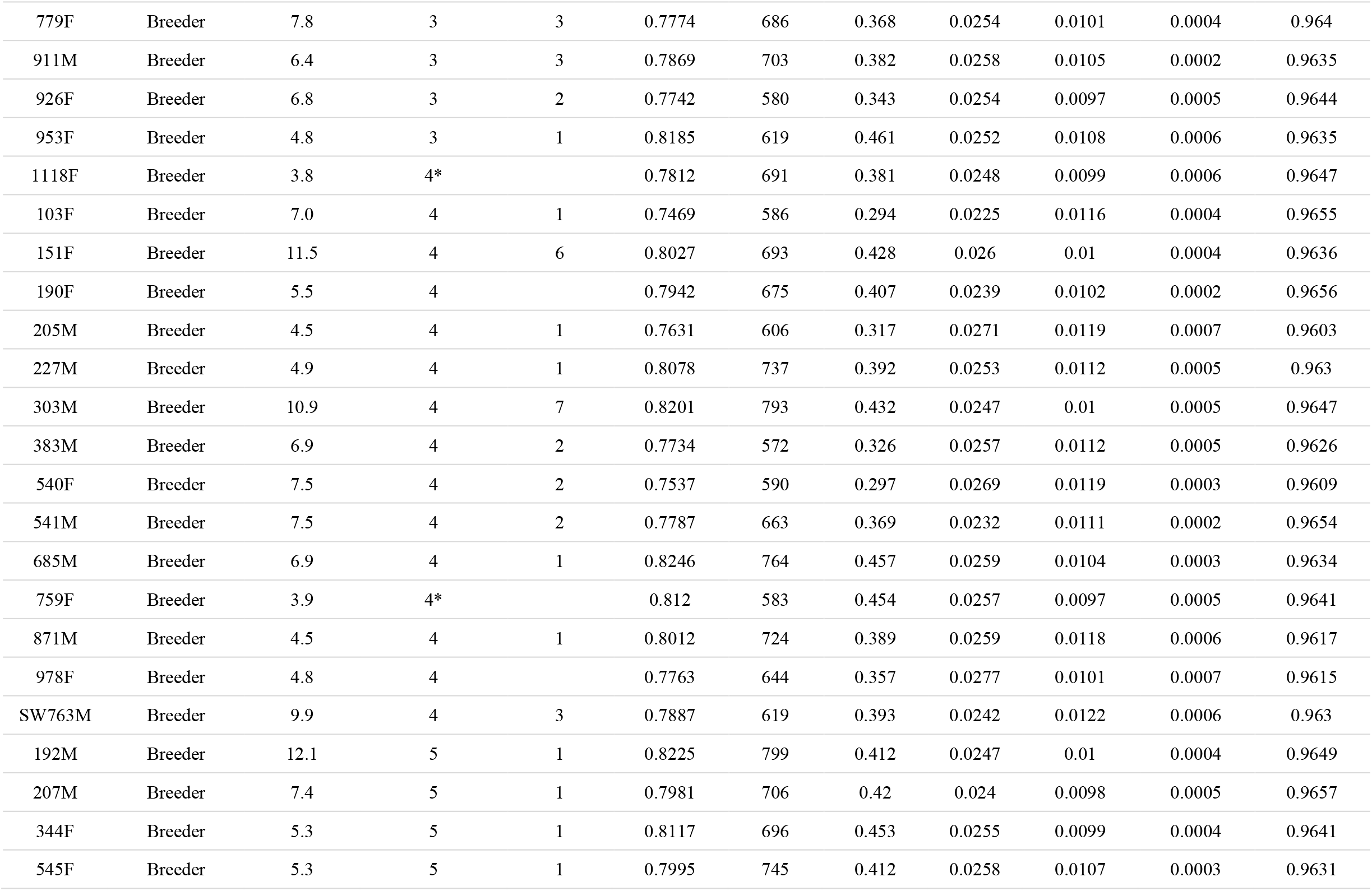
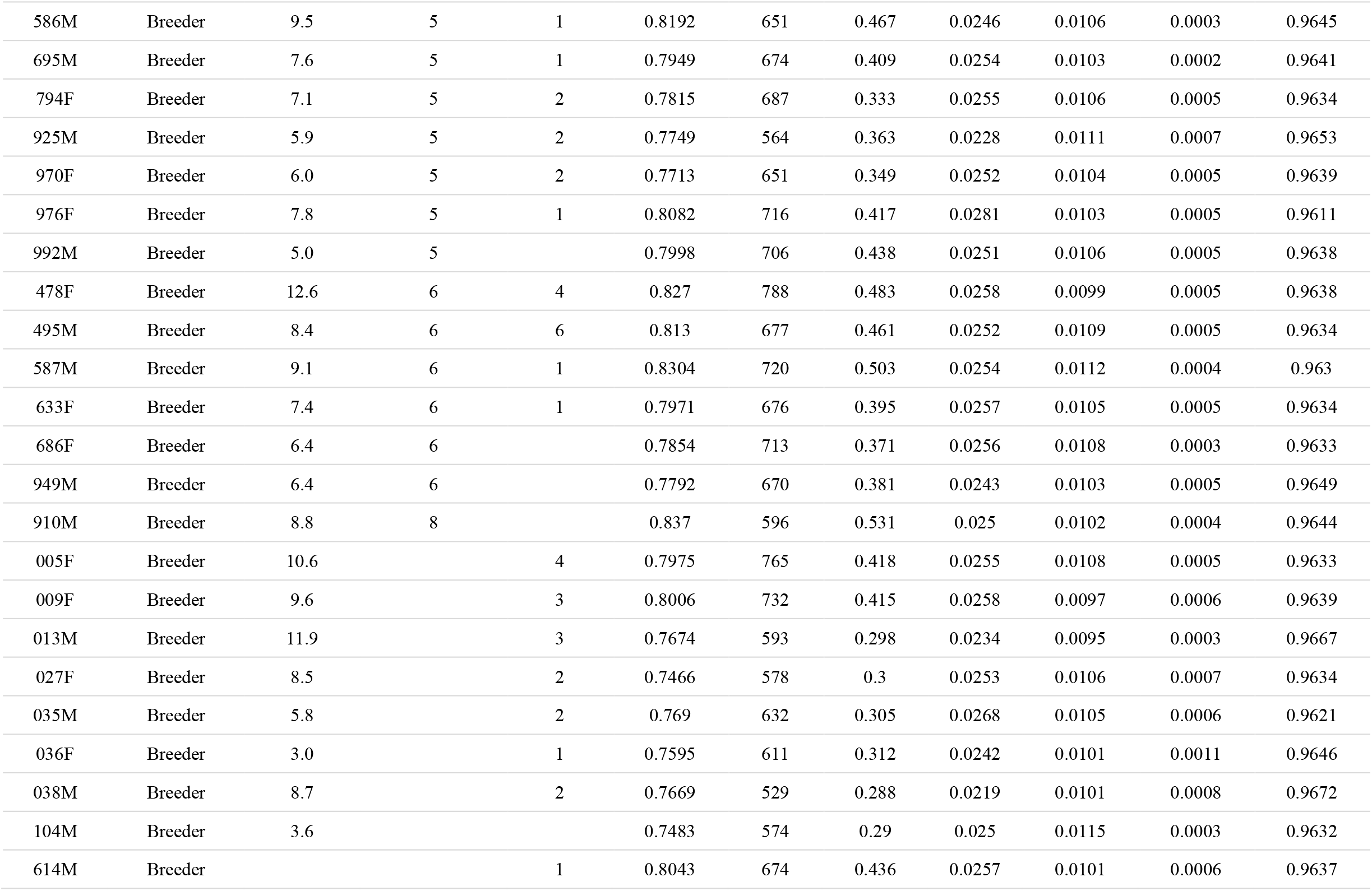
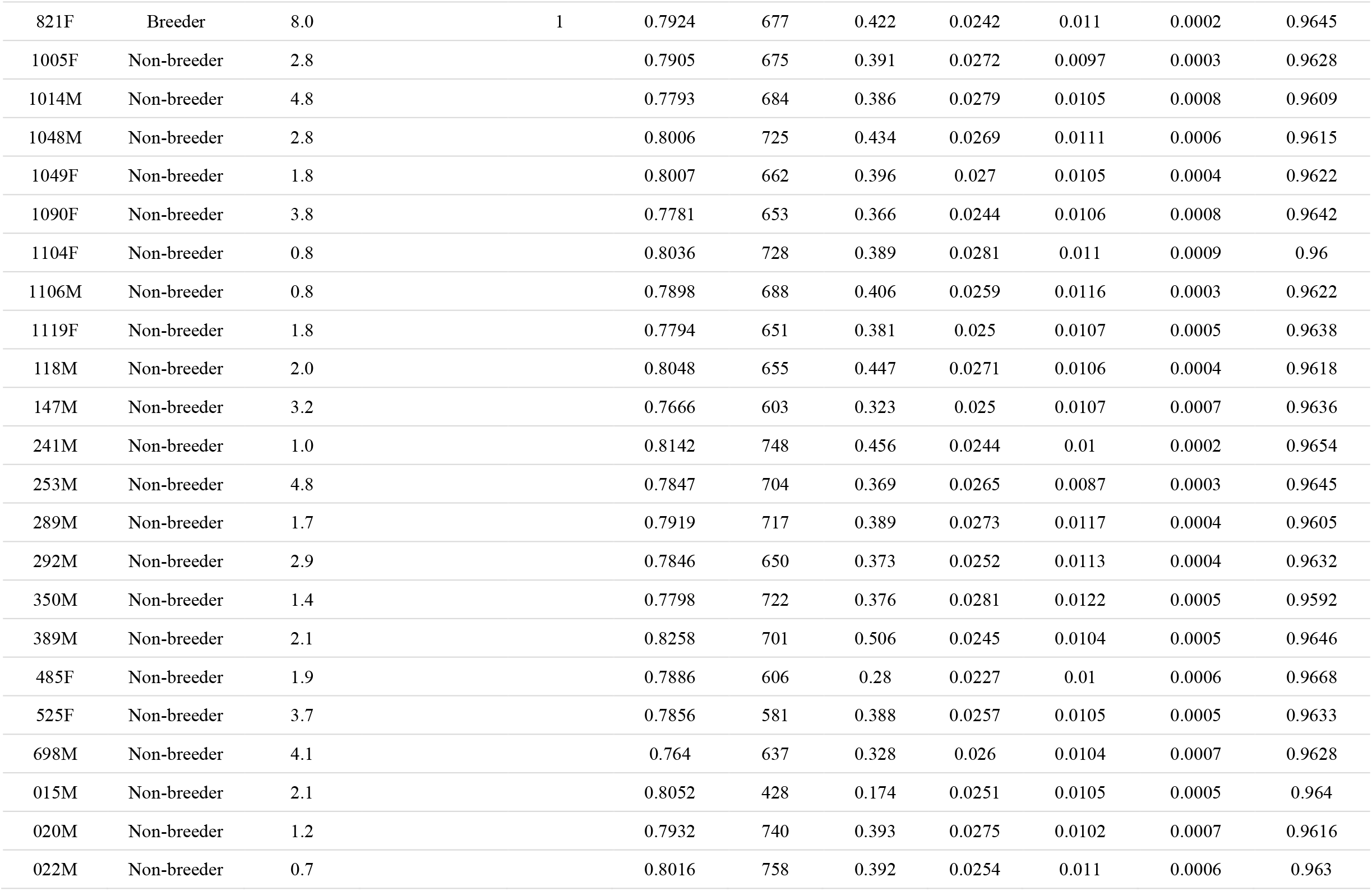
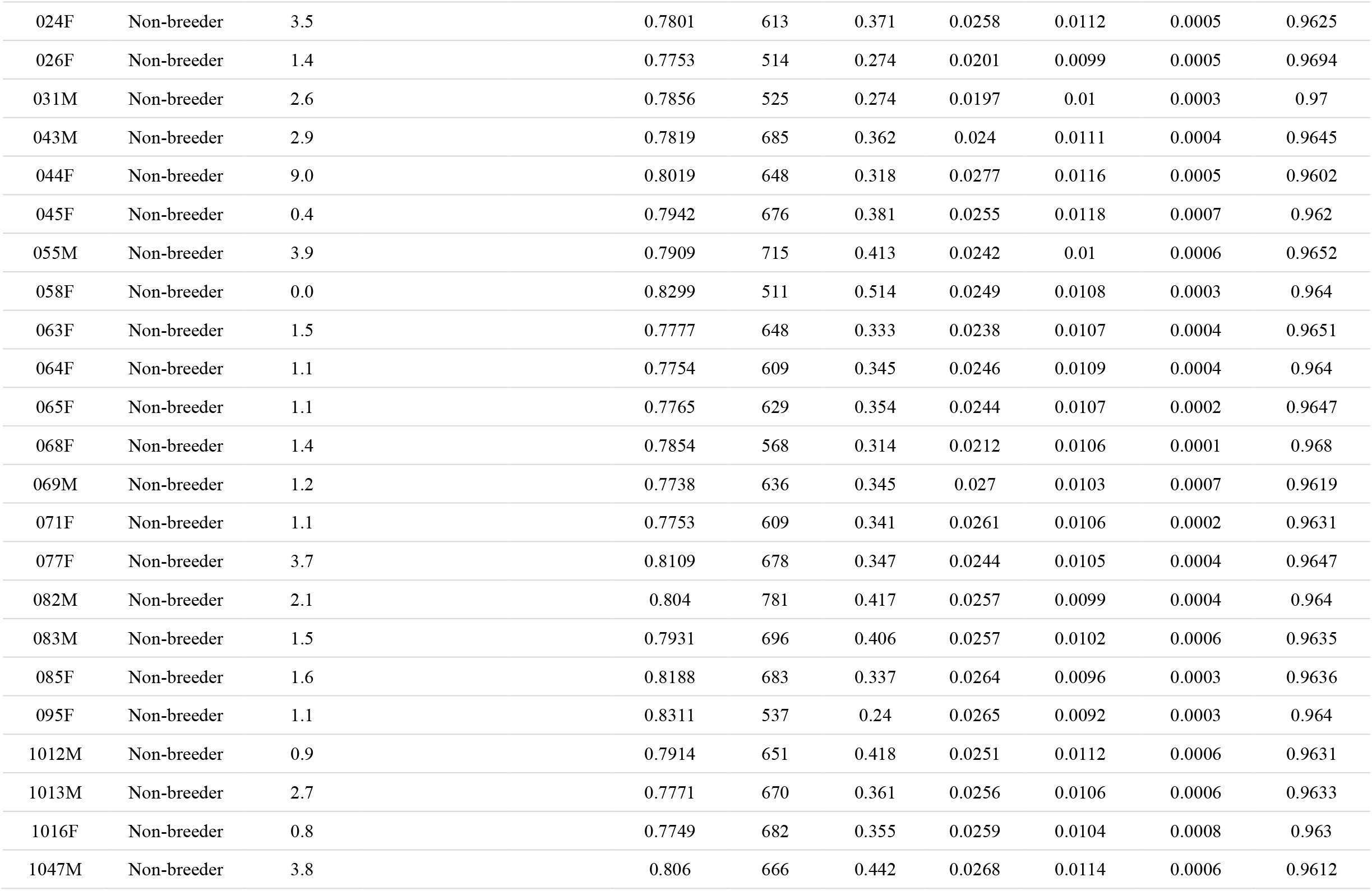
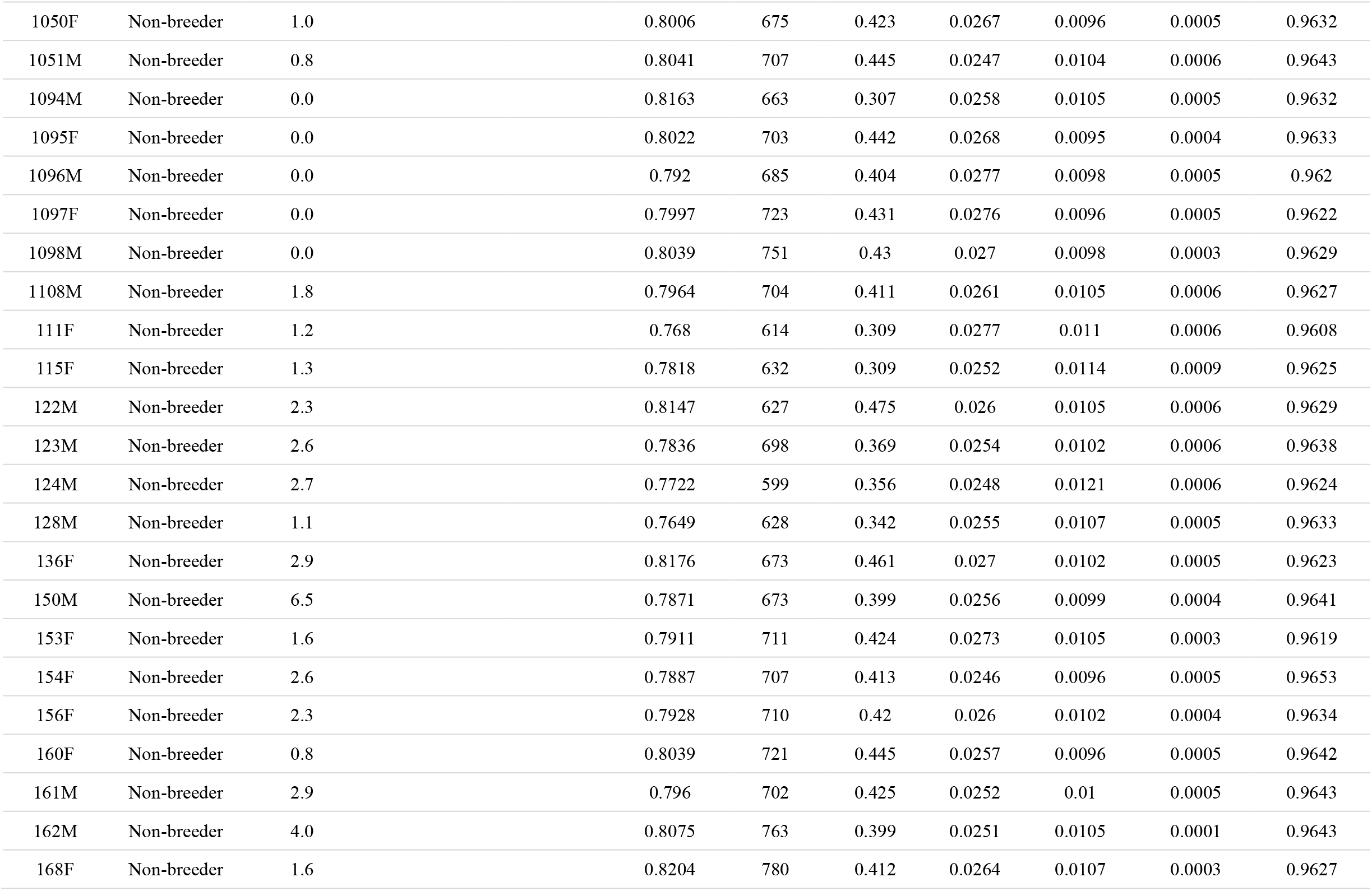
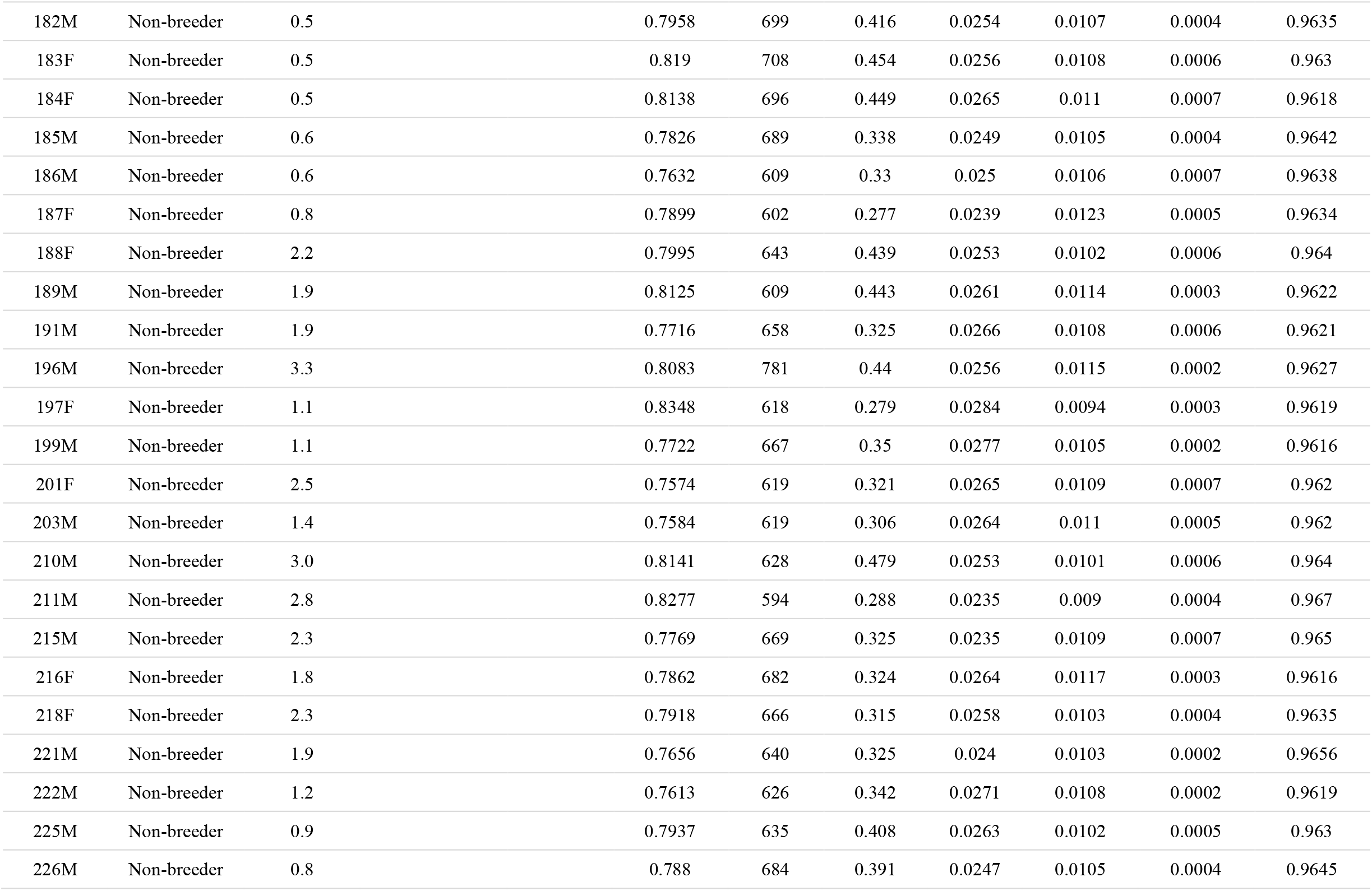
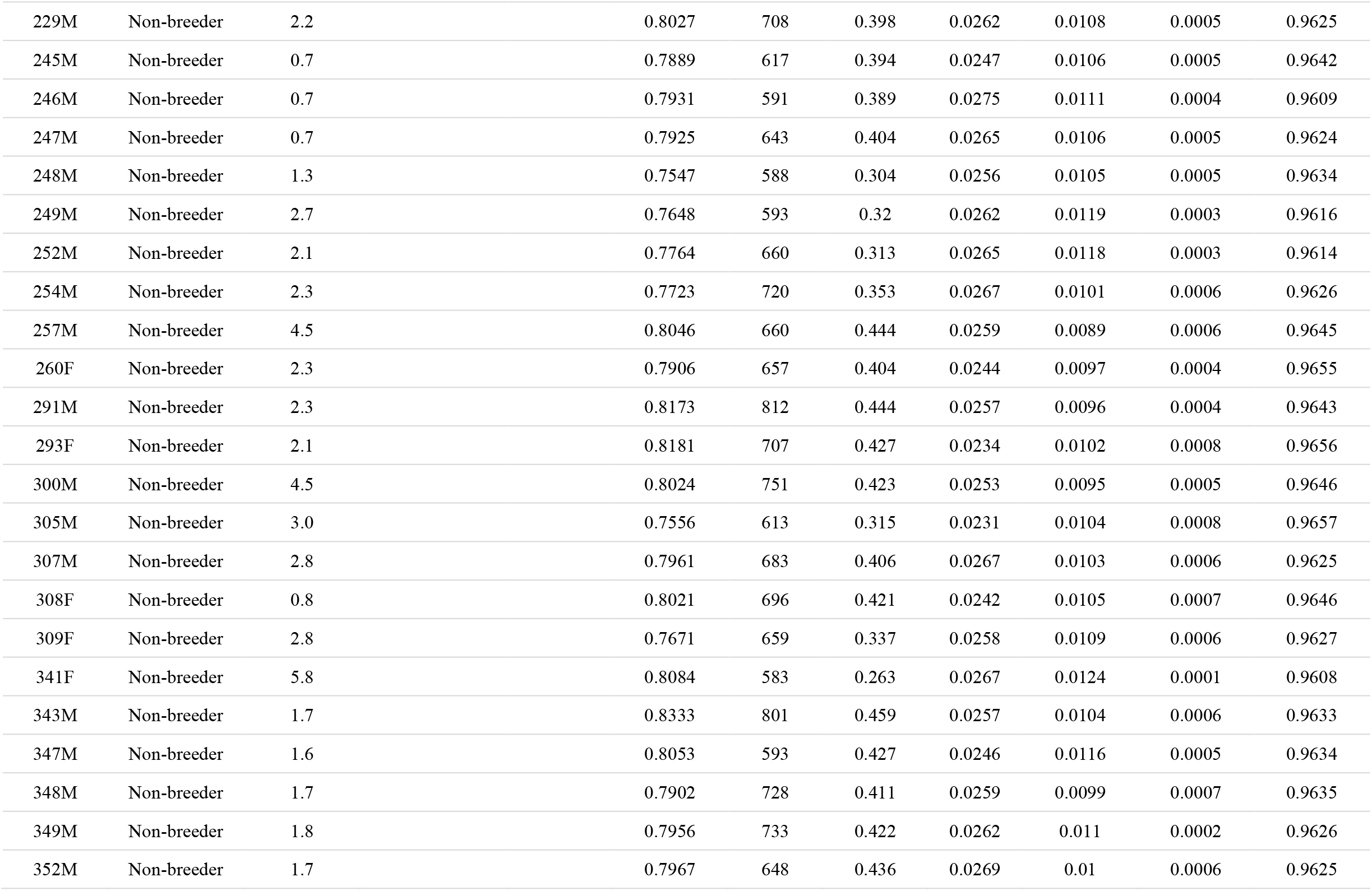
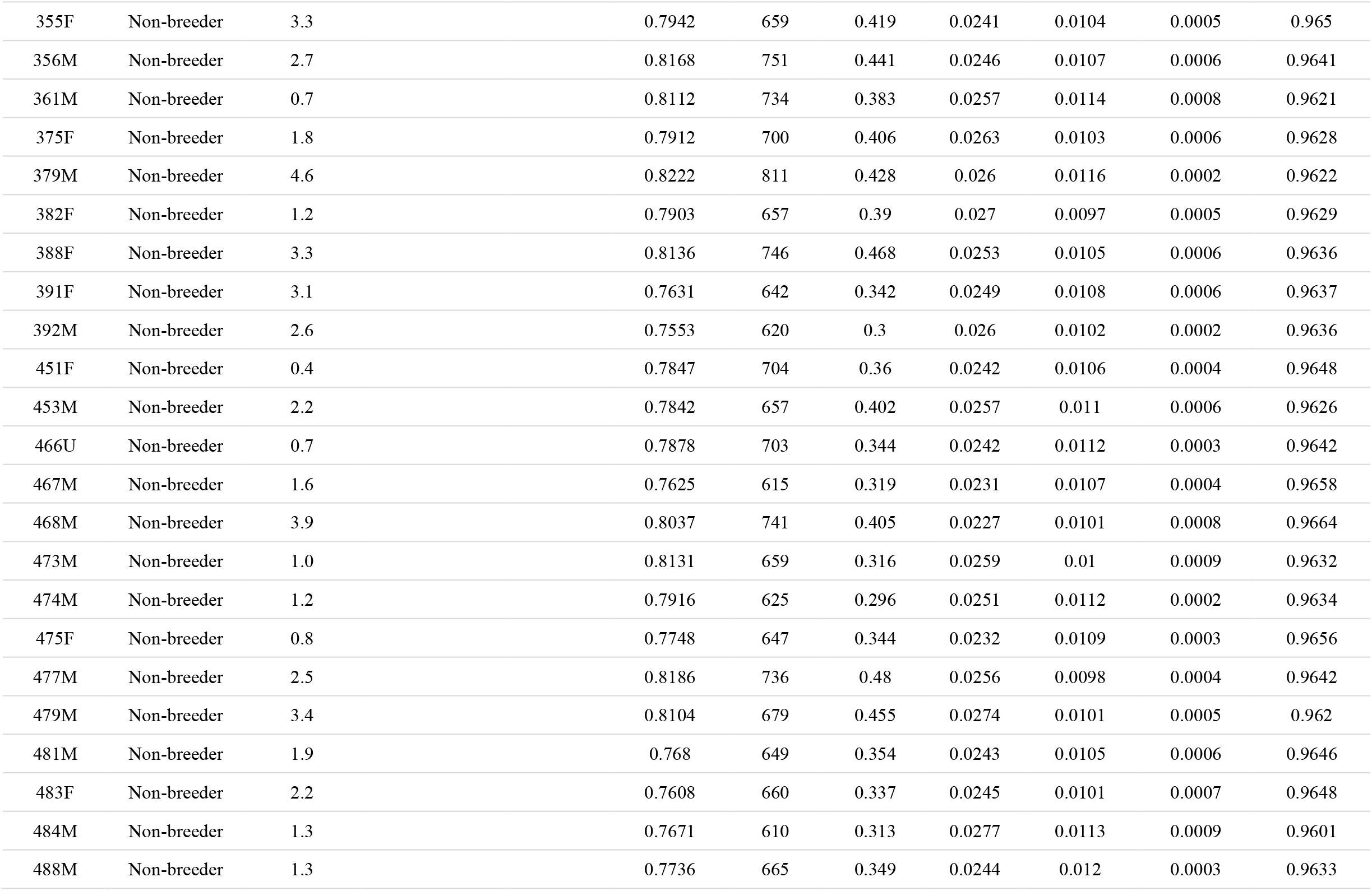
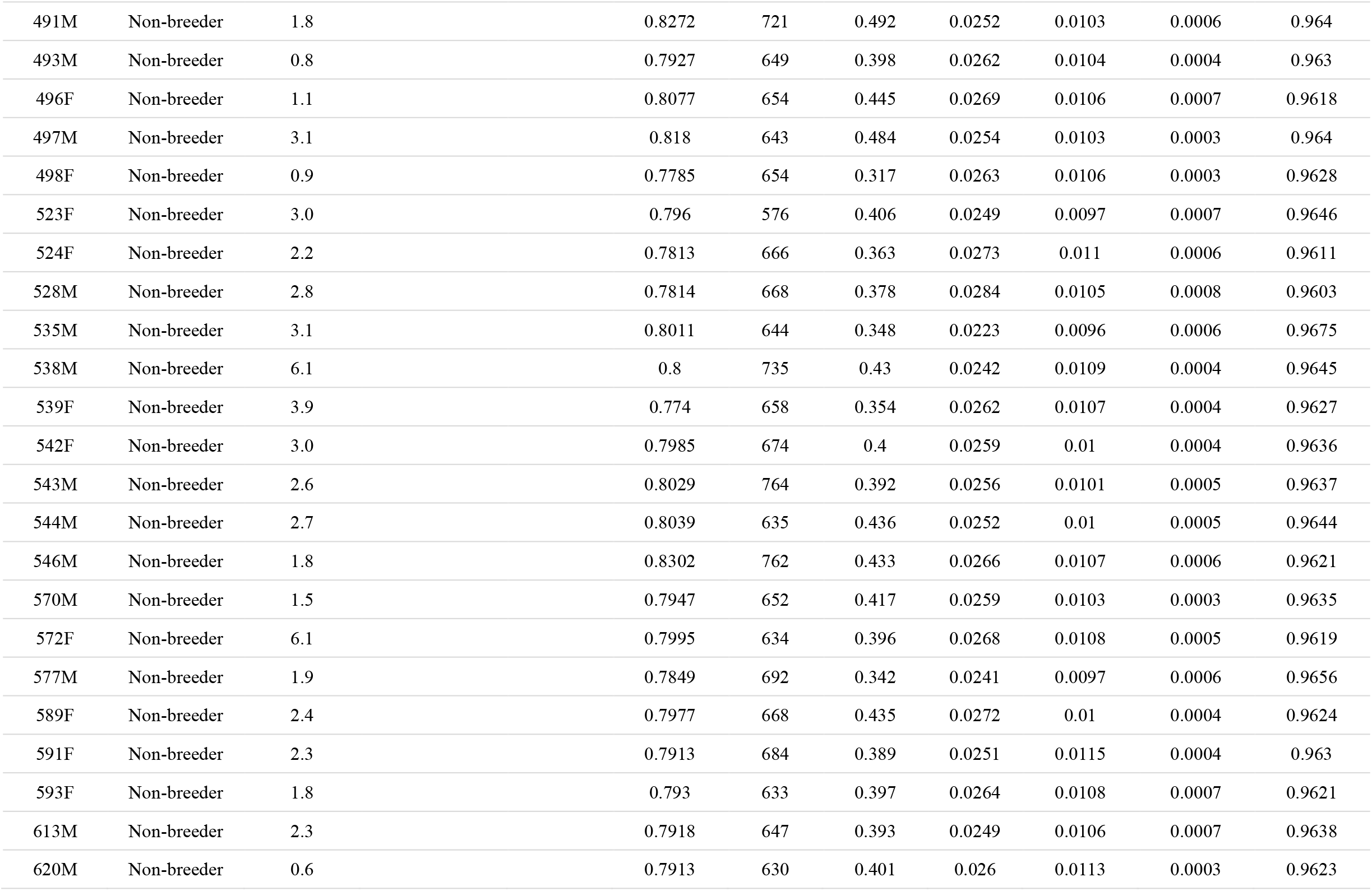
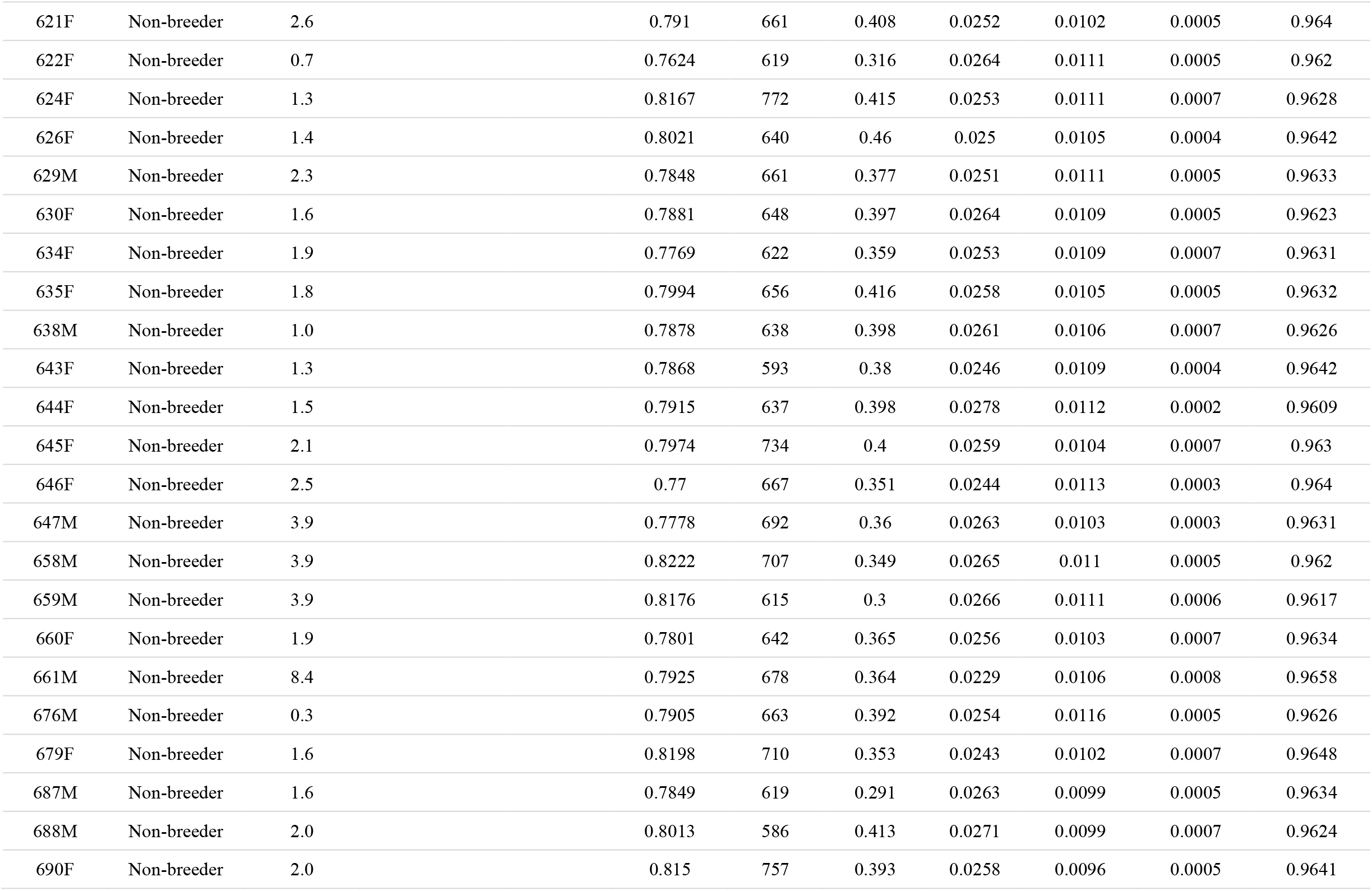
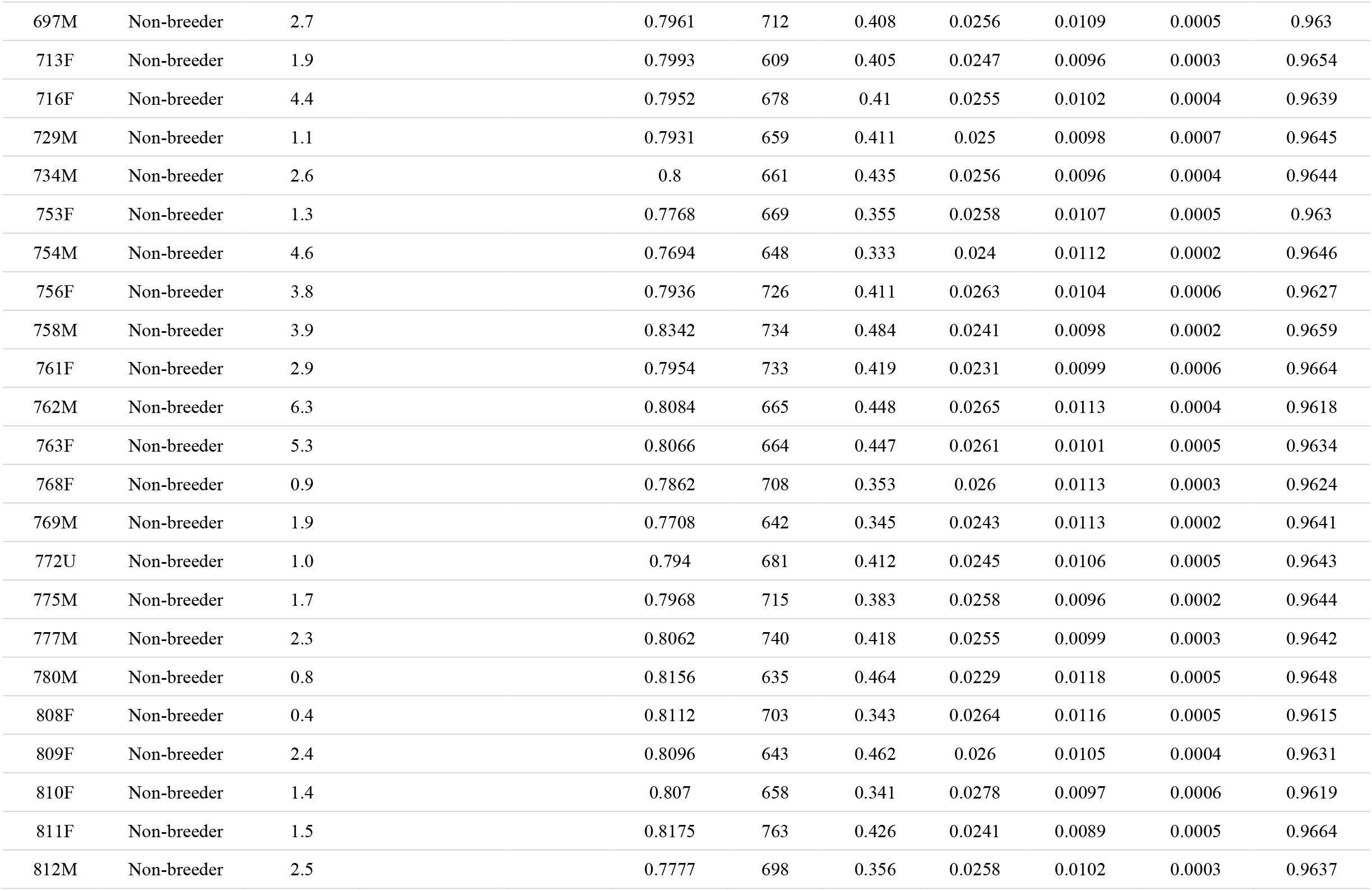
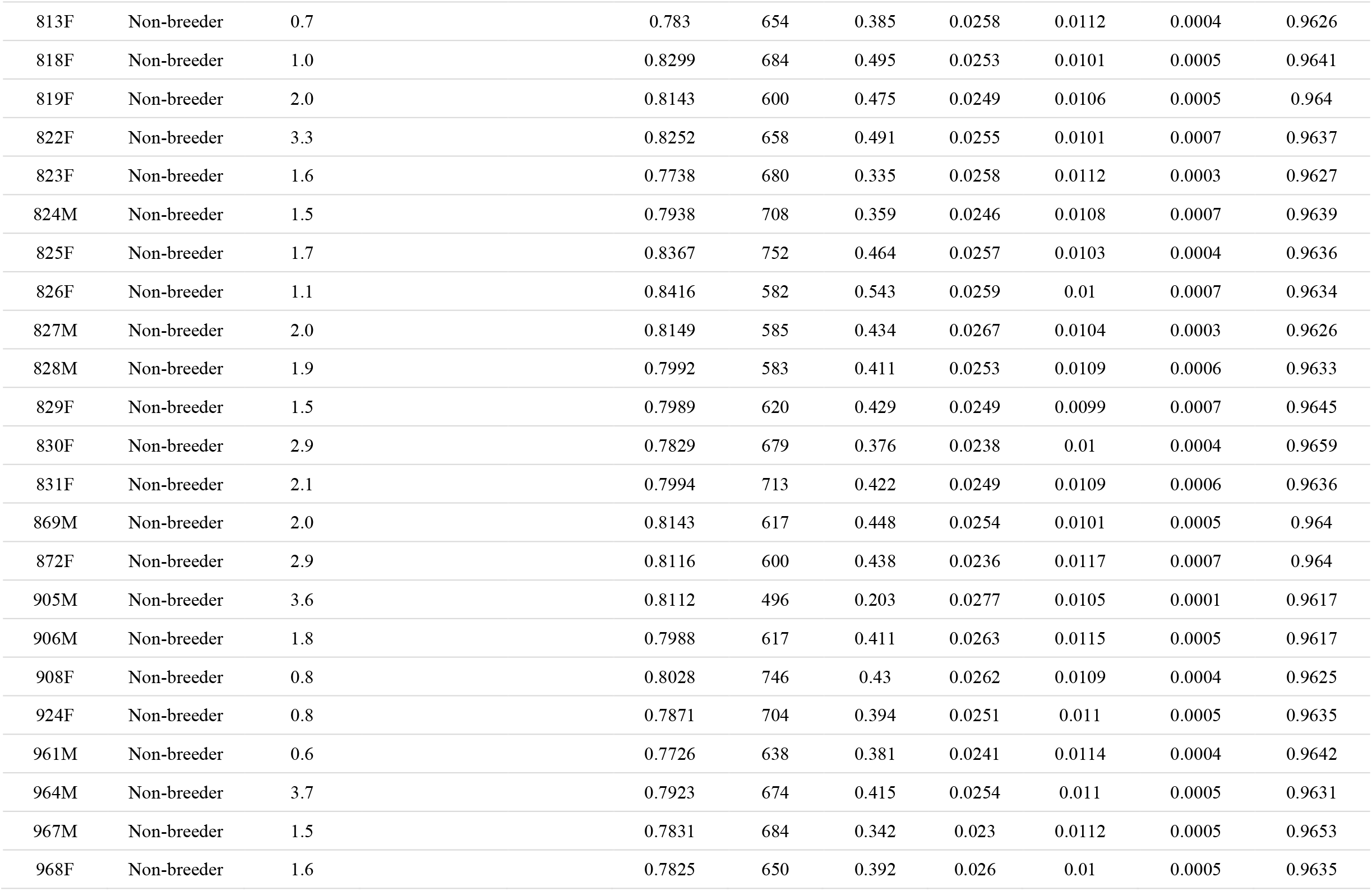
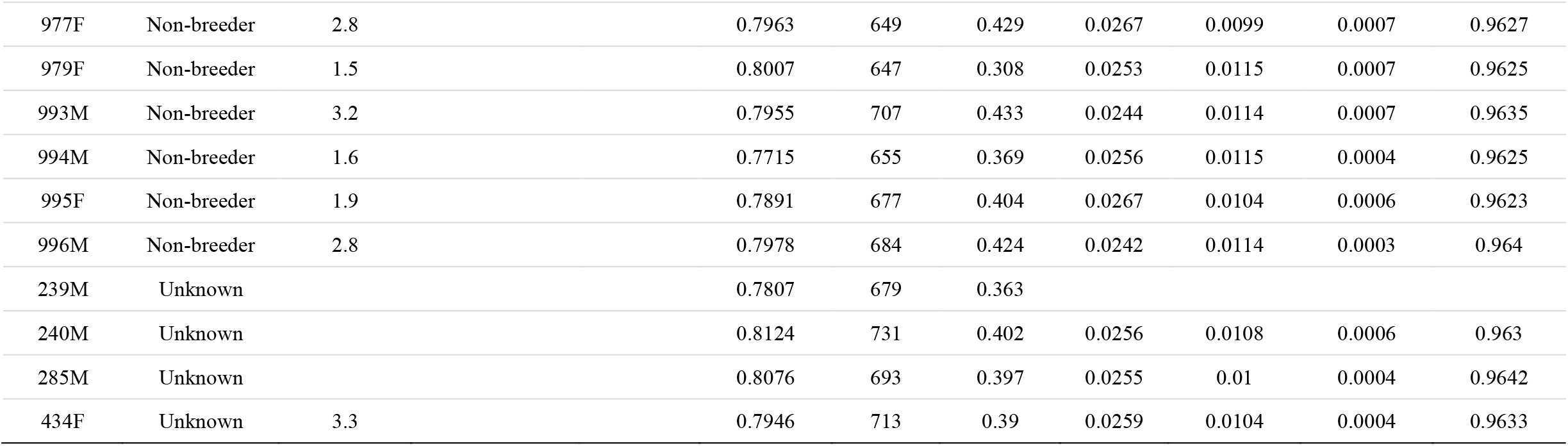
Sample information, meta-data, and descriptive values for each of the 391 Yellowstone National Park gray wolves in the genetic analysis. Reference to “end event” is either age at death or the last observation documented for that individual. Time in years to first observed litter; for animals that were translocated into YNP with offspring or reproduced after leaving YNP will lack an observed time of first litter and likely considered non-breeding individuals. All ROH estimates have at least 10 SNPs within the 10Kb tract and estimated from the 24K statistically unlinked set of SNPs. Asterisks indicate wolves with first observed litter dates older than age at end event that were excluded from the survival model analysis. (Abbreviations: *F_ROH_*; inbreeding coefficient estimated from runs of homozygosity; HO, observed heterozygosity; ROH, runs of homozygosity; N_litters, number of litters identified from the pedigree; propr., proportion; YOB, year of birth; YOD, year of death)

| Field_ID | Breeding status | Age (years) at end event | Time (years) to first [observed] litter | N_litters | HO | ROH | FROH | Propr. of Low impact SNPs | Propr. of Moderate impact SNPs | Propr. of High impact SNPs | Propr. of Modifying SNPs |
| --- | --- | --- | --- | --- | --- | --- | --- | --- | --- | --- | --- |
| 016F | Breeder | 5.3 | 1 | 5 | 0.7829 | 712 | 0.393 | 0.0243 | 0.0097 | 0.0003 | 0.9657 |
| 165M | Breeder | 2.5 | 1 | 2 | 0.7985 | 763 | 0.402 | 0.0248 | 0.0107 | 0.0005 | 0.964 |
| 380F | Breeder | 5.4 | 1 | 2 | 0.7898 | 701 | 0.401 | 0.0254 | 0.0115 | 0.0004 | 0.9628 |
| 870F | Breeder | 6.9 | 1 | 2 | 0.7785 | 666 | 0.356 | 0.0253 | 0.0108 | 0.0006 | 0.9633 |
| 002M | Breeder | 8.7 | 2 | 6 | 0.8244 | 658 | 0.311 | 0.0261 | 0.0117 | 0.0006 | 0.9616 |
| 007F | Breeder | 8.1 | 2 | 6 | 0.7764 | 699 | 0.364 | 0.0236 | 0.0093 | 0.0006 | 0.9665 |
| 008M | Breeder | 6.2 | 2 | 6 | 0.792 | 719 | 0.405 | 0.0255 | 0.0095 | 0.0007 | 0.9643 |
| 014F | Breeder | 7.0 | 2 | 4 | 0.7861 | 678 | 0.349 | 0.0254 | 0.0106 | 0.0005 | 0.9635 |
| 017F | Breeder | 2.2 | 2 | 1 | 0.8279 | 475 | 0.21 | 0.0206 | 0.0069 | 0.0005 | 0.972 |
| 019F | Breeder | 2.0 | 2 |  | 0.7906 | 685 | 0.392 | 0.0261 | 0.0101 | 0.0005 | 0.9633 |
| 021M | Breeder | 9.2 | 2 | 11 | 0.8177 | 704 | 0.328 | 0.0278 | 0.0105 | 0.0007 | 0.9611 |
| 029M | Breeder | 3.5 | 2 | 2 | 0.7716 | 680 | 0.351 | 0.0246 | 0.0108 | 0.0006 | 0.964 |
| 037F | Breeder | 2.6 | 2 | 1 | 0.758 | 603 | 0.309 | 0.025 | 0.0113 | 0.0009 | 0.9628 |
| 041F | Breeder | 2.6 | 2 | 2 | 0.7821 | 641 | 0.348 | 0.0254 | 0.0109 | 0.0005 | 0.9633 |
| 042F | Breeder | 8.8 | 2 | 5 | 0.7763 | 570 | 0.387 | 0.0262 | 0.0113 | 0.0007 | 0.9618 |
| 048F | Breeder | 9.6 | 2 | 6 | 0.7512 | 593 | 0.3 | 0.0269 | 0.0123 | 0.0008 | 0.96 |
| 078F | Breeder | 2.6 | 2 |  | 0.7859 | 689 | 0.408 | 0.0235 | 0.01 | 0.0005 | 0.9661 |
| 1091F | Breeder | 2.7 | 2 | 1 | 0.7909 | 659 | 0.403 | 0.0236 | 0.0106 | 0.0006 | 0.9652 |
| 1093F | Breeder | 11.0 | 2 | 1 | 0.777 | 668 | 0.359 | 0.0248 | 0.0099 | 0.0004 | 0.9649 |
| 148F | Breeder | 2.9 | 2 |  | 0.7863 | 712 | 0.398 | 0.0238 | 0.0106 | 0.0007 | 0.9649 |
| 152F | Breeder | 7.3 | 2 | 5 | 0.7877 | 703 | 0.402 | 0.025 | 0.0105 | 0.0008 | 0.9638 |
| 155F | Breeder | 2.9 | 2 |  | 0.8124 | 766 | 0.387 | 0.025 | 0.0107 | 0.0004 | 0.9639 |
| 212F | Breeder | 2.8 | 2 | 1 | 0.8066 | 554 | 0.459 | 0.0248 | 0.0103 | 0.0009 | 0.964 |
| 220F | Breeder | 3.3 | 2 | 1 | 0.8041 | 713 | 0.427 | 0.0261 | 0.01 | 0.0005 | 0.9633 |
| 224M | Breeder | 1.9 | 2* | 1 | 0.7748 | 656 | 0.354 | 0.0248 | 0.012 | 0.0002 | 0.963 |
| 251F | Breeder | 3.2 | 2 | 2 | 0.7735 | 653 | 0.348 | 0.0265 | 0.0119 | 0.0004 | 0.9612 |
| 255F | Breeder | 4.6 | 2 | 1 | 0.7809 | 692 | 0.347 | 0.0276 | 0.0108 | 0.0002 | 0.9614 |
| 261M | Breeder | 3.1 | 2 | 2 | 0.8204 | 791 | 0.471 | 0.0247 | 0.0101 | 0.0005 | 0.9648 |
| 271F | Breeder | 8.4 | 2 | 2 | 0.8154 | 720 | 0.456 | 0.0243 | 0.01 | 0.0003 | 0.9654 |
| 288F | Breeder | 4.7 | 2 | 2 | 0.8172 | 616 | 0.29 | 0.0253 | 0.009 | 0.0005 | 0.9651 |
| 295M | Breeder | 5.2 | 2 | 5 | 0.7774 | 618 | 0.349 | 0.0254 | 0.0116 | 0.0007 | 0.9622 |
| 296M | Breeder | 2.9 | 2 | 1 | 0.8083 | 771 | 0.407 | 0.0243 | 0.0091 | 0.0003 | 0.9663 |
| 301M | Breeder | 4.9 | 2 | 1 | 0.8175 | 669 | 0.308 | 0.0263 | 0.0105 | 0.0009 | 0.9624 |
| 353F | Breeder | 6.9 | 2 | 2 | 0.8054 | 606 | 0.285 | 0.0271 | 0.0085 | 0.0002 | 0.9642 |
| 386F | Breeder | 3.8 | 2 | 1 | 0.7766 | 701 | 0.349 | 0.0278 | 0.0109 | 0.0002 | 0.961 |
| 469F | Breeder | 8.9 | 2 | 4 | 0.7919 | 685 | 0.418 | 0.0274 | 0.0097 | 0.0004 | 0.9625 |
| 470F | Breeder | 6.8 | 2 | 1 | 0.8202 | 709 | 0.355 | 0.0258 | 0.0103 | 0.0005 | 0.9634 |
| 471F | Breeder | 9.0 | 2 | 2 | 0.7801 | 630 | 0.367 | 0.029 | 0.0115 | 0.0003 | 0.9592 |
| 480M | Breeder | 6.9 | 2 | 3 | 0.7895 | 675 | 0.397 | 0.0253 | 0.0099 | 0.0008 | 0.9639 |
| 486F | Breeder | 7.6 | 2 | 6 | 0.7602 | 635 | 0.33 | 0.0244 | 0.011 | 0.0004 | 0.9641 |
| 489M | Breeder | 5.0 | 2 | 1 | 0.8134 | 645 | 0.448 | 0.0238 | 0.0098 | 0.0005 | 0.9659 |
| 490M | Breeder | 5.7 | 2 | 2 | 0.8365 | 773 | 0.43 | 0.0236 | 0.0094 | 0.0003 | 0.9666 |
| 527F | Breeder | 6.5 | 2 | 2 | 0.7851 | 668 | 0.377 | 0.0251 | 0.0103 | 0.0003 | 0.9642 |
| 529F | Breeder | 2.6 | 2 |  | 0.8049 | 696 | 0.437 | 0.0253 | 0.0103 | 0.0006 | 0.9639 |
| 537F | Breeder | 7.4 | 2 | 7 | 0.7529 | 612 | 0.314 | 0.0261 | 0.0101 | 0.0004 | 0.9634 |
| 569F | Breeder | 5.5 | 2 | 4 | 0.8011 | 577 | 0.433 | 0.025 | 0.0115 | 0.0004 | 0.9631 |
| 571F | Breeder | 4.0 | 2 |  | 0.8035 | 730 | 0.365 | 0.024 | 0.0115 | 0.0006 | 0.9639 |
| 590M | Breeder | 2.7 | 2 |  | 0.7731 | 648 | 0.351 | 0.027 | 0.0124 | 0.0005 | 0.9602 |
| 625F | Breeder | 2.9 | 2 |  | 0.8083 | 668 | 0.45 | 0.0245 | 0.0101 | 0.0008 | 0.9646 |
| 639M | Breeder | 2.0 | 2 |  | 0.7859 | 700 | 0.403 | 0.0254 | 0.0109 | 0.0005 | 0.9632 |
| 640F | Breeder | 3.0 | 2 | 1 | 0.7795 | 664 | 0.364 | 0.0219 | 0.0104 | 0.0003 | 0.9674 |
| 642F | Breeder | 4.5 | 2 |  | 0.8326 | 532 | 0.248 | 0.0281 | 0.011 | 0.0003 | 0.9606 |
| 691F | Breeder | 3.8 | 2 |  | 0.791 | 627 | 0.387 | 0.0255 | 0.0104 | 0.0005 | 0.9636 |
| 694F | Breeder | 3.0 | 2 |  | 0.8086 | 729 | 0.393 | 0.0258 | 0.0101 | 0.0006 | 0.9635 |
| 733F | Breeder | 3.5 | 2 |  | 0.7913 | 672 | 0.411 | 0.0254 | 0.0106 | 0.0006 | 0.9634 |
| 752F | Breeder | 3.6 | 2 |  | 0.7903 | 722 | 0.39 | 0.026 | 0.0101 | 0.0008 | 0.9632 |
| 755M | Breeder | 8.9 | 2 | 4 | 0.7659 | 602 | 0.329 | 0.0253 | 0.0111 | 0.0005 | 0.9631 |
| 820F | Breeder | 2.4 | 2 |  | 0.7733 | 600 | 0.348 | 0.0258 | 0.0105 | 0.0005 | 0.9632 |
| 838M | Breeder | 5.1 | 2 | 1 | 0.8137 | 707 | 0.353 | 0.0254 | 0.0094 | 0.0006 | 0.9645 |
| 889F | Breeder | 3.6 | 2 |  | 0.7967 | 719 | 0.332 | 0.025 | 0.0103 | 0.0004 | 0.9642 |
| 890M | Breeder | 6.8 | 2 | 1 | 0.7802 | 708 | 0.408 | 0.0275 | 0.0102 | 0.0005 | 0.9618 |
| 907F | Breeder | 4.8 | 2 |  | 0.781 | 672 | 0.376 | 0.0222 | 0.012 | 0.0005 | 0.9653 |
| 909F | Breeder | 8.0 | 2 | 5 | 0.8142 | 681 | 0.324 | 0.0259 | 0.0101 | 0.0004 | 0.9636 |
| 963F | Breeder | 3.8 | 2 | 1 | 0.7991 | 657 | 0.428 | 0.0248 | 0.0099 | 0.0007 | 0.9647 |
| 966M | Breeder | 3.5 | 2 |  | 0.803 | 689 | 0.446 | 0.0254 | 0.0111 | 0.0005 | 0.963 |
| 969F | Breeder | 4.8 | 2 |  | 0.7845 | 622 | 0.367 | 0.025 | 0.0109 | 0.0006 | 0.9635 |
| SW147M | Breeder | 6.5 | 2 | 1 | 0.7982 | 633 | 0.39 | 0.0252 | 0.0106 | 0.0008 | 0.9635 |
| 006M | Breeder | 4.4 | 3 | 3 | 0.802 | 724 | 0.362 | 0.0259 | 0.0107 | 0.0003 | 0.9631 |
| 033F | Breeder | 6.3 | 3 | 4 | 0.7583 | 600 | 0.309 | 0.027 | 0.0115 | 0.0005 | 0.961 |
| 034M | Breeder | 7.6 | 3 | 6 | 0.7668 | 501 | 0.259 | 0.0239 | 0.01 | 0.0006 | 0.9655 |
| 040F | Breeder | 5.1 | 3 | 2 | 0.7507 | 582 | 0.304 | 0.0252 | 0.0094 | 0.0008 | 0.9647 |
| 070M | Breeder | 9.1 | 3 |  | 0.8006 | 537 | 0.248 | 0.0271 | 0.0116 | 0.0002 | 0.9611 |
| 072M | Breeder | 7.5 | 3 | 7 | 0.7796 | 621 | 0.347 | 0.0249 | 0.0108 | 0.0005 | 0.9638 |
| 1015M | Breeder | 3.8 | 3 | 3 | 0.7749 | 635 | 0.354 | 0.026 | 0.0097 | 0.0004 | 0.9639 |
| 105F | Breeder | 6.1 | 3 | 1 | 0.7553 | 593 | 0.313 | 0.024 | 0.0101 | 0.0003 | 0.9655 |
| 106F | Breeder | 8.0 | 3 | 4 | 0.747 | 567 | 0.297 | 0.0258 | 0.0102 | 0.0006 | 0.9634 |
| 120M | Breeder | 3.2 | 3 | 1 | 0.8174 | 618 | 0.472 | 0.0251 | 0.0105 | 0.0006 | 0.9639 |
| 126F | Breeder | 11.9 | 3 | 7 | 0.7793 | 645 | 0.35 | 0.026 | 0.0105 | 0.0006 | 0.9629 |
| 206M | Breeder | 6.9 | 3 | 4 | 0.7783 | 665 | 0.354 | 0.025 | 0.0095 | 0.0006 | 0.9649 |
| 209F | Breeder | 8.4 | 3 | 3 | 0.8252 | 558 | 0.262 | 0.0243 | 0.011 | 0.0006 | 0.9641 |
| 217F | Breeder | 3.7 | 3 | 1 | 0.7891 | 685 | 0.335 | 0.0268 | 0.0114 | 0.0004 | 0.9613 |
| 243M | Breeder | 4.6 | 3 | 1 | 0.8321 | 694 | 0.333 | 0.0262 | 0.0094 | 0.0005 | 0.9639 |
| 259F | Breeder | 3.0 | 3 | 1 | 0.7991 | 684 | 0.412 | 0.0258 | 0.0097 | 0.0006 | 0.9639 |
| 287M | Breeder | 4.8 | 3 | 1 | 0.7936 | 736 | 0.373 | 0.0251 | 0.0101 | 0.0004 | 0.9643 |
| 294M | Breeder | 4.0 | 3 | 3 | 0.8057 | 732 | 0.461 | 0.0247 | 0.0104 | 0.0007 | 0.9641 |
| 302M | Breeder | 9.5 | 3 | 4 | 0.7939 | 710 | 0.422 | 0.0231 | 0.0104 | 0.0005 | 0.966 |
| 377M | Breeder | 5.0 | 3 | 2 | 0.8176 | 667 | 0.486 | 0.0248 | 0.0103 | 0.0003 | 0.9646 |
| 381M | Breeder | 3.3 | 3 | 1 | 0.776 | 718 | 0.362 | 0.0254 | 0.0105 | 0.0007 | 0.9634 |
| 472F | Breeder | 10.6 | 3 | 7 | 0.7612 | 630 | 0.323 | 0.0274 | 0.01 | 0.0007 | 0.9619 |
| 482M | Breeder | 8.9 | 3 | 5 | 0.7943 | 705 | 0.415 | 0.025 | 0.0099 | 0.0005 | 0.9647 |
| 534M | Breeder | 8.4 | 3 | 7 | 0.7635 | 682 | 0.349 | 0.027 | 0.0111 | 0.0005 | 0.9615 |
| 536F | Breeder | 6.6 | 3 | 3 | 0.7817 | 641 | 0.37 | 0.0248 | 0.0091 | 0.0008 | 0.9652 |
| 588F | Breeder | 3.6 | 3 | 1 | 0.8068 | 709 | 0.426 | 0.0255 | 0.0101 | 0.0006 | 0.9638 |
| 627M | Breeder | 5.4 | 3 | 4 | 0.7958 | 714 | 0.338 | 0.0263 | 0.0097 | 0.0006 | 0.9634 |
| 636M | Breeder | 4.6 | 3 | 2 | 0.8246 | 715 | 0.488 | 0.0269 | 0.0104 | 0.0003 | 0.9623 |
| 641M | Breeder | 4.7 | 3 | 1 | 0.7891 | 720 | 0.372 | 0.0256 | 0.0104 | 0.0004 | 0.9636 |
| 689M | Breeder | 8.6 | 3 | 4 | 0.7698 | 665 | 0.343 | 0.0252 | 0.0114 | 0.0006 | 0.9627 |
| 692F | Breeder | 5.6 | 3 | 2 | 0.7746 | 605 | 0.353 | 0.0306 | 0.0112 | 0.0007 | 0.9575 |
| 693F | Breeder | 7.5 | 3 | 2 | 0.7677 | 631 | 0.327 | 0.0272 | 0.0109 | 0.0004 | 0.9614 |
| 696F | Breeder | 4.8 | 3 | 1 | 0.8279 | 764 | 0.437 | 0.0255 | 0.0092 | 0.0003 | 0.965 |
| 712M | Breeder | 10.8 | 3 | 2 | 0.789 | 727 | 0.362 | 0.0253 | 0.0105 | 0.0003 | 0.9639 |
| 715F | Breeder | 4.6 | 3 |  | 0.8163 | 729 | 0.419 | 0.0254 | 0.0112 | 0.0005 | 0.9629 |
| 757F | Breeder | 4.2 | 3 |  | 0.8274 | 793 | 0.41 | 0.0249 | 0.0106 | 0.0005 | 0.964 |
| 760M | Breeder | 6.9 | 3 | 2 | 0.7822 | 640 | 0.363 | 0.0251 | 0.0117 | 0.0006 | 0.9627 |
| 778M | Breeder | 8.7 | 3 | 1 | 0.7929 | 752 | 0.404 | 0.0254 | 0.0096 | 0.0006 | 0.9644 |
| 779F | Breeder | 7.8 | 3 | 3 | 0.7774 | 686 | 0.368 | 0.0254 | 0.0101 | 0.0004 | 0.964 |
| 911M | Breeder | 6.4 | 3 | 3 | 0.7869 | 703 | 0.382 | 0.0258 | 0.0105 | 0.0002 | 0.9635 |
| 926F | Breeder | 6.8 | 3 | 2 | 0.7742 | 580 | 0.343 | 0.0254 | 0.0097 | 0.0005 | 0.9644 |
| 953F | Breeder | 4.8 | 3 | 1 | 0.8185 | 619 | 0.461 | 0.0252 | 0.0108 | 0.0006 | 0.9635 |
| 1118F | Breeder | 3.8 | 4* |  | 0.7812 | 691 | 0.381 | 0.0248 | 0.0099 | 0.0006 | 0.9647 |
| 103F | Breeder | 7.0 | 4 | 1 | 0.7469 | 586 | 0.294 | 0.0225 | 0.0116 | 0.0004 | 0.9655 |
| 151F | Breeder | 11.5 | 4 | 6 | 0.8027 | 693 | 0.428 | 0.026 | 0.01 | 0.0004 | 0.9636 |
| 190F | Breeder | 5.5 | 4 |  | 0.7942 | 675 | 0.407 | 0.0239 | 0.0102 | 0.0002 | 0.9656 |
| 205M | Breeder | 4.5 | 4 | 1 | 0.7631 | 606 | 0.317 | 0.0271 | 0.0119 | 0.0007 | 0.9603 |
| 227M | Breeder | 4.9 | 4 | 1 | 0.8078 | 737 | 0.392 | 0.0253 | 0.0112 | 0.0005 | 0.963 |
| 303M | Breeder | 10.9 | 4 | 7 | 0.8201 | 793 | 0.432 | 0.0247 | 0.01 | 0.0005 | 0.9647 |
| 383M | Breeder | 6.9 | 4 | 2 | 0.7734 | 572 | 0.326 | 0.0257 | 0.0112 | 0.0005 | 0.9626 |
| 540F | Breeder | 7.5 | 4 | 2 | 0.7537 | 590 | 0.297 | 0.0269 | 0.0119 | 0.0003 | 0.9609 |
| 541M | Breeder | 7.5 | 4 | 2 | 0.7787 | 663 | 0.369 | 0.0232 | 0.0111 | 0.0002 | 0.9654 |
| 685M | Breeder | 6.9 | 4 | 1 | 0.8246 | 764 | 0.457 | 0.0259 | 0.0104 | 0.0003 | 0.9634 |
| 759F | Breeder | 3.9 | 4* |  | 0.812 | 583 | 0.454 | 0.0257 | 0.0097 | 0.0005 | 0.9641 |
| 871M | Breeder | 4.5 | 4 | 1 | 0.8012 | 724 | 0.389 | 0.0259 | 0.0118 | 0.0006 | 0.9617 |
| 978F | Breeder | 4.8 | 4 |  | 0.7763 | 644 | 0.357 | 0.0277 | 0.0101 | 0.0007 | 0.9615 |
| SW763M | Breeder | 9.9 | 4 | 3 | 0.7887 | 619 | 0.393 | 0.0242 | 0.0122 | 0.0006 | 0.963 |
| 192M | Breeder | 12.1 | 5 | 1 | 0.8225 | 799 | 0.412 | 0.0247 | 0.01 | 0.0004 | 0.9649 |
| 207M | Breeder | 7.4 | 5 | 1 | 0.7981 | 706 | 0.42 | 0.024 | 0.0098 | 0.0005 | 0.9657 |
| 344F | Breeder | 5.3 | 5 | 1 | 0.8117 | 696 | 0.453 | 0.0255 | 0.0099 | 0.0004 | 0.9641 |
| 545F | Breeder | 5.3 | 5 | 1 | 0.7995 | 745 | 0.412 | 0.0258 | 0.0107 | 0.0003 | 0.9631 |
| 586M | Breeder | 9.5 | 5 | 1 | 0.8192 | 651 | 0.467 | 0.0246 | 0.0106 | 0.0003 | 0.9645 |
| 695M | Breeder | 7.6 | 5 | 1 | 0.7949 | 674 | 0.409 | 0.0254 | 0.0103 | 0.0002 | 0.9641 |
| 794F | Breeder | 7.1 | 5 | 2 | 0.7815 | 687 | 0.333 | 0.0255 | 0.0106 | 0.0005 | 0.9634 |
| 925M | Breeder | 5.9 | 5 | 2 | 0.7749 | 564 | 0.363 | 0.0228 | 0.0111 | 0.0007 | 0.9653 |
| 970F | Breeder | 6.0 | 5 | 2 | 0.7713 | 651 | 0.349 | 0.0252 | 0.0104 | 0.0005 | 0.9639 |
| 976F | Breeder | 7.8 | 5 | 1 | 0.8082 | 716 | 0.417 | 0.0281 | 0.0103 | 0.0005 | 0.9611 |
| 992M | Breeder | 5.0 | 5 |  | 0.7998 | 706 | 0.438 | 0.0251 | 0.0106 | 0.0005 | 0.9638 |
| 478F | Breeder | 12.6 | 6 | 4 | 0.827 | 788 | 0.483 | 0.0258 | 0.0099 | 0.0005 | 0.9638 |
| 495M | Breeder | 8.4 | 6 | 6 | 0.813 | 677 | 0.461 | 0.0252 | 0.0109 | 0.0005 | 0.9634 |
| 587M | Breeder | 9.1 | 6 | 1 | 0.8304 | 720 | 0.503 | 0.0254 | 0.0112 | 0.0004 | 0.963 |
| 633F | Breeder | 7.4 | 6 | 1 | 0.7971 | 676 | 0.395 | 0.0257 | 0.0105 | 0.0005 | 0.9634 |
| 686F | Breeder | 6.4 | 6 |  | 0.7854 | 713 | 0.371 | 0.0256 | 0.0108 | 0.0003 | 0.9633 |
| 949M | Breeder | 6.4 | 6 |  | 0.7792 | 670 | 0.381 | 0.0243 | 0.0103 | 0.0005 | 0.9649 |
| 910M | Breeder | 8.8 | 8 |  | 0.837 | 596 | 0.531 | 0.025 | 0.0102 | 0.0004 | 0.9644 |
| 005F | Breeder | 10.6 |  | 4 | 0.7975 | 765 | 0.418 | 0.0255 | 0.0108 | 0.0005 | 0.9633 |
| 009F | Breeder | 9.6 |  | 3 | 0.8006 | 732 | 0.415 | 0.0258 | 0.0097 | 0.0006 | 0.9639 |
| 013M | Breeder | 11.9 |  | 3 | 0.7674 | 593 | 0.298 | 0.0234 | 0.0095 | 0.0003 | 0.9667 |
| 027F | Breeder | 8.5 |  | 2 | 0.7466 | 578 | 0.3 | 0.0253 | 0.0106 | 0.0007 | 0.9634 |
| 035M | Breeder | 5.8 |  | 2 | 0.769 | 632 | 0.305 | 0.0268 | 0.0105 | 0.0006 | 0.9621 |
| 036F | Breeder | 3.0 |  | 1 | 0.7595 | 611 | 0.312 | 0.0242 | 0.0101 | 0.0011 | 0.9646 |
| 038M | Breeder | 8.7 |  | 2 | 0.7669 | 529 | 0.288 | 0.0219 | 0.0101 | 0.0008 | 0.9672 |
| 104M | Breeder | 3.6 |  |  | 0.7483 | 574 | 0.29 | 0.025 | 0.0115 | 0.0003 | 0.9632 |
| 614M | Breeder |  |  | 1 | 0.8043 | 674 | 0.436 | 0.0257 | 0.0101 | 0.0006 | 0.9637 |

|  |  |  |  |  |  |  |  |  |  |  |
| --- | --- | --- | --- | --- | --- | --- | --- | --- | --- | --- |
| 821F | Breeder | 8.0 | 1 | 0.7924 | 677 | 0.422 | 0.0242 | 0.011 | 0.0002 | 0.9645 |
| 1005F | Non-breeder | 2.8 |  | 0.7905 | 675 | 0.391 | 0.0272 | 0.0097 | 0.0003 | 0.9628 |
| 1014M | Non-breeder | 4.8 |  | 0.7793 | 684 | 0.386 | 0.0279 | 0.0105 | 0.0008 | 0.9609 |
| 1048M | Non-breeder | 2.8 |  | 0.8006 | 725 | 0.434 | 0.0269 | 0.0111 | 0.0006 | 0.9615 |
| 1049F | Non-breeder | 1.8 |  | 0.8007 | 662 | 0.396 | 0.027 | 0.0105 | 0.0004 | 0.9622 |
| 1090F | Non-breeder | 3.8 |  | 0.7781 | 653 | 0.366 | 0.0244 | 0.0106 | 0.0008 | 0.9642 |
| 1104F | Non-breeder | 0.8 |  | 0.8036 | 728 | 0.389 | 0.0281 | 0.011 | 0.0009 | 0.96 |
| 1106M | Non-breeder | 0.8 |  | 0.7898 | 688 | 0.406 | 0.0259 | 0.0116 | 0.0003 | 0.9622 |
| 1119F | Non-breeder | 1.8 |  | 0.7794 | 651 | 0.381 | 0.025 | 0.0107 | 0.0005 | 0.9638 |
| 118M | Non-breeder | 2.0 |  | 0.8048 | 655 | 0.447 | 0.0271 | 0.0106 | 0.0004 | 0.9618 |
| 147M | Non-breeder | 3.2 |  | 0.7666 | 603 | 0.323 | 0.025 | 0.0107 | 0.0007 | 0.9636 |
| 241M | Non-breeder | 1.0 |  | 0.8142 | 748 | 0.456 | 0.0244 | 0.01 | 0.0002 | 0.9654 |
| 253M | Non-breeder | 4.8 |  | 0.7847 | 704 | 0.369 | 0.0265 | 0.0087 | 0.0003 | 0.9645 |
| 289M | Non-breeder | 1.7 |  | 0.7919 | 717 | 0.389 | 0.0273 | 0.0117 | 0.0004 | 0.9605 |
| 292M | Non-breeder | 2.9 |  | 0.7846 | 650 | 0.373 | 0.0252 | 0.0113 | 0.0004 | 0.9632 |
| 350M | Non-breeder | 1.4 |  | 0.7798 | 722 | 0.376 | 0.0281 | 0.0122 | 0.0005 | 0.9592 |
| 389M | Non-breeder | 2.1 |  | 0.8258 | 701 | 0.506 | 0.0245 | 0.0104 | 0.0005 | 0.9646 |
| 485F | Non-breeder | 1.9 |  | 0.7886 | 606 | 0.28 | 0.0227 | 0.01 | 0.0006 | 0.9668 |
| 525F | Non-breeder | 3.7 |  | 0.7856 | 581 | 0.388 | 0.0257 | 0.0105 | 0.0005 | 0.9633 |
| 698M | Non-breeder | 4.1 |  | 0.764 | 637 | 0.328 | 0.026 | 0.0104 | 0.0007 | 0.9628 |
| 015M | Non-breeder | 2.1 |  | 0.8052 | 428 | 0.174 | 0.0251 | 0.0105 | 0.0005 | 0.964 |
| 020M | Non-breeder | 1.2 |  | 0.7932 | 740 | 0.393 | 0.0275 | 0.0102 | 0.0007 | 0.9616 |
| 022M | Non-breeder | 0.7 |  | 0.8016 | 758 | 0.392 | 0.0254 | 0.011 | 0.0006 | 0.963 |

|  |  |  |  |  |  |  |  |  |  |
| --- | --- | --- | --- | --- | --- | --- | --- | --- | --- |
| 024F | Non-breeder | 3.5 | 0.7801 | 613 | 0.371 | 0.0258 | 0.0112 | 0.0005 | 0.9625 |
| 026F | Non-breeder | 1.4 | 0.7753 | 514 | 0.274 | 0.0201 | 0.0099 | 0.0005 | 0.9694 |
| 031M | Non-breeder | 2.6 | 0.7856 | 525 | 0.274 | 0.0197 | 0.01 | 0.0003 | 0.97 |
| 043M | Non-breeder | 2.9 | 0.7819 | 685 | 0.362 | 0.024 | 0.0111 | 0.0004 | 0.9645 |
| 044F | Non-breeder | 9.0 | 0.8019 | 648 | 0.318 | 0.0277 | 0.0116 | 0.0005 | 0.9602 |
| 045F | Non-breeder | 0.4 | 0.7942 | 676 | 0.381 | 0.0255 | 0.0118 | 0.0007 | 0.962 |
| 055M | Non-breeder | 3.9 | 0.7909 | 715 | 0.413 | 0.0242 | 0.01 | 0.0006 | 0.9652 |
| 058F | Non-breeder | 0.0 | 0.8299 | 511 | 0.514 | 0.0249 | 0.0108 | 0.0003 | 0.964 |
| 063F | Non-breeder | 1.5 | 0.7777 | 648 | 0.333 | 0.0238 | 0.0107 | 0.0004 | 0.9651 |
| 064F | Non-breeder | 1.1 | 0.7754 | 609 | 0.345 | 0.0246 | 0.0109 | 0.0004 | 0.964 |
| 065F | Non-breeder | 1.1 | 0.7765 | 629 | 0.354 | 0.0244 | 0.0107 | 0.0002 | 0.9647 |
| 068F | Non-breeder | 1.4 | 0.7854 | 568 | 0.314 | 0.0212 | 0.0106 | 0.0001 | 0.968 |
| 069M | Non-breeder | 1.2 | 0.7738 | 636 | 0.345 | 0.027 | 0.0103 | 0.0007 | 0.9619 |
| 071F | Non-breeder | 1.1 | 0.7753 | 609 | 0.341 | 0.0261 | 0.0106 | 0.0002 | 0.9631 |
| 077F | Non-breeder | 3.7 | 0.8109 | 678 | 0.347 | 0.0244 | 0.0105 | 0.0004 | 0.9647 |
| 082M | Non-breeder | 2.1 | 0.804 | 781 | 0.417 | 0.0257 | 0.0099 | 0.0004 | 0.964 |
| 083M | Non-breeder | 1.5 | 0.7931 | 696 | 0.406 | 0.0257 | 0.0102 | 0.0006 | 0.9635 |
| 085F | Non-breeder | 1.6 | 0.8188 | 683 | 0.337 | 0.0264 | 0.0096 | 0.0003 | 0.9636 |
| 095F | Non-breeder | 1.1 | 0.8311 | 537 | 0.24 | 0.0265 | 0.0092 | 0.0003 | 0.964 |
| 1012M | Non-breeder | 0.9 | 0.7914 | 651 | 0.418 | 0.0251 | 0.0112 | 0.0006 | 0.9631 |
| 1013M | Non-breeder | 2.7 | 0.7771 | 670 | 0.361 | 0.0256 | 0.0106 | 0.0006 | 0.9633 |
| 1016F | Non-breeder | 0.8 | 0.7749 | 682 | 0.355 | 0.0259 | 0.0104 | 0.0008 | 0.963 |
| 1047M | Non-breeder | 3.8 | 0.806 | 666 | 0.442 | 0.0268 | 0.0114 | 0.0006 | 0.9612 |
| 1050F | Non-breeder | 1.0 | 0.8006 | 675 | 0.423 | 0.0267 | 0.0096 | 0.0005 | 0.9632 |
| 1051M | Non-breeder | 0.8 | 0.8041 | 707 | 0.445 | 0.0247 | 0.0104 | 0.0006 | 0.9643 |
| 1094M | Non-breeder | 0.0 | 0.8163 | 663 | 0.307 | 0.0258 | 0.0105 | 0.0005 | 0.9632 |
| 1095F | Non-breeder | 0.0 | 0.8022 | 703 | 0.442 | 0.0268 | 0.0095 | 0.0004 | 0.9633 |
| 1096M | Non-breeder | 0.0 | 0.792 | 685 | 0.404 | 0.0277 | 0.0098 | 0.0005 | 0.962 |
| 1097F | Non-breeder | 0.0 | 0.7997 | 723 | 0.431 | 0.0276 | 0.0096 | 0.0005 | 0.9622 |
| 1098M | Non-breeder | 0.0 | 0.8039 | 751 | 0.43 | 0.027 | 0.0098 | 0.0003 | 0.9629 |
| 1108M | Non-breeder | 1.8 | 0.7964 | 704 | 0.411 | 0.0261 | 0.0105 | 0.0006 | 0.9627 |
| 111F | Non-breeder | 1.2 | 0.768 | 614 | 0.309 | 0.0277 | 0.011 | 0.0006 | 0.9608 |
| 115F | Non-breeder | 1.3 | 0.7818 | 632 | 0.309 | 0.0252 | 0.0114 | 0.0009 | 0.9625 |
| 122M | Non-breeder | 2.3 | 0.8147 | 627 | 0.475 | 0.026 | 0.0105 | 0.0006 | 0.9629 |
| 123M | Non-breeder | 2.6 | 0.7836 | 698 | 0.369 | 0.0254 | 0.0102 | 0.0006 | 0.9638 |
| 124M | Non-breeder | 2.7 | 0.7722 | 599 | 0.356 | 0.0248 | 0.0121 | 0.0006 | 0.9624 |
| 128M | Non-breeder | 1.1 | 0.7649 | 628 | 0.342 | 0.0255 | 0.0107 | 0.0005 | 0.9633 |
| 136F | Non-breeder | 2.9 | 0.8176 | 673 | 0.461 | 0.027 | 0.0102 | 0.0005 | 0.9623 |
| 150M | Non-breeder | 6.5 | 0.7871 | 673 | 0.399 | 0.0256 | 0.0099 | 0.0004 | 0.9641 |
| 153F | Non-breeder | 1.6 | 0.7911 | 711 | 0.424 | 0.0273 | 0.0105 | 0.0003 | 0.9619 |
| 154F | Non-breeder | 2.6 | 0.7887 | 707 | 0.413 | 0.0246 | 0.0096 | 0.0005 | 0.9653 |
| 156F | Non-breeder | 2.3 | 0.7928 | 710 | 0.42 | 0.026 | 0.0102 | 0.0004 | 0.9634 |
| 160F | Non-breeder | 0.8 | 0.8039 | 721 | 0.445 | 0.0257 | 0.0096 | 0.0005 | 0.9642 |
| 161M | Non-breeder | 2.9 | 0.796 | 702 | 0.425 | 0.0252 | 0.01 | 0.0005 | 0.9643 |
| 162M | Non-breeder | 4.0 | 0.8075 | 763 | 0.399 | 0.0251 | 0.0105 | 0.0001 | 0.9643 |
| 168F | Non-breeder | 1.6 | 0.8204 | 780 | 0.412 | 0.0264 | 0.0107 | 0.0003 | 0.9627 |
| 182M | Non-breeder | 0.5 | 0.7958 | 699 | 0.416 | 0.0254 | 0.0107 | 0.0004 | 0.9635 |
| 183F | Non-breeder | 0.5 | 0.819 | 708 | 0.454 | 0.0256 | 0.0108 | 0.0006 | 0.963 |
| 184F | Non-breeder | 0.5 | 0.8138 | 696 | 0.449 | 0.0265 | 0.011 | 0.0007 | 0.9618 |
| 185M | Non-breeder | 0.6 | 0.7826 | 689 | 0.338 | 0.0249 | 0.0105 | 0.0004 | 0.9642 |
| 186M | Non-breeder | 0.6 | 0.7632 | 609 | 0.33 | 0.025 | 0.0106 | 0.0007 | 0.9638 |
| 187F | Non-breeder | 0.8 | 0.7899 | 602 | 0.277 | 0.0239 | 0.0123 | 0.0005 | 0.9634 |
| 188F | Non-breeder | 2.2 | 0.7995 | 643 | 0.439 | 0.0253 | 0.0102 | 0.0006 | 0.964 |
| 189M | Non-breeder | 1.9 | 0.8125 | 609 | 0.443 | 0.0261 | 0.0114 | 0.0003 | 0.9622 |
| 191M | Non-breeder | 1.9 | 0.7716 | 658 | 0.325 | 0.0266 | 0.0108 | 0.0006 | 0.9621 |
| 196M | Non-breeder | 3.3 | 0.8083 | 781 | 0.44 | 0.0256 | 0.0115 | 0.0002 | 0.9627 |
| 197F | Non-breeder | 1.1 | 0.8348 | 618 | 0.279 | 0.0284 | 0.0094 | 0.0003 | 0.9619 |
| 199M | Non-breeder | 1.1 | 0.7722 | 667 | 0.35 | 0.0277 | 0.0105 | 0.0002 | 0.9616 |
| 201F | Non-breeder | 2.5 | 0.7574 | 619 | 0.321 | 0.0265 | 0.0109 | 0.0007 | 0.962 |
| 203M | Non-breeder | 1.4 | 0.7584 | 619 | 0.306 | 0.0264 | 0.011 | 0.0005 | 0.962 |
| 210M | Non-breeder | 3.0 | 0.8141 | 628 | 0.479 | 0.0253 | 0.0101 | 0.0006 | 0.964 |
| 211M | Non-breeder | 2.8 | 0.8277 | 594 | 0.288 | 0.0235 | 0.009 | 0.0004 | 0.967 |
| 215M | Non-breeder | 2.3 | 0.7769 | 669 | 0.325 | 0.0235 | 0.0109 | 0.0007 | 0.965 |
| 216F | Non-breeder | 1.8 | 0.7862 | 682 | 0.324 | 0.0264 | 0.0117 | 0.0003 | 0.9616 |
| 218F | Non-breeder | 2.3 | 0.7918 | 666 | 0.315 | 0.0258 | 0.0103 | 0.0004 | 0.9635 |
| 221M | Non-breeder | 1.9 | 0.7656 | 640 | 0.325 | 0.024 | 0.0103 | 0.0002 | 0.9656 |
| 222M | Non-breeder | 1.2 | 0.7613 | 626 | 0.342 | 0.0271 | 0.0108 | 0.0002 | 0.9619 |
| 225M | Non-breeder | 0.9 | 0.7937 | 635 | 0.408 | 0.0263 | 0.0102 | 0.0005 | 0.963 |
| 226M | Non-breeder | 0.8 | 0.788 | 684 | 0.391 | 0.0247 | 0.0105 | 0.0004 | 0.9645 |
| 229M | Non-breeder | 2.2 | 0.8027 | 708 | 0.398 | 0.0262 | 0.0108 | 0.0005 | 0.9625 |
| 245M | Non-breeder | 0.7 | 0.7889 | 617 | 0.394 | 0.0247 | 0.0106 | 0.0005 | 0.9642 |
| 246M | Non-breeder | 0.7 | 0.7931 | 591 | 0.389 | 0.0275 | 0.0111 | 0.0004 | 0.9609 |
| 247M | Non-breeder | 0.7 | 0.7925 | 643 | 0.404 | 0.0265 | 0.0106 | 0.0005 | 0.9624 |
| 248M | Non-breeder | 1.3 | 0.7547 | 588 | 0.304 | 0.0256 | 0.0105 | 0.0005 | 0.9634 |
| 249M | Non-breeder | 2.7 | 0.7648 | 593 | 0.32 | 0.0262 | 0.0119 | 0.0003 | 0.9616 |
| 252M | Non-breeder | 2.1 | 0.7764 | 660 | 0.313 | 0.0265 | 0.0118 | 0.0003 | 0.9614 |
| 254M | Non-breeder | 2.3 | 0.7723 | 720 | 0.353 | 0.0267 | 0.0101 | 0.0006 | 0.9626 |
| 257M | Non-breeder | 4.5 | 0.8046 | 660 | 0.444 | 0.0259 | 0.0089 | 0.0006 | 0.9645 |
| 260F | Non-breeder | 2.3 | 0.7906 | 657 | 0.404 | 0.0244 | 0.0097 | 0.0004 | 0.9655 |
| 291M | Non-breeder | 2.3 | 0.8173 | 812 | 0.444 | 0.0257 | 0.0096 | 0.0004 | 0.9643 |
| 293F | Non-breeder | 2.1 | 0.8181 | 707 | 0.427 | 0.0234 | 0.0102 | 0.0008 | 0.9656 |
| 300M | Non-breeder | 4.5 | 0.8024 | 751 | 0.423 | 0.0253 | 0.0095 | 0.0005 | 0.9646 |
| 305M | Non-breeder | 3.0 | 0.7556 | 613 | 0.315 | 0.0231 | 0.0104 | 0.0008 | 0.9657 |
| 307M | Non-breeder | 2.8 | 0.7961 | 683 | 0.406 | 0.0267 | 0.0103 | 0.0006 | 0.9625 |
| 308F | Non-breeder | 0.8 | 0.8021 | 696 | 0.421 | 0.0242 | 0.0105 | 0.0007 | 0.9646 |
| 309F | Non-breeder | 2.8 | 0.7671 | 659 | 0.337 | 0.0258 | 0.0109 | 0.0006 | 0.9627 |
| 341F | Non-breeder | 5.8 | 0.8084 | 583 | 0.263 | 0.0267 | 0.0124 | 0.0001 | 0.9608 |
| 343M | Non-breeder | 1.7 | 0.8333 | 801 | 0.459 | 0.0257 | 0.0104 | 0.0006 | 0.9633 |
| 347M | Non-breeder | 1.6 | 0.8053 | 593 | 0.427 | 0.0246 | 0.0116 | 0.0005 | 0.9634 |
| 348M | Non-breeder | 1.7 | 0.7902 | 728 | 0.411 | 0.0259 | 0.0099 | 0.0007 | 0.9635 |
| 349M | Non-breeder | 1.8 | 0.7956 | 733 | 0.422 | 0.0262 | 0.011 | 0.0002 | 0.9626 |
| 352M | Non-breeder | 1.7 | 0.7967 | 648 | 0.436 | 0.0269 | 0.01 | 0.0006 | 0.9625 |
| 355F | Non-breeder | 3.3 | 0.7942 | 659 | 0.419 | 0.0241 | 0.0104 | 0.0005 | 0.965 |
| 356M | Non-breeder | 2.7 | 0.8168 | 751 | 0.441 | 0.0246 | 0.0107 | 0.0006 | 0.9641 |
| 361M | Non-breeder | 0.7 | 0.8112 | 734 | 0.383 | 0.0257 | 0.0114 | 0.0008 | 0.9621 |
| 375F | Non-breeder | 1.8 | 0.7912 | 700 | 0.406 | 0.0263 | 0.0103 | 0.0006 | 0.9628 |
| 379M | Non-breeder | 4.6 | 0.8222 | 811 | 0.428 | 0.026 | 0.0116 | 0.0002 | 0.9622 |
| 382F | Non-breeder | 1.2 | 0.7903 | 657 | 0.39 | 0.027 | 0.0097 | 0.0005 | 0.9629 |
| 388F | Non-breeder | 3.3 | 0.8136 | 746 | 0.468 | 0.0253 | 0.0105 | 0.0006 | 0.9636 |
| 391F | Non-breeder | 3.1 | 0.7631 | 642 | 0.342 | 0.0249 | 0.0108 | 0.0006 | 0.9637 |
| 392M | Non-breeder | 2.6 | 0.7553 | 620 | 0.3 | 0.026 | 0.0102 | 0.0002 | 0.9636 |
| 451F | Non-breeder | 0.4 | 0.7847 | 704 | 0.36 | 0.0242 | 0.0106 | 0.0004 | 0.9648 |
| 453M | Non-breeder | 2.2 | 0.7842 | 657 | 0.402 | 0.0257 | 0.011 | 0.0006 | 0.9626 |
| 466U | Non-breeder | 0.7 | 0.7878 | 703 | 0.344 | 0.0242 | 0.0112 | 0.0003 | 0.9642 |
| 467M | Non-breeder | 1.6 | 0.7625 | 615 | 0.319 | 0.0231 | 0.0107 | 0.0004 | 0.9658 |
| 468M | Non-breeder | 3.9 | 0.8037 | 741 | 0.405 | 0.0227 | 0.0101 | 0.0008 | 0.9664 |
| 473M | Non-breeder | 1.0 | 0.8131 | 659 | 0.316 | 0.0259 | 0.01 | 0.0009 | 0.9632 |
| 474M | Non-breeder | 1.2 | 0.7916 | 625 | 0.296 | 0.0251 | 0.0112 | 0.0002 | 0.9634 |
| 475F | Non-breeder | 0.8 | 0.7748 | 647 | 0.344 | 0.0232 | 0.0109 | 0.0003 | 0.9656 |
| 477M | Non-breeder | 2.5 | 0.8186 | 736 | 0.48 | 0.0256 | 0.0098 | 0.0004 | 0.9642 |
| 479M | Non-breeder | 3.4 | 0.8104 | 679 | 0.455 | 0.0274 | 0.0101 | 0.0005 | 0.962 |
| 481M | Non-breeder | 1.9 | 0.768 | 649 | 0.354 | 0.0243 | 0.0105 | 0.0006 | 0.9646 |
| 483F | Non-breeder | 2.2 | 0.7608 | 660 | 0.337 | 0.0245 | 0.0101 | 0.0007 | 0.9648 |
| 484M | Non-breeder | 1.3 | 0.7671 | 610 | 0.313 | 0.0277 | 0.0113 | 0.0009 | 0.9601 |
| 488M | Non-breeder | 1.3 | 0.7736 | 665 | 0.349 | 0.0244 | 0.012 | 0.0003 | 0.9633 |
| 491M | Non-breeder | 1.8 | 0.8272 | 721 | 0.492 | 0.0252 | 0.0103 | 0.0006 | 0.964 |
| 493M | Non-breeder | 0.8 | 0.7927 | 649 | 0.398 | 0.0262 | 0.0104 | 0.0004 | 0.963 |
| 496F | Non-breeder | 1.1 | 0.8077 | 654 | 0.445 | 0.0269 | 0.0106 | 0.0007 | 0.9618 |
| 497M | Non-breeder | 3.1 | 0.818 | 643 | 0.484 | 0.0254 | 0.0103 | 0.0003 | 0.964 |
| 498F | Non-breeder | 0.9 | 0.7785 | 654 | 0.317 | 0.0263 | 0.0106 | 0.0003 | 0.9628 |
| 523F | Non-breeder | 3.0 | 0.796 | 576 | 0.406 | 0.0249 | 0.0097 | 0.0007 | 0.9646 |
| 524F | Non-breeder | 2.2 | 0.7813 | 666 | 0.363 | 0.0273 | 0.011 | 0.0006 | 0.9611 |
| 528M | Non-breeder | 2.8 | 0.7814 | 668 | 0.378 | 0.0284 | 0.0105 | 0.0008 | 0.9603 |
| 535M | Non-breeder | 3.1 | 0.8011 | 644 | 0.348 | 0.0223 | 0.0096 | 0.0006 | 0.9675 |
| 538M | Non-breeder | 6.1 | 0.8 | 735 | 0.43 | 0.0242 | 0.0109 | 0.0004 | 0.9645 |
| 539F | Non-breeder | 3.9 | 0.774 | 658 | 0.354 | 0.0262 | 0.0107 | 0.0004 | 0.9627 |
| 542F | Non-breeder | 3.0 | 0.7985 | 674 | 0.4 | 0.0259 | 0.01 | 0.0004 | 0.9636 |
| 543M | Non-breeder | 2.6 | 0.8029 | 764 | 0.392 | 0.0256 | 0.0101 | 0.0005 | 0.9637 |
| 544M | Non-breeder | 2.7 | 0.8039 | 635 | 0.436 | 0.0252 | 0.01 | 0.0005 | 0.9644 |
| 546M | Non-breeder | 1.8 | 0.8302 | 762 | 0.433 | 0.0266 | 0.0107 | 0.0006 | 0.9621 |
| 570M | Non-breeder | 1.5 | 0.7947 | 652 | 0.417 | 0.0259 | 0.0103 | 0.0003 | 0.9635 |
| 572F | Non-breeder | 6.1 | 0.7995 | 634 | 0.396 | 0.0268 | 0.0108 | 0.0005 | 0.9619 |
| 577M | Non-breeder | 1.9 | 0.7849 | 692 | 0.342 | 0.0241 | 0.0097 | 0.0006 | 0.9656 |
| 589F | Non-breeder | 2.4 | 0.7977 | 668 | 0.435 | 0.0272 | 0.01 | 0.0004 | 0.9624 |
| 591F | Non-breeder | 2.3 | 0.7913 | 684 | 0.389 | 0.0251 | 0.0115 | 0.0004 | 0.963 |
| 593F | Non-breeder | 1.8 | 0.793 | 633 | 0.397 | 0.0264 | 0.0108 | 0.0007 | 0.9621 |
| 613M | Non-breeder | 2.3 | 0.7918 | 647 | 0.393 | 0.0249 | 0.0106 | 0.0007 | 0.9638 |
| 620M | Non-breeder | 0.6 | 0.7913 | 630 | 0.401 | 0.026 | 0.0113 | 0.0003 | 0.9623 |
| 621F | Non-breeder | 2.6 | 0.791 | 661 | 0.408 | 0.0252 | 0.0102 | 0.0005 | 0.964 |
| 622F | Non-breeder | 0.7 | 0.7624 | 619 | 0.316 | 0.0264 | 0.0111 | 0.0005 | 0.962 |
| 624F | Non-breeder | 1.3 | 0.8167 | 772 | 0.415 | 0.0253 | 0.0111 | 0.0007 | 0.9628 |
| 626F | Non-breeder | 1.4 | 0.8021 | 640 | 0.46 | 0.025 | 0.0105 | 0.0004 | 0.9642 |
| 629M | Non-breeder | 2.3 | 0.7848 | 661 | 0.377 | 0.0251 | 0.0111 | 0.0005 | 0.9633 |
| 630F | Non-breeder | 1.6 | 0.7881 | 648 | 0.397 | 0.0264 | 0.0109 | 0.0005 | 0.9623 |
| 634F | Non-breeder | 1.9 | 0.7769 | 622 | 0.359 | 0.0253 | 0.0109 | 0.0007 | 0.9631 |
| 635F | Non-breeder | 1.8 | 0.7994 | 656 | 0.416 | 0.0258 | 0.0105 | 0.0005 | 0.9632 |
| 638M | Non-breeder | 1.0 | 0.7878 | 638 | 0.398 | 0.0261 | 0.0106 | 0.0007 | 0.9626 |
| 643F | Non-breeder | 1.3 | 0.7868 | 593 | 0.38 | 0.0246 | 0.0109 | 0.0004 | 0.9642 |
| 644F | Non-breeder | 1.5 | 0.7915 | 637 | 0.398 | 0.0278 | 0.0112 | 0.0002 | 0.9609 |
| 645F | Non-breeder | 2.1 | 0.7974 | 734 | 0.4 | 0.0259 | 0.0104 | 0.0007 | 0.963 |
| 646F | Non-breeder | 2.5 | 0.77 | 667 | 0.351 | 0.0244 | 0.0113 | 0.0003 | 0.964 |
| 647M | Non-breeder | 3.9 | 0.7778 | 692 | 0.36 | 0.0263 | 0.0103 | 0.0003 | 0.9631 |
| 658M | Non-breeder | 3.9 | 0.8222 | 707 | 0.349 | 0.0265 | 0.011 | 0.0005 | 0.962 |
| 659M | Non-breeder | 3.9 | 0.8176 | 615 | 0.3 | 0.0266 | 0.0111 | 0.0006 | 0.9617 |
| 660F | Non-breeder | 1.9 | 0.7801 | 642 | 0.365 | 0.0256 | 0.0103 | 0.0007 | 0.9634 |
| 661M | Non-breeder | 8.4 | 0.7925 | 678 | 0.364 | 0.0229 | 0.0106 | 0.0008 | 0.9658 |
| 676M | Non-breeder | 0.3 | 0.7905 | 663 | 0.392 | 0.0254 | 0.0116 | 0.0005 | 0.9626 |
| 679F | Non-breeder | 1.6 | 0.8198 | 710 | 0.353 | 0.0243 | 0.0102 | 0.0007 | 0.9648 |
| 687M | Non-breeder | 1.6 | 0.7849 | 619 | 0.291 | 0.0263 | 0.0099 | 0.0005 | 0.9634 |
| 688M | Non-breeder | 2.0 | 0.8013 | 586 | 0.413 | 0.0271 | 0.0099 | 0.0007 | 0.9624 |
| 690F | Non-breeder | 2.0 | 0.815 | 757 | 0.393 | 0.0258 | 0.0096 | 0.0005 | 0.9641 |
| 697M | Non-breeder | 2.7 | 0.7961 | 712 | 0.408 | 0.0256 | 0.0109 | 0.0005 | 0.963 |
| 713F | Non-breeder | 1.9 | 0.7993 | 609 | 0.405 | 0.0247 | 0.0096 | 0.0003 | 0.9654 |
| 716F | Non-breeder | 4.4 | 0.7952 | 678 | 0.41 | 0.0255 | 0.0102 | 0.0004 | 0.9639 |
| 729M | Non-breeder | 1.1 | 0.7931 | 659 | 0.411 | 0.025 | 0.0098 | 0.0007 | 0.9645 |
| 734M | Non-breeder | 2.6 | 0.8 | 661 | 0.435 | 0.0256 | 0.0096 | 0.0004 | 0.9644 |
| 753F | Non-breeder | 1.3 | 0.7768 | 669 | 0.355 | 0.0258 | 0.0107 | 0.0005 | 0.963 |
| 754M | Non-breeder | 4.6 | 0.7694 | 648 | 0.333 | 0.024 | 0.0112 | 0.0002 | 0.9646 |
| 756F | Non-breeder | 3.8 | 0.7936 | 726 | 0.411 | 0.0263 | 0.0104 | 0.0006 | 0.9627 |
| 758M | Non-breeder | 3.9 | 0.8342 | 734 | 0.484 | 0.0241 | 0.0098 | 0.0002 | 0.9659 |
| 761F | Non-breeder | 2.9 | 0.7954 | 733 | 0.419 | 0.0231 | 0.0099 | 0.0006 | 0.9664 |
| 762M | Non-breeder | 6.3 | 0.8084 | 665 | 0.448 | 0.0265 | 0.0113 | 0.0004 | 0.9618 |
| 763F | Non-breeder | 5.3 | 0.8066 | 664 | 0.447 | 0.0261 | 0.0101 | 0.0005 | 0.9634 |
| 768F | Non-breeder | 0.9 | 0.7862 | 708 | 0.353 | 0.026 | 0.0113 | 0.0003 | 0.9624 |
| 769M | Non-breeder | 1.9 | 0.7708 | 642 | 0.345 | 0.0243 | 0.0113 | 0.0002 | 0.9641 |
| 772U | Non-breeder | 1.0 | 0.794 | 681 | 0.412 | 0.0245 | 0.0106 | 0.0005 | 0.9643 |
| 775M | Non-breeder | 1.7 | 0.7968 | 715 | 0.383 | 0.0258 | 0.0096 | 0.0002 | 0.9644 |
| 777M | Non-breeder | 2.3 | 0.8062 | 740 | 0.418 | 0.0255 | 0.0099 | 0.0003 | 0.9642 |
| 780M | Non-breeder | 0.8 | 0.8156 | 635 | 0.464 | 0.0229 | 0.0118 | 0.0005 | 0.9648 |
| 808F | Non-breeder | 0.4 | 0.8112 | 703 | 0.343 | 0.0264 | 0.0116 | 0.0005 | 0.9615 |
| 809F | Non-breeder | 2.4 | 0.8096 | 643 | 0.462 | 0.026 | 0.0105 | 0.0004 | 0.9631 |
| 810F | Non-breeder | 1.4 | 0.807 | 658 | 0.341 | 0.0278 | 0.0097 | 0.0006 | 0.9619 |
| 811F | Non-breeder | 1.5 | 0.8175 | 763 | 0.426 | 0.0241 | 0.0089 | 0.0005 | 0.9664 |
| 812M | Non-breeder | 2.5 | 0.7777 | 698 | 0.356 | 0.0258 | 0.0102 | 0.0003 | 0.9637 |
| 813F | Non-breeder | 0.7 | 0.783 | 654 | 0.385 | 0.0258 | 0.0112 | 0.0004 | 0.9626 |
| 818F | Non-breeder | 1.0 | 0.8299 | 684 | 0.495 | 0.0253 | 0.0101 | 0.0005 | 0.9641 |
| 819F | Non-breeder | 2.0 | 0.8143 | 600 | 0.475 | 0.0249 | 0.0106 | 0.0005 | 0.964 |
| 822F | Non-breeder | 3.3 | 0.8252 | 658 | 0.491 | 0.0255 | 0.0101 | 0.0007 | 0.9637 |
| 823F | Non-breeder | 1.6 | 0.7738 | 680 | 0.335 | 0.0258 | 0.0112 | 0.0003 | 0.9627 |
| 824M | Non-breeder | 1.5 | 0.7938 | 708 | 0.359 | 0.0246 | 0.0108 | 0.0007 | 0.9639 |
| 825F | Non-breeder | 1.7 | 0.8367 | 752 | 0.464 | 0.0257 | 0.0103 | 0.0004 | 0.9636 |
| 826F | Non-breeder | 1.1 | 0.8416 | 582 | 0.543 | 0.0259 | 0.01 | 0.0007 | 0.9634 |
| 827M | Non-breeder | 2.0 | 0.8149 | 585 | 0.434 | 0.0267 | 0.0104 | 0.0003 | 0.9626 |
| 828M | Non-breeder | 1.9 | 0.7992 | 583 | 0.411 | 0.0253 | 0.0109 | 0.0006 | 0.9633 |
| 829F | Non-breeder | 1.5 | 0.7989 | 620 | 0.429 | 0.0249 | 0.0099 | 0.0007 | 0.9645 |
| 830F | Non-breeder | 2.9 | 0.7829 | 679 | 0.376 | 0.0238 | 0.01 | 0.0004 | 0.9659 |
| 831F | Non-breeder | 2.1 | 0.7994 | 713 | 0.422 | 0.0249 | 0.0109 | 0.0006 | 0.9636 |
| 869M | Non-breeder | 2.0 | 0.8143 | 617 | 0.448 | 0.0254 | 0.0101 | 0.0005 | 0.964 |
| 872F | Non-breeder | 2.9 | 0.8116 | 600 | 0.438 | 0.0236 | 0.0117 | 0.0007 | 0.964 |
| 905M | Non-breeder | 3.6 | 0.8112 | 496 | 0.203 | 0.0277 | 0.0105 | 0.0001 | 0.9617 |
| 906M | Non-breeder | 1.8 | 0.7988 | 617 | 0.411 | 0.0263 | 0.0115 | 0.0005 | 0.9617 |
| 908F | Non-breeder | 0.8 | 0.8028 | 746 | 0.43 | 0.0262 | 0.0109 | 0.0004 | 0.9625 |
| 924F | Non-breeder | 0.8 | 0.7871 | 704 | 0.394 | 0.0251 | 0.011 | 0.0005 | 0.9635 |
| 961M | Non-breeder | 0.6 | 0.7726 | 638 | 0.381 | 0.0241 | 0.0114 | 0.0004 | 0.9642 |
| 964M | Non-breeder | 3.7 | 0.7923 | 674 | 0.415 | 0.0254 | 0.011 | 0.0005 | 0.9631 |
| 967M | Non-breeder | 1.5 | 0.7831 | 684 | 0.342 | 0.023 | 0.0112 | 0.0005 | 0.9653 |
| 968F | Non-breeder | 1.6 | 0.7825 | 650 | 0.392 | 0.026 | 0.01 | 0.0005 | 0.9635 |
| 977F | Non-breeder | 2.8 | 0.7963 | 649 | 0.429 | 0.0267 | 0.0099 | 0.0007 | 0.9627 |
| 979F | Non-breeder | 1.5 | 0.8007 | 647 | 0.308 | 0.0253 | 0.0115 | 0.0007 | 0.9625 |
| 993M | Non-breeder | 3.2 | 0.7955 | 707 | 0.433 | 0.0244 | 0.0114 | 0.0007 | 0.9635 |
| 994M | Non-breeder | 1.6 | 0.7715 | 655 | 0.369 | 0.0256 | 0.0115 | 0.0004 | 0.9625 |
| 995F | Non-breeder | 1.9 | 0.7891 | 677 | 0.404 | 0.0267 | 0.0104 | 0.0006 | 0.9623 |
| 996M | Non-breeder | 2.8 | 0.7978 | 684 | 0.424 | 0.0242 | 0.0114 | 0.0003 | 0.964 |
| 239M | Unknown |  | 0.7807 | 679 | 0.363 |  |  |  |  |
| 240M | Unknown |  | 0.8124 | 731 | 0.402 | 0.0256 | 0.0108 | 0.0006 | 0.963 |
| 285M | Unknown |  | 0.8076 | 693 | 0.397 | 0.0255 | 0.01 | 0.0004 | 0.9642 |
| 434F | Unknown | 3.3 | 0.7946 | 713 | 0.39 | 0.0259 | 0.0104 | 0.0004 | 0.9633 |

